# Microglial activation and alpha-synuclein oligomers drive the early inflammatory phase of Parkinson’s disease

**DOI:** 10.1101/2025.08.25.671921

**Authors:** James R. Evans, Melissa Grant-Peters, Joseph S. Beckwith, Christina E. Toomey, Rebeka Popovic, Jonathan C. Breiter, Jonathan W. Brenton, Aine Fairbrother-Browne, Stephanie Strohbuecker, Joanne Lachica, Maria Rodriguez-Lopez, Emma E. Brock, Bin Fu, Leila Nahidiazar, Patricia Lopez-Garcia, Ryan Ferguson, Rebecca S. Saleeb, Hannah Lucas-Clarke, Alexis Penverne, Karishma D’Sa, Chun Wei Pang, Mathew H. Horrocks, Michele Vendruscolo, Nicholas Wood, Steven F. Lee, Mina Ryten, Sonia Gandhi

## Abstract

Parkinson’s disease (PD) is characterised by insoluble α-synuclein (αSyn) aggregates in Lewy bodies (LBs) within the substantia nigra, with cortical pathology appearing as the disease progresses. Late-stage LB deposition, cellular stress, and neuronal loss obscure disease-driving events, we therefore performed multi-regional transcriptomic and aggregate profiling in early-midstage PD brains (Braak 3–4), where cortical regions are pathologically unaffected. We report neuroimmune activation as an early PD feature, characterised by the expansion of a high-*SNCA*-expressing microglial state. This robust immune signature occurs prior to LB formation, but is associated with oligomeric αSyn within cortical microglia. In hiPSC-derived microglia, both endogenous αSyn oligomerisation, and exogenous oligomer uptake, trigger transcriptional reprogramming, characterised by interferon-driven inflammation, antigen presentation, and mitochondrial suppression, closely mirroring the early PD brain. These findings describe mechanisms by which αSyn oligomerisation potently initiates early neuroinflammation, highlighting a critical interplay between proteinopathy and immune activation at the earliest stages of disease.

## INTRODUCTION

Parkinson’s disease (PD) is a progressive neurodegenerative disorder characterised by the misfolding and aggregation of alpha-synuclein (αSyn) into neuronal inclusions known as Lewy bodies (LBs).^1,2^ Pathology advances in a stereotyped anatomical sequence, emerging first in selected brainstem nuclei and subsequently in cortical regions at later stages. This progression is classified neuropathologically into six Braak stages, with stages 5–6 representing end-stage disease marked by widespread cortical involvement, extensive LB deposition, and profound dopaminergic neuronal loss.^1,3^ Most studies of human PD have examined late pathological stages (Braak 5–6), where LB pathology and neuronal loss dominate the molecular landscape, obscuring early pathogenic events that drive disease onset and progression. Although these analyses have identified alterations in synaptic transmission, mitochondrial function, proteostasis, and inflammation,^4–6^ such signatures at this stage of disease likely reflect secondary consequences of advanced neurodegeneration.

Dopaminergic neuron loss in the substantia nigra, driven by neuronal αSyn aggregation, and the resulting dopaminergic deficit, underlies the motor syndrome.^1^ Glial pathology, such as microglial and astrocytic activation, may occur as a consequence of protein misfolding and neuronal dysfunction, and thus contribute to PD pathogenesis.^7,8^ However, genome-wide association studies link PD risk loci to genes enriched in glial cell types, in addition to dopaminergic neurons,^9,10^ whilst experimental models show that glia can modulate αSyn aggregation, promote its intercellular transmission, and influence neuronal survival.^7,8,11–13^ Together, these findings raise the hypothesis that glial biology may be a primary, or causal event, in PD pathogenesis.

Whilst LBs are the defining pathological hallmark of PD, and form the basis of post-mortem diagnosis, their toxicity is unclear.^14^ They represent an end-stage phenomenon, found in surviving neurons after prolonged cellular stress and degeneration, and may play a protective role by sequestering otherwise more toxic species.^15^ The earliest aggregation events occurring prior to LB formation, including the first appearance of αSyn assemblies, and their distribution across cell types remain poorly defined. αSyn aggregation occurs through the self-assembly of monomers into small soluble protein assemblies, which gradually increase in b-sheet structure, ultimately forming insoluble fibrils and LBs.^16^ Although the identity of the primary pathogenic αSyn species remains debated, *in vitro* and *in vivo* studies indicate that specific oligomeric conformers are highly cytotoxic.^15^ These assemblies permeabilise membranes,^17^ impair mitochondrial function and induce oxidative stress,^18,19^ disrupt calcium dynamics^20,21^ and synaptic transmission,^21,22^ and elicit inflammatory responses in astrocytes.^21,23^ A major challenge has been the detection of these oligomeric species, or nanoscale assemblies, in tissue in vivo - we recently developed ASA-PD,^24^ a sensitive imaging method for visualising αSyn oligomers in post-mortem brain, and this approach has enabled the in vivo characterisation of early aggregation events in relation to specific cell types and stages of pathology.

To identify pathways early in the disease course, postmortem brain cohorts consisting of early/mid-stage PD (Braak 3–4) are particularly valuable – as they have restricted distribution of neuronal LB pathology to the brainstem and striatum, with minimal or no cortical involvement. This stage offers a unique opportunity to investigate pathogenic processes before cortical pathology commences, enabling a cortical prodrome to be captured. This approach, which we have previously applied in proteomic studies of the PD brain,^25^ distinguishes early disease mechanisms from downstream consequences and provides insights into the molecular basis of PD preceding clinical (cortical) symptom onset. Our study investigates the relationship between biological processes and aggregation in mid-stage PD across multiple brain regions. Our cohort included 18 PD brains and 21 control brains, sampling 8 brain regions across a range of LB pathological loads, from high (e.g. substantia nigra) to low (e.g. parietal cortex). We generated transcriptomic data to study biological processes associated with disease, including bulk RNA sequencing and paired multiome snRNAseq and snATACseq. This provided insight into disease-associated cell states, gene expression and pathway changes. We performed LB and cell-type specific oligomer mapping on the same samples. Our findings were then validated *in vitro* using human iPSC-derived models. We identify an early inflammatory cellular phase of PD that occurs in the unaffected cortical regions, prior to LB formation, and describe the mechanisms underlying this inflammatory phase.

## RESULTS

### Early immune activation in cortical regions precedes Lewy Body pathology

Our sex and age-matched sample cohort consisted of post-mortem brain tissue from 18 individuals with mid-stage PD (without dementia) and 20 healthy controls (Figure S1A-B; see Table S1; Data S1). Samples underwent detailed neuropathological assessment consisting of αSyn staging using the Braak and McKeith criteria and were classified as Braak stages 3 (N = 5) and 4 (N = 13). It is established that LB pathology in mid-stage disease follows a gradient of severity, therefore we performed bulk RNA sequencing on samples from 8 brain regions (234 samples in total) separating them by severe (substantia nigra), moderate (caudate and putamen), mild (anterior cingulate cortex, temporal cortex and parahippocampal gyrus) and absent (frontal cortex and parietal cortex) LB pathology.

Differential gene expression between cases and controls was compared within each brain region as well as modelled across all brain regions (pan-regional), all cortical regions and all sub-cortical regions. We identified 1,415 differentially expressed genes (DEGs) pan-regionally and within individual regions, with a mean of 407 (standard deviation = 184) DEGs identified. All analyses demonstrated a higher proportion of upregulated versus downregulated DEGs. This pattern of expression was most evident in cortical regions (up/down = 2.67) as compared to subcortical (up/down = 1.47) brain regions (Data S2 for DEGs). This finding differs from previous analyses of late-stage PD which have demonstrated predominant downregulation of gene expression at a bulk and single-cell level.^26,27^

Across all brain regions, except for the caudate, we also identified a higher number of significant pathway enrichments amongst upregulated versus downregulated genes, with the highest proportion of upregulated pathways detected in cortical as compared to subcortical brain regions (cortical pathway up/down = 62, sub-cortical pathway up/down = 25). Significantly downregulated pathways included those involved in amino acid metabolism and mitochondrial function. Significantly upregulated pathways related primarily to immune function. Key pathways involved in innate and adaptive immune activation such as toll-like receptor signalling (FDR-corrected p = 1.24×10^-11^), interleukin-1 and interleukin-6 production (FDR-corrected p = 1.95×10^-15^; FDR-corrected p = 3.15×10^-27^), MHC class II antigen presentation (FDR-corrected p = 5.63×10^-8^) and neutrophil degranulation (FDR-corrected p = 1.34×10^-6^) were upregulated across all cortical regions (Figure 2A-B, Figure S1D). Cell type-specific analysis of upregulated DEGs demonstrated significant enrichment for microglia-associated genes (Figure 2C, All brain regions, FDR-corrected p < 0.001). Vascular cell–associated genes were also enriched, although to a lesser extent (Figure 2C, All brain regions, FDR-corrected p < 0.001).

**Figure 1.**
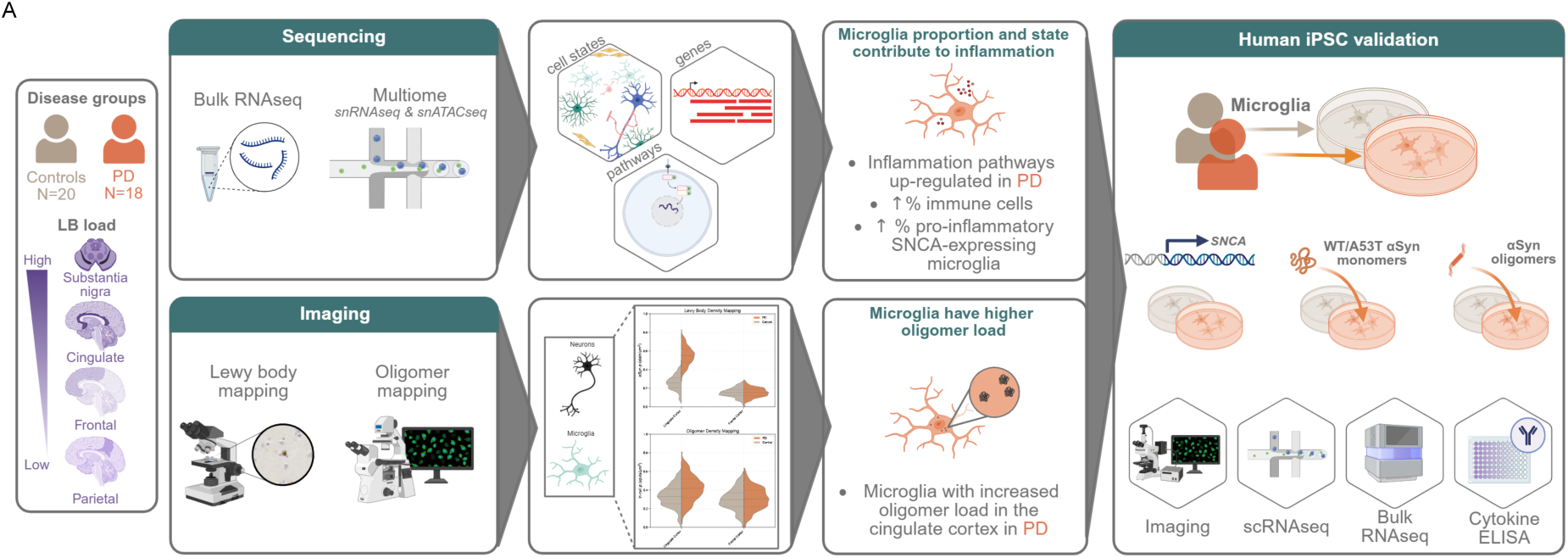
Schematic illustration of experimental paradigms across post-mortem brain and human iPSC-derived models.

**Figure 2.**
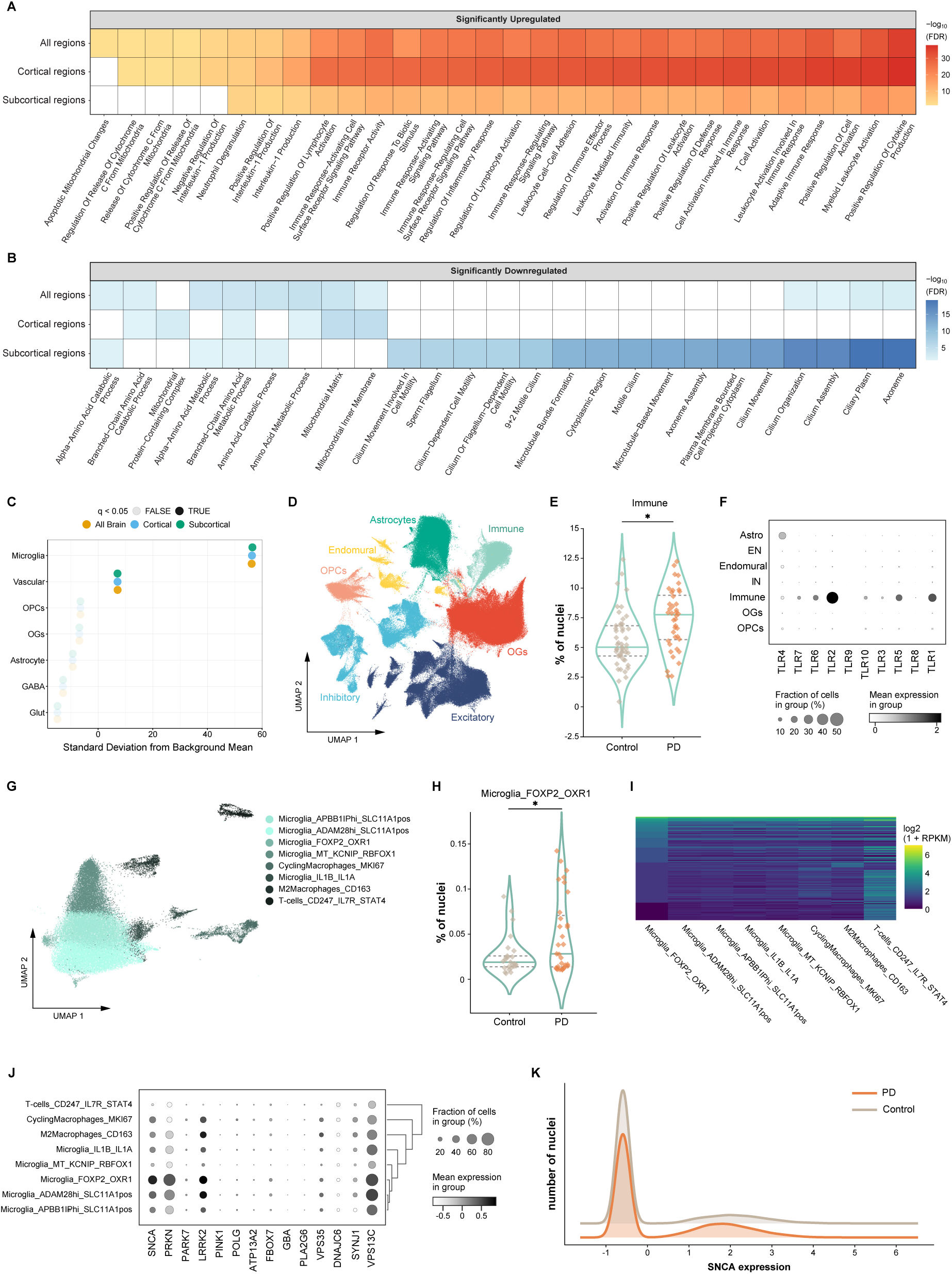
Transcriptomic analysis of mid-stage Parkinson’s disease across brain regions. (A) Gene Ontology pathway enrichment of upregulated differentially expressed genes detected in bulk RNA sequencing data. (B) Gene Ontology pathway enrichment of downregulated differentially expressed genes detected in bulk RNA sequencing data. (C) Expression-weighted cell enrichment analysis of significantly upregulated genes across all brain regions in bulk RNA sequencing. Upregulated genes were significantly enriched in microglial and vascular cell populations. (D) Paired snRNAseq and snATACseq included 955,571 nuclei, of which 740,157 passed QC. Following data-driven clustering, Leiden community detection and annotation, we identified 7 major cell types. (E) Cell type proportion analysis of snRNAseq data revealed that the immune cell compartment, which includes microglia and peripheral immune cells, was significantly more represented in PD than in controls (FDR-corrected p value = 1.63 x 10^-2^). (F) The immune compartment identified by snRNAseq analysis had the highest expression of TLR genes, which are key players in the Toll-like receptor activity observed in the bulk RNAseq analysis. (G) UMAP showing the 8 cell types/states identified within the immune cell compartment. This included T-cells, cycling macrophages, M2 macrophages as well as 5 states of microglia. (H) Microglia marked by expression of FOXP2 and OXR1 were proportionally more represented in PD when compared to controls (FDR-adjusted p value = 1.77 ×10^-2^). (I) Microglia_FOXP2_OXR1 have a distinct ATAC peak profile when compared to other cells in the immune compartment. (J) Dotplot showing across immune cell states the expression of PD causal genes, such as *SNCA*, *PRKN* and *LRRK2*. (K) Density plot showing the normalised expression of SNCA in microglia in controls and PD. A greater proportion of microglia express SNCA in PD than in controls - in controls, 23.1% of nuclei express SNCA (expression >0), while this is 35.5% of microglia in PD.

Given the enrichment for microglia-associated genes, we compared our DEGs with established disease-associated microglial signatures.^28–30^ Our DEGs showed significant overlap with a number of disease-associated microglial (DAM) states, patterns that persisted in cortical-only analyses (Figure S2).

Together, these gene expression changes evidence widespread alterations in the early-mid PD brain and critically identify a strong microglia-associated inflammatory response in cortical areas.

### Microglia are critical mediators of innate and adaptive immunity in early Parkinson’s disease

To dissect the biological mechanisms and cell types underlying the inflammatory signal within cortical regions we performed single-nuclear RNA (snRNAseq) and ATAC sequencing on samples from the parietal, frontal and anterior cingulate cortex. We sequenced a total of 955,571 nuclei, of which 740,157 passed quality control for RNAseq metrics and formed the basis of this analysis. Following batch correction, dimensionality reduction and Leiden community detection, canonical markers were used to assign clusters to major cell types, namely excitatory neurons, inhibitory neurons, astrocytes, immune, oligodendrocytes (OGs), oligodendrocyte progenitor cells (OPCs), and a mixed compartment containing mural and endothelial cells termed endomural (Figure 2D). Given the innate immune transcriptional signature in the cortex, we determined whether there were accompanying changes in cell state. We thus further stratified cell types to identify: 8 immune, 19 excitatory neuron, 17 inhibitory neuron, 10 endomural, 13 astrocyte, 9 oligodendrocyte and 2 OPC cell states (Figure S3-5).

To explain the robust innate immune responses identified in the bulk RNA-seq data, we began by investigating changes in cell type proportion. Pooled cell abundance across all three surveyed cortical brain regions revealed a significantly higher proportion of immune cells in PD when compared to controls (Figure 2E, Figure S6-7, FDR-corrected p value = 1.63×10-2). Given that immune cells express the highest level of TLRs among brain cells, it is likely that this change in immune cell proportion is sufficient to explain the increased TLR expression in bulk RNA sequencing data (Figure 2F). To investigate this further, we leveraged the high numbers of nuclei captured in each sample to investigate microglial states in disease, since microglia are the most abundant of all immune cell types in the brain (Med = 88.8%) (Figure 2G-I). We found that in mid stage PD there was a higher proportion of microglia of a specific state, namely Microglia_FOXP2_OXR1 (Figure 2H, FDR-adjusted p value = 1.77×10-2), characterised by a distinct open chromatin profile compared to other immune cells, as demonstrated by the ATAC sequencing data (Figure 2I). This cell state also had upregulated lymphocyte, chemokine and ribosomal pathways (Figure S8A-B) and had the highest expression levels of TLR genes (Figure S8C). This suggested that the PD-associated changes were not exclusively due to a higher proportion of total immune cells, but that there was a shift in the proportion of microglia across cellular states.

We next investigated whether cells in this state might be associated with other pathological processes, such that overrepresentation of this population might impact pathology as well. Indeed, this cell state had particularly high expression levels of αSyn encoding gene *SNCA*, as well as other genes causally implicated in PD such as *PRKN* and *LRRK2* (Figure 2J). This indicates that in the context of PD, microglia might have a role in increasing overall *SNCA* expression within the brain tissue by the presence of a higher proportion of microglia expressing *SNCA*. We found that in controls 23.1% of nuclei expressed *SNCA*, whilst in PD this proportion was 35.5% of microglia (Figure 2K).

In addition to microglia, we identified clusters among other cell types which were associated with inflammatory markers. As might have been expected, these included astrocyte and oligodendrocyte clusters, which have increasingly been implicated in PD-related inflammation.^5,7,8,23^ Interestingly, we also identified excitatory neuronal cell states characterised by the expression of genes implicated in inflammation. One of these subtypes, RORB_MET_IL1RAPL2, was marked by high expression levels of interleukin receptors (*IL1RAPL2*, *IL1RAP*, *IL12RB2*, *IL1RL2* and *IL6ST*) and was proportionally more represented in the parietal cortex in PD (Figure S8C-D, FDR-adjusted p value = 3.70×10^-3^). Overall, this indicates that all cell types may participate in contributing to the immune environment in early PD.

We postulated that the significant shifts in cell state, as captured through analyses of cell type proportion, would also be associated with significant cell type-specific differential gene expression. While we found widespread gene expression changes in all major cell types and in all cortical brain regions (Figure 3A), we detected the highest numbers of DEGs amongst subtypes of excitatory neurons and oligodendrocytes. To explore the biological processes underlying these changes in gene expression, we performed gene ontology and pathway enrichment analysis (Figure 3B). This analysis revealed that across all brain regions, pathway enrichments were most seen amongst DEGs detected in excitatory neuronal clusters, and these often involved voltage-gated channel activity. However, we also noted significant pathway enrichments detected in single cortical regions. In the frontal cortex, 60% of the pathways disrupted in 3 or more cell types related to translation (translation regulator activity, translation initiation factor activity). In the cingulate cortex, we found significant enrichments across a range of immune processes, including elements of both the adaptive and innate immune response, specifically amongst DEGs detected in CUX2 and RORB neuronal clusters.

**Figure 3.**
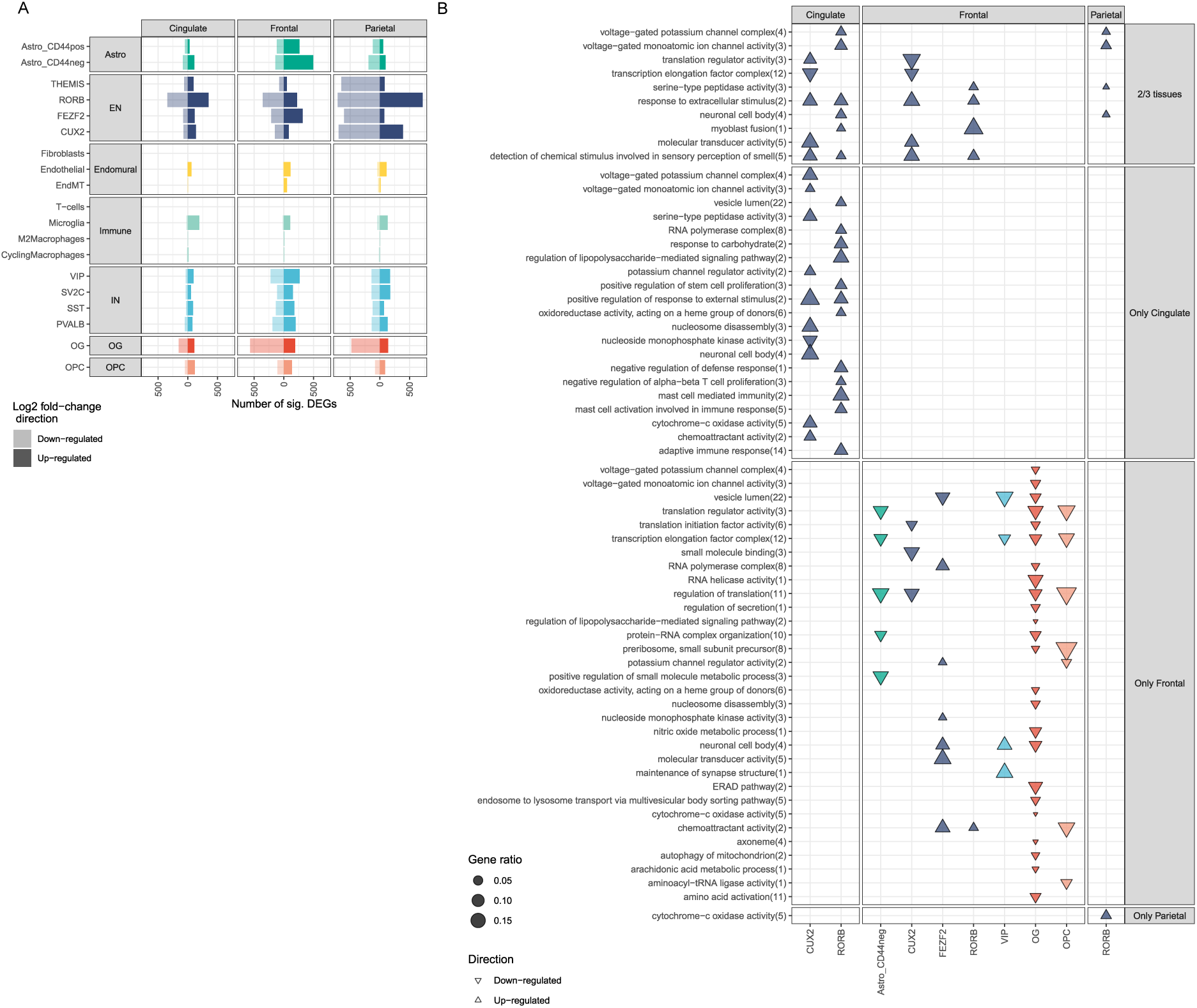
Transcriptomic analysis of mid-stage Parkinson’s disease across brain regions. (A) Differential gene expression analysis results for snRNAseq data, results shown for each brain region and cell type. (B) Pathway enrichment analysis of differentially expressed genes showed that neurons had the most pathway changes in PD across all brain regions. In the cingulate cortex, this included several immune associated pathways such as adaptive immune response (14 pathways), mast cell activation involved in immune response (5 pathways), mast cell mediated immunity (2 pathways), negative regulation of alpha-beta T cell proliferation (3 pathways).

We further characterised cell-specific DEGs by investigating the enrichment of genes causally-implicated in neurodegeneration. Given that pathological studies have demonstrated that co-pathology is a common feature of PD with amyloid plaques and tau tangles detected in a significant proportion of brain samples derived from PD patients, we assessed genes causally-implicated in AD as well as PD. While we found that the PD genes were not enriched in any cell type in any of the three tested regions (Figure S8E), genes causally implicated in AD were significantly enriched in microglia of the cingulate cortex (FDR-adjusted p value = 4.12 x 10^-2^). Thus, these results highlighted a key role for microglia in inflammatory processes in mid-stage PD.

### An early cellular phase is associated with αSyn oligomers, rather than large aggregate pathology

Inflammation in PD is prominent in the midbrain at stages where neuronal loss and LB pathology are extensive.^5,31^ Despite the expected lack of widespread LB deposition in cortical regions in mid-stage PD, we observed a strong and widespread transcriptional inflammatory signature in these regions. We therefore investigated how this related to the extent of αSyn aggregation.

First, to characterise the extent of LB pathology across our cohort, we performed αSyn immunohistochemical staining in all cases and regions and quantified the percentage of αSyn-positive area relative to total tissue area (‘αSyn positivity’). In accordance with Braak 3–4 staging, we found significant differences in the amount of αSyn pathology across the studied brain regions (Figure 4A, Figure S9, mixed effects model p = 0.0047) with the highest levels in the putamen (0.084% mean positivity), caudate (0.032%) and parahippocampus (0.041%), followed by the cingulate cortex (0.024%). Pathology was very minimal in cortical regions (temporal cortex 0.009%, frontal cortex 0.004%, parietal cortex 0.003%). This quantification confirms that in cortical regions, where inflammation is observed, there is minimal LB pathology.

**Figure 4.**
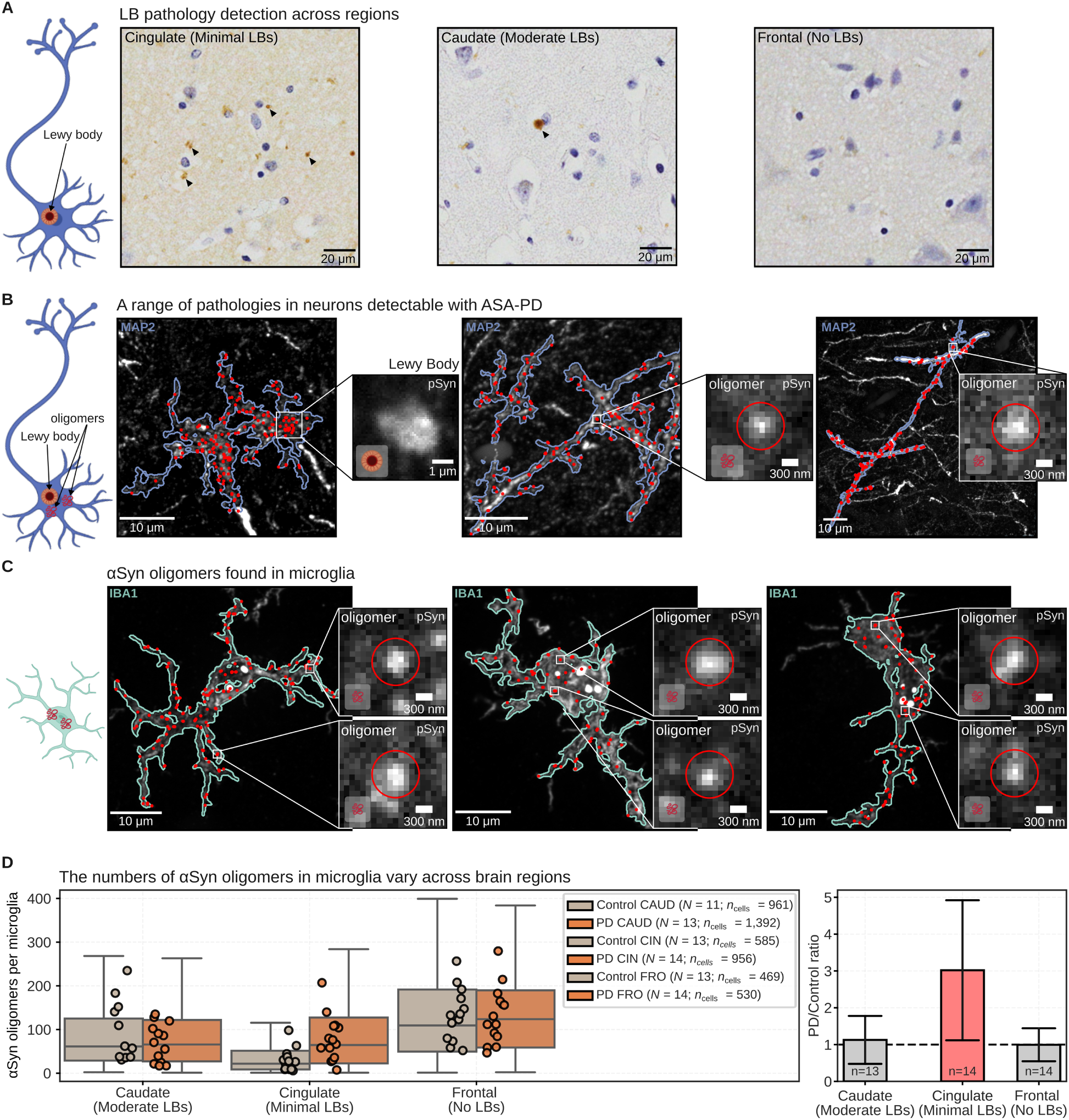
DAB and single-oligomer imaging show a range of pathology in neurons and microglia. (A) Exemplar images across brain regions DAB-stained for αSyn, highlighting differing pathology dependent on brain region. Sections were counterstained with Haematoxylin (blue). Arrows indicate LB pathology. Scale bars represent 20µm. (B) Fluorescence images of larger aggregates and single oligomers (red puncta) in neurons (blue outline, MAP2 staining) detected by ASA-PD. (C) ASA-PD also detects single oligomers (red puncta) of αSyn inside microglial cells (green outline, IBA1 staining). (D) Analysis of the number of oligomers inside microglia, showing that the number of oligomers inside microglia (assuming a microglia to be a sphere of 5 μm radius) depends on brain region (left), with a strong difference (summary bar plot, right) observed in the cingulate cortex between PD and control.

Prior to LB formation, smaller assemblies of αSyn, termed oligomers, can form. We therefore examined whether cortical regions in Braak stage 3–4 brains showed evidence of these earliest stages of protein aggregation. Our previously developed ASA-PD pipeline^24^ enables the detection of both small oligomeric and larger aggregate (LB type) species of αSyn (Figure S10), and to determine their spatial relation to cell types (Figure S11). We applied the ASA-PD pipeline to the mid-stage samples to investigate the relationship between the abundance of αSyn oligomers and the inflammatory state.

Consistent with our DAB imaging, ASA-PD revealed oligomeric αSyn (phosphorylated at Serine 129; pSyn) within neurons (Figure 4B; Figure S12A) in both control and PD brain, where evidence suggests oligomerisation is necessary for its physiological function.^32–34^ In LB-free regions, there was no significant difference in the brightness (size) of the neuronal oligomers in PD compared to controls (Figure S12B). Notably, oligomeric αSyn was also observed in microglia across all cases (Figure 4C). In the cingulate cortex, microglia contained significantly more αSyn oligomers in PD than in controls (Figure 4D).

Oligomer accumulation may result from increased intracellular αSyn monomer concentration, as in SNCA gene dosage models, or from uptake of aggregates from the extracellular space. The higher oligomer load within microglia in the cingulate cortex could therefore reflect the increased proportion of high-*SNCA*-expressing FOXP2_OXR1 microglia we identified in our transcriptomic analyses, or active phagocytosis of neuronally derived oligomers.

Given the increased microglial αSyn oligomer density in the cingulate cortex we applied spatial transcriptomics (CosMx Human 6K Discovery panel) to this region to investigate changes in microglial localisation. Four Braak stage 3–4 PD cases and four matched controls were profiled (Figure S13A). Cells were annotated to the seven major cell types defined in our snRNAseq transcriptomic analysis (Figure S13B-C). Louvain clustering of immune cells based on their extended spatial neighbourhoods revealed three distinct immune neighbourhood clusters (INC). INC comprised immune cells neighbouring other immune cells, neurons and astrocytes; INC2 comprised immune cells adjacent to oligodendrocytes; and INC3 comprised immune cells predominantly neighbouring endomural cells (Figure S13D-F).

Since activated microglia cluster around sites of injury,^35^ and around amyloid-β plaques^29,36–38^, we examined immune-immune neighbourhood interactions. These were most frequent in INC1, where *COX1* and *COX2*, enzymes in prostaglandin biosynthesis,^39^ and the AD and PD risk factor genes *APOE*^40^ *and CTSB*^41,42^ were enriched (Figure S13G). Stratification of immune–immune interactions into high (>5 immune cell neighbours; n = 410) and low (<5 immune cell neighbours; n = 2,037), revealed a shift toward an increased frequency of high immune–immune interactions in PD (Figure S13H). Differential gene expression analysis of INC1 between PD and controls, followed by GSEA of genes ranked by log_2_FC, revealed receptor clustering as significantly upregulated in PD (Figure S13I). This pathway included increased expression of *APOE*, associated with cognitive decline and mortality in PD^43,44^ as well as AD risk,^40^ and previously reported to be elevated in microglia from AD models and patient brain tissue.^28,29,40,45^ These findings indicate that the microenvironments in which immune cells, including microglia, reside, influence their inflammatory profiles, supporting a role for neighbouring cells in shaping microglial identity and signalling.

### hiPSC-derived microglia-like cells exhibit heterogeneous transcriptional states in vitro

Our brain snRNAseq identified an increased proportion of immune cells in PD, along with a shift toward a greater fraction of microglia expressing *SNCA.* Notably, this shift included an increased proportion of microglial subtypes such as FOXP2_OXR1, characterised by high *SNCA* expression. This may suggest that dysregulated endogenous *SNCA* in microglia, and not just αSyn uptake, could contribute to the increased intra-microglial oligomer load detected by ASA-PD. To model this *in vitro,* we generated hiPSC-derived microglia-like cells (iMGL) from healthy controls and PD patients carrying the *SNCA* A53T mutation, using an adapted published protocol (Figure 5A).^46^ The A53T mutation is causative of familial early-onset PD,^47^ and alters αSyn aggregation kinetics, promoting the formation of cytotoxic oligomers.^48,49^ We have previously demonstrated PD-associated pathology in *SNCA* A53T patient-derived midbrain^50^ and cortical neuron^18^ models.

**Figure 5.**
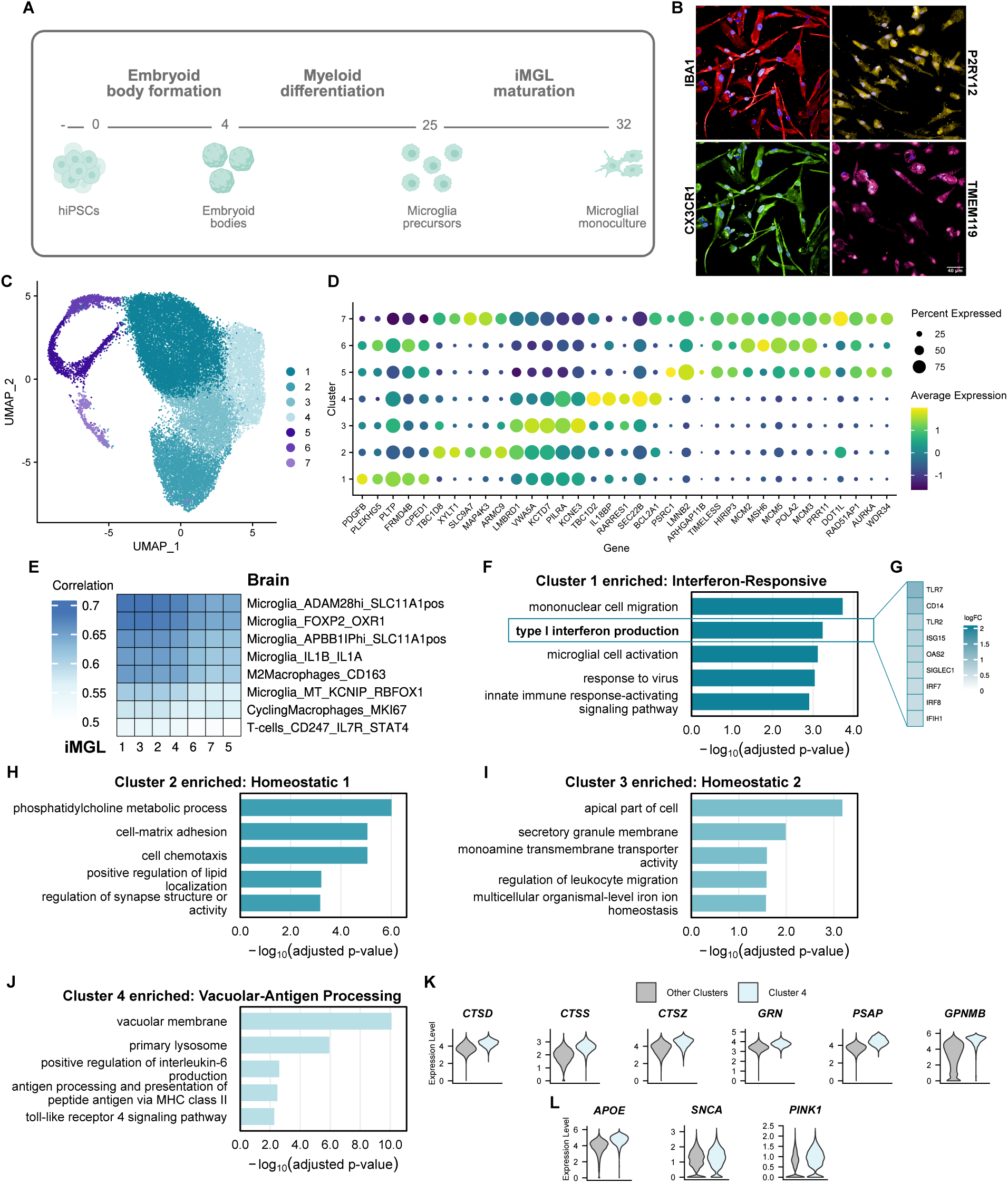
Transcriptionally defined iMGL clusters reveal functional specialisation. (A) Schematic of differentiation protocol to generate iMGL model from hiPSCs. (B) Representative immunocytochemistry images demonstrating that iMGLs express the microglia markers IBA1, CX3CR1, P2RY12, and TMEM119. (C) Uniform Manifold Approximation and Projection (UMAP) plot showing the clustering of the integrated dataset using the cells across all the samples. Non-proliferative iMGL clusters in cyan and proliferative iMGL clusters in purple. (D) Dot plot of top enriched genes (n = 5 per cluster) defining each iMGL cluster. (E) Correlation of iMGL clusters with immune subtypes from the brain snRNA-seq. (F) Selected unique Gene Ontology terms associated with the genes significantly enriched in Cluster 1. (G) Upregulated genes driving type I interferon signalling enrichment in Cluster 1. (H) Selected unique Gene Ontology terms associated with the genes significantly enriched in Cluster 2. (I) Selected unique Gene Ontology terms associated with the genes significantly enriched in Cluster 3. (J) Selected unique Gene Ontology terms associated with the genes significantly enriched in Cluster 4. (K) Violin plots showing significant enrichment of cathepsins (*CTSD*, *CTSS*, *CTSZ*) and lysosomal genes (*PSAP*, *GRN*, *GPNMB*) in Cluster 4 versus all other clusters. (L) Violin plots showing significant enrichment of *APOE*, *SNCA*, and *PINK1* in Cluster 4 versus all other clusters.

iMGL cultures were highly enriched, expressing canonical microglia markers (IBA1, CX3CR1, P2RY12, and TMEM119) by immunocytochemistry (Figure 5B). They exhibited robust and typical functional properties, including phagocytosis of pHrodo™ Zymosan particles (Figure S14), and a strong transcriptional response to LPS (10 ng/ml), with significant upregulation of pathways related to bacterial sensing, chemotaxis, and innate immune regulation, alongside downregulation of mitochondrial pathways such as oxidative phosphorylation and aerobic respiration across both control and A53T genotypes (Figure S15A-B). LPS stimulation also significantly suppressed *TREM2* expression, consistent with previous microglia models,^51^ and downregulated homeostatic microglial genes including *PSRY12*, *HEXB*, and *GPR34* (Figure S15C).

Human microglia, as shown previously, display heterogenous transcriptional states.^30^ Importantly, this transcriptional diversity can be recapitulated *in vitro* using human iMGL models.^52^ Given that shifts in state drive the inflammatory phenotype of early PD brain, we performed single-cell RNA sequencing (scRNAseq) on iMGL across control (n = 3) and *SNCA* A53T (n = 3) donors. Across the integrated dataset of 30,009 high-quality cells, we identified 7 distinct clusters (Figure 5C), all of which were represented across both genotypes (Figure S16A-B). As previously reported, iMGL models generate both non-proliferative and proliferative clusters,^52^ with clusters 5-7 identified as proliferative based on cell cycle scoring and expression of proliferation-associated genes (Figure S16C, Figure 5D). The microglial markers *AIF1* (IBA1), *TREM2*, *TYROBP*, and *C1QA* were found to be expressed across all clusters (Figure S16D), whilst markers indicative of astrocytes (*GFAP*), oligodendrocytes (*SOX10*), and neurons (*MAP2*) were absent from the culture in nearly all cells (Figure S16E). We next performed correlation analysis of our *in* vitro iMGL model with the immune clusters defined in our snRNAseq data from post-mortem brain. This revealed that all iMGL clusters had the highest similarity to microglia subtypes, notably ADAM28_SLC11A1 and FOXP2_OXR1, and the lowest similarity to cycling macrophages or T cells (Figure 5E), confirming the relevance of this model for studying microglia phenotypes in the context of PD.

To further characterise the iMGL clusters, we performed functional enrichment analysis of genes significantly upregulated in each cluster, to reveal distinct, cluster-specific pathways. Cluster 1 was uniquely enriched for type I interferon signalling (FDR-corrected p = 5.84×10^-4^) and antiviral responses such as response to virus (FDR-corrected p = 9.15×10^-4^; Figure 5F), with marked upregulation of toll-like receptors (*TLR2*, *TLR7*) and associated transcriptional regulators including *IRF7* and *IRF8*, as well as downstream interferon-stimulated genes such as *OAS2* (Figure 5G). Clusters 2 and 3 largely reflected homeostatic microglial functions, with Cluster 2 enriched for cell chemotaxis (FDR-corrected p = 8.91×10^-6^) and regulation of synapse structure or activity (FDR-corrected p = 6.66×10^-4^), and Cluster 3 enriched for monoamine transmembrane transporter activity (FDR-corrected p = 2.56×10^-2^) and iron ion homeostasis (FDR-corrected p = 2.69×10^-2^; Figures 5H–I). Cluster 4 was specifically enriched for vacuolar pathways such as vacuolar membrane (FDR-corrected p = 2.73×10^-13^) and antigen processing pathways such as antigen processing and presentation of peptide antigen via MHC class II (FDR-corrected p = 3.34×10^-3^; Figure 5J), and showed significant upregulation of cathepsins (*CTSD*, *CTSS*, *CTSZ*), which mediate the vacuolar route of antigen presentation,^53–55^ and have been implicated in PD and other neurodegenerative conditions.^56^ This was accompanied by increased expression of other lysosomal genes linked to PD pathogenesis, including *GRN*, *PSAP*, and *GPNMB* (Figure 5K).^41,57–59^ Cluster 4 also displayed significantly elevated levels of AD risk gene *APOE*^40^, as well as PD genes *SNCA* and *PINK1* (Figure 5L).

### Inflammatory profile of SNCA-mutant iMGL mirrors PD brain signatures

Having established the transcriptional landscape of iMGL, we next utilised our scRNAseq data to determine whether endogenous *SNCA* dysregulation within microglia drives a disease-relevant inflammatory phenotype (Figure 6A–B). We performed differential gene expression analysis on pseudobulked data for each cluster, followed by ranked GSEA, using log_2_FC effect size, to identify pathways altered in *SNCA* A53T versus control iMGL. GSEA revealed broad activation of inflammatory pathways across all clusters, characterised by antiviral responses and type I interferon signalling (Figure 6C). Notably, the high-*SNCA*-expressing Cluster 4 exhibited the strongest inflammatory signature, with response to virus (FDR-corrected p = 4.47×10^-8^), defence response to virus (FDR-corrected p = 9.30×10^-7^), and negative regulation of viral genome replication (FDR-corrected p = 7.22×10^-6^) the three most significant activated pathways (Figure 6C; Data S11). Examination of the genes driving the response to virus (GO:0009615) enrichment (Figure 6D) revealed upregulation of interferon-induced proteins (*IFI6*, *IFI44L*, *IFI44*, *IFI27*, *IFIT1*, *IFIT2*) and the interferon regulatory factor *IRF7*, recently identified to drive microglial inflammation in a PD model.^60^ Interestingly, Cluster 4 also showed marked upregulation of *HMGA2*, a known driver of inflammation through NF-κB signalling,^61^ and the highest upregulation of the interferon-inducible protein *RSAD2*/viperin, which has well-described mitochondrial consequences, including reduced cellular ATP and downregulation of mitochondria-encoded genes.^62,63^

**Figure 6.**
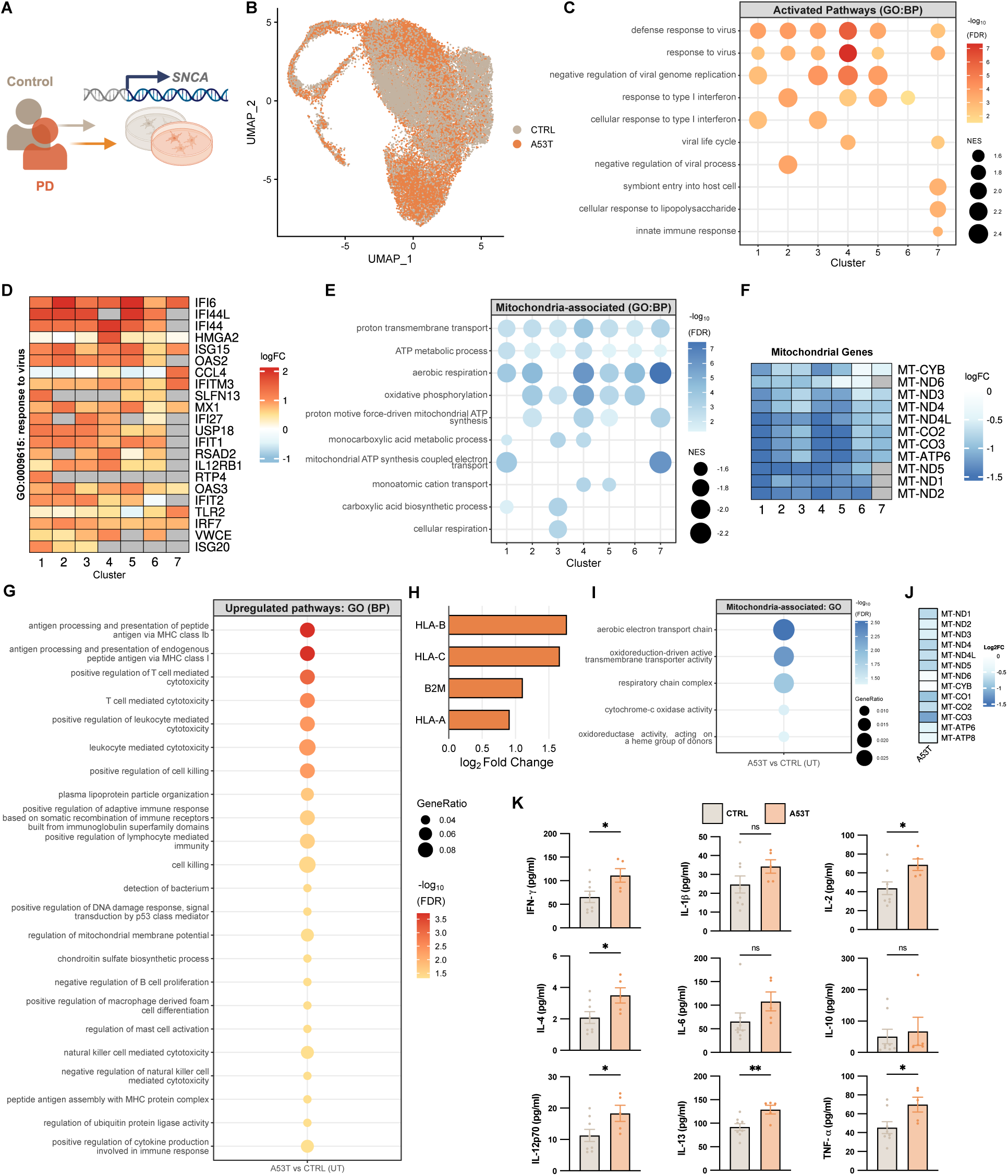
Endogenous *SNCA* dysregulation in iMGL induces inflammatory phenotypes with mitochondrial suppression. (A) Schematic of experimental design of iMGL from control and *SNCA* A53T donors. (B) Uniform Manifold Approximation and Projection (UMAP) plot showing the clustering of the integrated dataset in the control and *SNCA* A53T genotypes. (C) GSEA pathway analysis of upregulated pathways in *SNCA* A53T iMGL clusters compared to control from scRNAseq, highlighting antiviral and type I interferon responses. (D) Heatmap of Genes underpinning the response to virus pathway (GO:0009615) across all clusters (log₂FC > 1 in ≥ 1 cluster). (E) GSEA pathway analysis of Mitochondria-associated GO:BP downregulated pathways in *SNCA* A53T iMGL clusters compared to control from scRNAseq, highlighting mitochondrial suppression. (F) Heatmap of mitochondrial-encoded genes, showing suppression in *SNCA* A53T iMGL compared to control. (G) Gene Ontology (BP) of the most significant upregulated pathways in *SNCA* A53T compared to control iMGL from bulk RNA sequencing. (H) Log2FC for significantly upregulated MHC-1 genes (*HLA-A*, *HLA-B*, *HLA-C*, *B2M*) in *SNCA* A53T iMGL compared to control. (I) Gene Ontology (BP) of mitochondria-associated Gene Ontology terms significantly downregulated in *SNCA* A53T compared to control from bulk RNA sequencing. (J) Log2FC for mitochondrial-encoded genes in *SNCA* A53T iMGL compared to control. (K) Conditioned media from control (n = 8 iPSC lines) and *SNCA* A53T (n = 5 iPSC lines) iMGL was measured for a panel of cytokines by ELISA (data represent mean ± SEM; unpaired t-tests; *p < 0.05, **p < 0.01).

Consistent with these known effects of *RSAD2*/viperin, and more broadly the impact of interferon on mitochondrial function and mitochondria-encoded RNA expression,^64,65^ we found that pathways including oxidative phosphorylation, proton transmembrane transport, and ATP synthesis were significantly suppressed across clusters in the *SNCA* A53T iMGL (Figure 6E; Data S11). This was underpinned by marked downregulation of mitochondrial-encoded transcripts, with genes encoding subunits of NADH dehydrogenase (Complex I) showing the greatest decrease. Suppression was most pronounced in Cluster 1 (interferon-responsive) and Cluster 4 (vacuole–antigen processing).

To understand the genes and pathways dysregulated in *SNCA* A53T iMGL in further detail, we analysed bulk RNA sequencing from the same iMGL cultures. We performed differential gene expression analysis followed by functional enrichment (Gene Ontology) of the significantly upregulated and downregulated genes. This confirmed upregulation of immune pathways in *SNCA* A53T iMGL compared to controls, including antigen processing and presentation of peptide antigen via MHC class Ib (FDR-corrected p = 1.93×10^-4^), antigen processing and presentation of endogenous peptide antigen via MHC class I (FDR-corrected p = 1.97×10^-4^) and positive regulation of T cell mediated cytotoxicity (FDR-corrected p = 8.74×10^-4^) as the most significantly upregulated (Figure 6G; Data S13). Notably, these pathways were driven by a significant upregulation of human leukocyte antigen genes *HLA-A*, *HLA-B*, *HLA-C*, and *B2M* (Figure 6H). These data indicate increased MHC-I expression in *SNCA* A53T iMGL, consistent with enhanced antigen presentation and a greater ability to activate T cells and engage adaptive immunity. As in the scRNAseq analysis, pathways involved in the mitochondrial electron transport chain were significantly downregulated (Figure 6I), driven by suppression of mitochondrial-encoded transcripts (Figure 6J), reinforcing the link between inflammatory activation, antigen presentation, and metabolic suppression in *SNCA* A53T iMGL.

Given the importance of secreted cytokines in coordinating and amplifying inflammatory responses, we next measured cytokine release from *SNCA* A53T and control iMGL. *SNCA* A53T iMGL secreted significantly higher levels of IFN-γ, IL-2, IL-4, IL-12p70, IL-13, and TNF-α compared with controls (Figure 6K). Whilst both genotypes secreted cytokines above basal levels following LPS stimulation (Figure S17), the fold-change relative to basal was significantly greater in controls for IFN-γ, IL-2, IL-13, and TNF-α, suggesting that the pre-activated state of *SNCA* A53T iMGL reduces their capacity to mount further responses to secondary inflammatory stimuli.

Together, these findings support a model in which dysregulation of endogenous *SNCA* expression specifically within microglia represents a potential disease-driving pathway in PD, triggering antiviral and interferon responses, MHC-I upregulation, and a metabolic shift away from oxidative phosphorylation, changes that closely mirror the immune–metabolic alterations observed in PD cortical regions at Braak stage 3–4.

### αSyn oligomers directly induce inflammatory activation in iMGL

The increased αSyn oligomer density observed in microglia in the cingulate cortex of mid-stage PD, together with the associated inflammatory transcriptomic signature, suggested a direct link between proteinopathy and microglial activation. Whilst our snRNAseq data indicate endogenous microglial SNCA as one potential source of intra-microglial oligomers, another possibility is uptake of extracellular αSyn species released from neighbouring cells. We therefore tested whether exogenous αSyn monomers could be internalised and form aggregates *de novo* within iMGL (Figure 7A). Control iMGL readily internalised fluorescently labelled monomeric αSyn, with wheat germ agglutinin (WGA) staining confirming intracellular localisation (Figure 7B). We have previously developed and validated an Förster Resonance Energy Transfer (FRET) assay to detect the assembly of monomeric αSyn into oligomers in cell systems.^18^ iMGL were treated with two populations of fluorescently labelled αSyn monomers, one conjugated to Alexa Fluor 488 (AF488) and the other to Alexa Fluor 594 (AF594). The presence of a FRET signal within iMGL confirmed the *de novo* formation of αSyn oligomers from monomers, for both WT and A53T proteins (Figure 7C). A53T-treated cells showed higher mean aggregate FRET intensity than WT-treated cells, driven by a greater proportion of high-intensity aggregates (Figures 7Di–Dii). Single-molecule FRET on iMGL lysates further confirmed the presence of oligomers 24 hours after monomer addition. Interestingly, these oligomers were found to persist within the cells for up to 10 days in culture (Figure S18).

**Figure 7.**
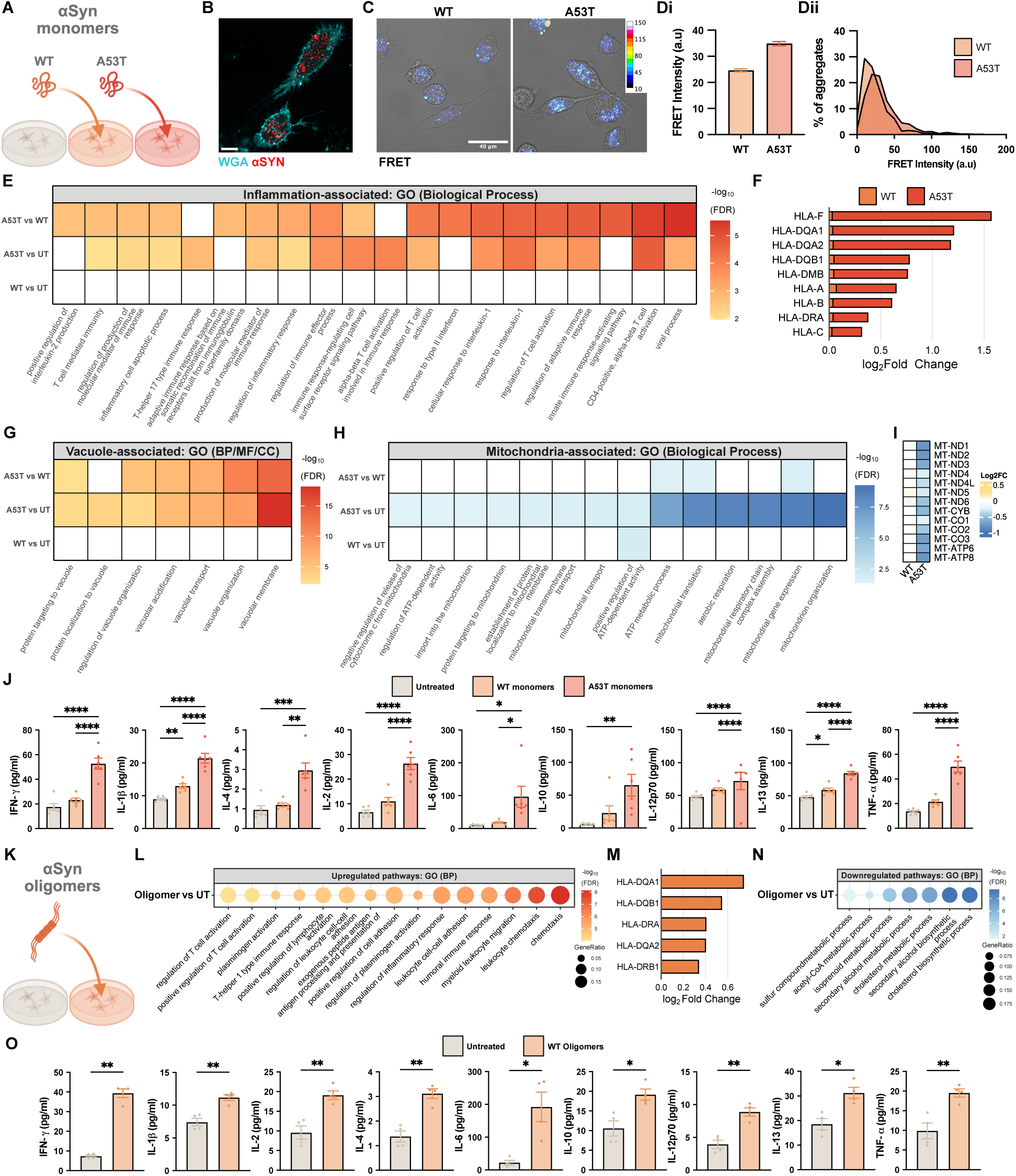
αSyn oligomers in human microglia recapitulate early PD cortical immune-metabolic signatures. (A) Schematic of experimental design of iMGL from controls treated with WT or A53T αSyn monomers. (B) Representative image of live-cell imaging of fluorescently-labelled αSyn monomers (red) and the dye Wheat Germ Agglutinin (WGA, cyan, cell plasma membrane) to show intracellular localisation of αSyn within iMGL. (C) Representative brightfield images with FRET heatmaps after incubation with WT or A53T monomeric αSyn for 24 hours. Colourbar indicates FRET intensity (a.u). (Di) Quantification of the mean FRET intensity after incubation with WT or A53T monomeric αSyn for 24 hours. (Dii) Distribution of αSyn aggregate FRET intensities after incubation with WT or A53T monomers for 24 hours. (E) Gene Ontology pathway enrichment of inflammation-associated terms from upregulated DEGs in bulk RNA sequencing of iMGL treated with WT or A53T αSyn monomers for 24 hours. (F) Log2FC for significantly upregulated MHC-1 and MHC-2 genes in iMGL treated with WT or A53T monomers compared to untreated cells. (G) Gene Ontology pathway enrichment of vacuole-associated terms from upregulated DEGs in bulk RNA sequencing of iMGL treated with WT or A53T αSyn monomers for 24 hours. (H) Gene Ontology pathway enrichment of mitochondria-associated terms from downregulated DEGs in bulk RNA sequencing of iMGL treated with WT or A53T αSyn monomers for 24 hours. (I) Log2FC for mitochondrial-encoded genes in iMGL treated with WT or A53T αSyn monomers for 24 hours. (J) Cytokine panel measured by ELISA, from conditioned media from control iMGL untreated or exposed to WT or A53T αSyn monomers for 24 hours. (n = 6 iPSC donor lines; data represent mean ± SEM; repeated-measures one-way ANOVA with Tukey’s multiple comparison test; *p < 0.05, **p < 0.01, ***p < 0.001, ****p < 0.0001). (K) Schematic of experimental design of iMGL from controls treated with WT αSyn oligomers. (L) Gene Ontology pathway enrichment of top 15 terms from upregulated DEGs in bulk RNA sequencing of iMGL treated with αSyn oligomers for 24 hours. (M) Log2FC for significantly upregulated MHC-2 genes in iMGL treated with αSyn oligomers compared to untreated cells. (N) Gene Ontology pathway enrichment of terms from downregulated DEGs in bulk RNA sequencing of iMGL treated with αSyn oligomers for 24 hours. (O) Cytokine panel, measured by ELISA from conditioned media from control iMGL untreated or exposed to WT oligomeric αSyn for 24 hours. (n = 4 iPSC donor lines; data represent mean ± SEM; paired t-test; *p < 0.05, **p < 0.01).

To assess the transcriptional consequences of *de novo* aggregation, we performed bulk RNA sequencing on iMGL exposed to WT or A53T monomers for 24 hours. WT-treated cells primarily upregulated pathways related to endosomal trafficking and GTPase-mediated signalling (Figure S19), consistent with uptake and intracellular processing, but did not induce inflammation-associated signatures. In contrast, A53T-treated iMGL exhibited robust activation of inflammatory pathways, including viral response, T cell activation and interleukin signalling (Figure 7E), as well as upregulated type I and type II interferon signalling (Data S15). This was accompanied by marked upregulation of MHC-I (HLA-A, HLA-B, HLA-C) and MHC-II (HLA-DQ family, HLA-DMB, HLA-DRA) genes in the A53T, but not WT, condition (Figure 7F). Antigen presentation may be facilitated through a vacuole-associated pathway.^53,55^ Interestingly, we defined an iMGL cluster from our scRNAseq dataset that was defined by antigen processing, vacuole pathways and PD risk (Cluster 4). This cluster also showed significant upregulation of *CTSS* (cathepsin S), a master regulator for antigen processing through the vacuole-pathway.^53–55^ Notably, in our bulk RNA sequencing of the Braak stage 3–4 brain we see a significant upregulation of *CTSS* in PD (Data S2), alongside enrichment of vacuole-associated pathways, including Vacuolar Localization, Vacuolar Lumen, Lytic Vacuole Membrane, and Vacuolar Membrane (Data S3). Here, in A53T-treated iMGL, we observed significant upregulation of pathways involved in vacuole organisation, vacuole acidification, and protein targeting to vacuoles (Figure 7G) with vacuolar membrane the most significantly enriched term (FDR-corrected p = 7.06×10^-19^; Figure 7G, Figure S19), none of which were altered by WT αSyn.

Whilst the upregulated pathways in WT- and A53T-treated cells were largely distinct (Figure S19), the most significant downregulated pathways were shared, involving DNA organisation, chromosome segregation, and cell cycle progression (Figure S20), suggesting αSyn uptake may shift proliferative microglia toward a phagocytic state. Unique to A53T treatment was pronounced suppression of mitochondrial processes, including mitochondrial organisation, respiratory chain complex assembly, ATP metabolism, and mitochondrial gene expression (Figure 7H), driven by downregulation of mitochondrial-encoded transcripts (Figure 7I), consistent with an interferon-driven metabolic shift.

Functionally, A53T-treated iMGL secreted significantly higher levels of all measured cytokines (IFN-γ, IL-1β, IL-2, IL-4, IL-6, IL-10, IL-12p70, IL-13, and TNF-α) compared to untreated cells, and all except IL-10 were significantly elevated relative to WT-treated cells (Figure 7J). In contrast, WT αSyn induced only IL-13 and IL-1β secretion. These findings indicate that the increased oligomerisation driven by the A53T mutation, rather than αSyn uptake and processing per se, is the critical trigger for the potent pro-inflammatory phenotype in microglia.

Given that A53T αSyn monomers drive both oligomer formation and inflammation, we next tested whether pre-formed recombinant WT αSyn oligomers could directly induce a similar response (Figure 7K). Oligomer treatment of control iMGL for 24 hours induced strong upregulation of inflammatory and immune cell migration pathways (Figure 7L; Data S17), including leukocyte chemotaxis (FDR-corrected p = 2.77×10^-8^), regulation of inflammatory response (FDR-corrected p = 2.52×10^-6^), and antigen processing and presentation of exogenous peptide antigen (FDR-corrected p = 1.27×10^-5^), along with significant increases in MHC-II genes, including *HLA-DQA*, *HLA-DQB*, *HLA-DRB*, and *HLA-DRA* (Figure 7M). As with A53T monomer treatment, oligomer exposure also suppressed key metabolic pathways, including cholesterol biosynthetic process (FDR-corrected p = 3.62×10^-10^) and acetyl-CoA metabolic process (FDR-corrected p = 2.49×10^-3^; Figure 7N). Functionally, oligomer-treated iMGL secreted significantly higher levels of all measured cytokines (IFN-γ, IL-1β, IL-2, IL-4, IL-6, IL-10, IL-12p70, IL-13, and TNF-α) compared to untreated cells (Figure 7O), with a concentration-dependent relationship between oligomer dose and cytokine release (Figure S21). Together, these data indicate that αSyn oligomers, whether formed intracellularly in microglia or acquired from their extracellular environment, are potent drivers of microglial activation, characterised by vacuolar antigen-presentation pathways, inflammatory gene expression, and mitochondrial suppression.

## DISCUSSION

Most human post-mortem studies of PD have focused on Braak stage 5–6, when extensive LB pathology, neuronal loss, and gliosis dominate the cellular and molecular landscape.^1,3,66^ Whilst invaluable for understanding late-stage neurodegeneration, such analyses are inherently limited in resolving early pathogenic processes, as molecular signatures at this stage are heavily shaped by secondary responses that obscure initiating events. In this study, we used a multi-region approach to examine Braak stages 3– 4, spanning a pathological gradient from severely affected subcortical regions to cortical areas largely spared from LB deposition. This design provides a unique opportunity to capture the molecular and cellular landscape that precedes widespread LB accumulation and large-scale neurodegeneration. To this end, we combined advanced transcriptomic profiling of the early-mid PD brain with detailed mapping of the proteopathic state of αSyn and directly modelled the impact of αSyn oligomers on the human transcriptome *in vitro*.

Together our analyses yielded four principal findings. First, robust inflammatory activation occurs in pathologically unaffected human brain regions before widespread LB pathology. Second, although this inflammatory signature is present across multiple cell types, microglia emerge as a central driver of the early inflammatory response in PD. Third, this early inflammatory phase is driven, in part, by αSyn misfolding and oligomerisation, which can arise within microglia. Finally, synucleinopathy-associated microglial states represent distinct and diverse phenotypes that reprogram microglial function towards antigen presentation and pro-inflammatory activity, thereby contributing to a neuroinflammatory milieu that precedes neuronal dysfunction and LB pathology.

Microglia are the specialised resident immune cells of the brain, dynamically surveying their microenvironment for molecular indicators of cellular stress, damage, infection, or the accumulation of misfolded proteins. In response to such cues, they rapidly adopt activated states that coordinate the inflammatory response through cytokine release, antigen presentation, and recruitment of other immune cells.^67,68^ Whilst these responses may be protective, sustained or maladaptive activation can conversely lead to chronic neuroinflammation that disrupts neuronal health and contributes to neurodegeneration. In PD, it remains unclear the extent to which microglial activation is neuroprotective or contributes to neurodegeneration, and how this balance may shift across the disease course.^68–70^

Alterations in the microglial population, or their activation state, are well documented in PD, with converging evidence from post-mortem histology, *in vivo* imaging, transcriptional profiling and experimental models. Immunohistochemical studies demonstrate increased microglia expressing MHC class II molecules in the substantia nigra and cortical regions, including the cingulate cortex, of PD patients, irrespective of the extent of LB pathology.^71,72^ PET imaging for peripheral benzodiazepine binding sites which mark activated microglia demonstrated increased microglial activation across brain regions in PD.^73^ More recently, snRNAseq has revealed an increased proportion of microglia in the substantia nigra with transcriptional profiles consistent with a pro-inflammatory state.^5^ Experimentally, aberrant αSyn can drive inflammatory microglial activation in murine cell lines,^74^ primary cultures,^75^ and human models.^76^ Furthermore, selective microglial αSyn overexpression in a mouse model induces both microglial activation and neuronal degeneration.^13^ Despite robust evidence from multiple methodologies implicating microglial activation as a critical feature of PD pathology, the timing of its onset, its relationship to αSyn aggregation, and the precise triggering mechanism(s) remain unclear.

Here, through snRNAseq of the PD cortex, we identified eight immune cell populations, five of which represented distinct microglial states. The cortical signature in PD was defined by a selective enrichment of FOXP2_OXR1 microglia, a state marked by inflammatory pathways such as lymphocyte recruitment, high TLR expression, and unexpectedly high *SNCA* expression. Although *SNCA* is typically associated with neurons, this raises the possibility that αSyn aggregates within microglia may originate from endogenous production as well as from phagocytosed neuronal αSyn. Beyond microglia, inflammatory signatures were also detected in other cell types, including excitatory neurons, where genes associated with innate immune signalling were upregulated. Whilst microglia actively orchestrate inflammatory responses,^67,68^ neurons influence immune activity primarily through signalling to glia,^77^ rather than by directly secreting pro-inflammatory cytokines or engaging in classical immune effector functions. This suggests that early microglial activation in the PD brain may reshape the local cellular milieu in ways that directly influence neuronal function.

Consistent with the idea that changes to the cellular milieu occur early in disease, our spatial transcriptomics of the cingulate cortex reveals that in PD, microglial neighbourhood relationships may shift, with microglia showing increased proximity to other cell types, potentially facilitating broader inflammatory crosstalk. Immune cells communicate via multiple mechanisms, including direct cell–cell contact, secretion of cytokines and chemokines, and release of extracellular vesicles, thereby influencing each other’s activation state, migration, and function.^78,79^ In PD we see an increase in receptor clustering, the colocalisation of receptors at specific membrane sites to amplify signalling,^80^ and an upregulation of *APOE*, which may serve here to propagate inflammatory signals across immune cells, potentially driven by the increased αSyn oligomer load, resulting in elevated immune–immune interactions.

Microglia are activated by diverse external and internal cues, including pathogen-associated molecular patterns such as LPS,^81^ environmental toxins,^82^ and endogenous misfolded proteins.^68^ Misfolded αSyn has been shown to induce microglial activation by engaging pattern-recognition receptors such as TLR2 and TLR4, triggering pro-inflammatory signalling cascades.^83,84^ To investigate the relationship between microglial activation and αSyn pathology, we combined conventional LB detection with sensitive oligomer imaging using our state-of-the-art ASA-PD pipeline.^24^ Whilst traditional immunohistochemistry reliably detects late-stage insoluble αSyn aggregates, predominantly neuronal LBs or Lewy neurites, emerging evidence highlights the pathogenic importance of smaller soluble assemblies, termed oligomers. We previously optimised ASA-PD to sensitively detect these nanoscale species in human brain, and here applied it alongside cell-type markers. Although αSyn oligomers were detectable in neurons, we observed a selective increase in phosphorylated αSyn oligomer density within microglia, but not neurons, in the cingulate cortex of PD cases, a region with minimal LB pathology.

Together, these findings in Braak stage 3–4 brains, showing that early αSyn oligomers accumulate within microglia—the cell type activated in early PD—raise the hypothesis that αSyn itself may act within microglia to directly trigger inflammatory responses. To test this causal link between αSyn oligomers and microglial activation, we developed hiPSC models of endogenous αSyn pathology (*SNCA* A53T), and exposure to exogenously applied αSyn in its monomeric and oligomeric forms. iMGL are a robust model for interrogating αSyn-microglia interactions, closely mapping onto microglial transcriptional states observed in the human PD brain. Microglia derived from PD patients carrying the *SNCA* A53T mutation exhibited a constitutively activated inflammatory phenotype, characterised by cytokine secretion and immune activation reminiscent of that identified in Braak stage 3–4 PD brain. This directly links dysregulated αSyn in microglia to the emergence of an inflammatory state in the absence of large aggregates. To further dissect the contribution of αSyn to microglial activation, we applied exogenous αSyn in its monomeric and oligomeric forms. iMGL efficiently phagocytosed αSyn monomers, which subsequently oligomerised intracellularly. This was detectable within 24 hours and was accelerated by the *SNCA* A53T mutation, consistent with its known increased oligomerisation kinetics.^48,49^ A53T monomeric αSyn, but not WT monomers, triggered a strong inflammatory response characterised by upregulated interleukin signalling and a shift to antigen presentation, accompanied with increased secretion of inflammatory cytokines. WT oligomers, in contrast to WT monomers, also induced this inflammatory phenotype, demonstrating that oligomeric αSyn, whether generated endogenously from A53T monomers or taken up exogenously, is sufficient to drive microglial activation. This inflammatory phenotype may occur directly through oligomers being sensed as damage-associated molecular patterns (DAMPs) or be the result of cellular damage induced through membrane-permeabilising assemblies.

scRNAseq of iMGL identified a cluster enriched for PD-associated genes, including *SNCA*, antigen-processing and vacuolar pathways, and the lysosomal protease CTSS (cathepsin S), a master regulator of vacuole-dependent antigen presentation. Across all models (*SNCA* A53T mutation, A53T monomer treatment, and WT oligomer treatment) we observed significant upregulation of genes associated with MHC class I and MHC class II. Interestingly, vacuolar pathways were also significantly enriched in the A53T monomer treatment, suggesting that αSyn oligomerisation within microglia drives a switch to antigen-presentation through vacuole-processing pathways. Consistently, bulk RNA sequencing of Braak stage 3–4 PD cortex also revealed upregulation of *CTSS*, MHC class II genes, and vacuole-associated terms, further implicating this process in early PD brain. Accompanying the inflammatory signature in both human brain and *in vitro* models, we also observed significant downregulation of pathways associated with mitochondrial function, consistent with our previous proteomic study of Braak stage 3–4 PD brain. Such metabolic and mitochondrial suppression is likely to reflect interferon-driven reprogramming of microglial bioenergetics, which may sustain chronic inflammation. This immune reprogramming at the earliest phase of disease may ultimately facilitate recruitment of the adaptive immune system downstream, aligning with evidence of T cell activation and dysregulation in PD.^31,85–87^

In summary, our study defines the earliest cortical phase of PD, the prodrome to cognitive decline, in which neuroinflammation, microglial activation, and intracellular αSyn oligomer abundance act in concert to cause pathology. Critically, we reveal the presence of inflammation prior to any LB formation suggesting that inflammatory responses are not merely secondary to cellular stress and neuronal loss but may actively drive disease progression. Whilst neuroinflammation has been well established within PD, it has been challenging to disentangle secondary bystander late consequences from early causal effects. This study provides evidence for neuroinflammation preceding neuronal dysfunction, revealing an early ‘cellular’ phase of the condition that may be causal. Microglial activation can be triggered by a variety of pathogens, environmental factors, as well as endogenous damage signals from misfolded proteins, and it is possible that several different triggers converge on the microglia during the early stages of motor PD, and perhaps even in prodromal PD, where there is *in vivo* imaging evidence of microglial activation.^88^ Here, using human post-mortem tissue and human *in vitro* models, we defined one trigger for the microglial activation to be αSyn itself, and specifically its oligomeric state. Furthermore, we demonstrate how endogenous and exogenous αSyn may be recognised by microglia, how microglia may undergo metabolic and inflammatory reprogramming, and how it induces an antigen presenting state, effects that lead to the recruitment of other parts of the innate and adaptive immune system. Once microglial activation has been triggered, irrespective of the initial trigger, we propose that a toxic positive feedback loop is established, in which oligomer formation and inflammation further exacerbate each other, and impact neuronal function, leading to spread and progression.

Our findings highlight the critical need to understand the role of neuroimmune mechanisms in causing disease at early stages, and exacerbating progression of existing PD pathology. Understanding these cellular mechanisms may pave the way to therapeutics that target the immune phenotypes of microglia, that could provide strategies to slow down or prevent the cortical pathology that underlies the cognitive symptoms and development of dementia in PD.

## RESOURCE AVAILABILITY

### Lead contact

Further information and requests for resources and reagents should be directed to and will be fulfilled by the lead contact, Prof. Sonia Gandhi

### Data and code availability

All bulk and single-cell RNA sequencing sample preprocessing and analysis code will be made available upon publication. Raw fastq files for Braak 3-4 PD sequencing are available upon request through the ASAP CRN Cloud platform Raw imaging data of post-mortem brain is archived independently at the Imaging Data Resource (IDR), with a DOI currently being generated (∼2 TB). [DOI under review at IDR]. Processed data quantifying the cellular density of αSyn is available as a structured database on Zenodo (https://doi.org/10.5281/zenodo.16421701). Exemplar data and analysis code for post-mortem imaging, suitable for reproducing key findings, are provided alongside this publication “pyRASP_copy_for_paper.zip” and on Zenodo at https://doi.org/10.5281/zenodo.16421701. Code supporting the post-mortem imaging is available both at GitHub (https://github.com/TheLeeLab/pyRASP) and an archived version with example data is available at Zenodo at https://doi.org/10.5281/zenodo.16421701.

## METHOD DETAILS

### Post-mortem sample selection

All brain samples used in this study were provided by the Parkinson’s Disease UK tissue bank with informed consent to use the material for research available for each donation. Selection of cases and controls was performed based upon disease status of each individual (Control or Parkinson’s disease) found in the clinical notes in patient records, no dementia incidence, post-mortem delay (less than 24 hours) and an absence of any confounding pathology. The cases with the highest RIN were selected from those that met these criteria. Formalin-fixed paraffin-embedded sections and frozen sections or chips of tissue were taken as necessary for all further downstream analyses and stored at room temperature or -80°C respectively until use.

Only those that were reported not to have dementia and mapped to the Brainstem predominant or limbic transitional stage were selected. PD cases had no confounding pathology, but some controls did have some minimal signs of age-related changes. The two cohorts, namely control and PD, had comparable mean age of death (age_controls_ = 80.4; age_PD_ = 79.5) and both groups had a higher representation of male subjects relative to female (ratio_controls_ = 1.857; ratio_PD_ = 2.0).

### Post-mortem bulk RNA sequencing

#### Bulk RNA isolation

Approximately 50-100 g of frozen tissue was used per brain sample for RNA extraction. Tissue lysis was performed using QIAzol and RNA was extracted using the RNeasy 96 Kit (Qiagen) with an on-membrane DNase treatment, as per manufacturer instructions. Samples were quantified by absorption on the QIAxpert (Qiagen), and RNA integrity number (RIN) measured using the Agilent 4200 Tapestation (Agilent Technologies).

#### Generation of bulk RNA sequencing data

Stranded cDNA libraries were constructed with the KAPA mRNA HyperPrep Kit (Roche) used as per manufacturer instructions. More specifically, 500ng of total RNA was used as input for each sample and xGen Dual Index UMI Adapters (Integrated DNA Technologies, Inc.) were added to each sample to minimise read mis-assignment when performing post-sequencing sample de-multiplexing. Libraries were multiplexed on S2 and S4 NovaSeq flow cells for paired-end 150 bp sequencing on the NovaSeq 6000 Sequencing System (Illumina) to obtain a target read depth of 110 million paired-end reads per sample. Sequenced reads were de-multiplexed and FASTQ files were generated using the BCL Convert software (Illumina). FASTQ files were run through a Nextflow pipeline^89^ that performed quality control (QC) of reads and aligned the samples to the genome and transcriptome (GRCh38 amd Gencode v41 annotations). Briefly, initial QC was performed using fastp (v0.23.2)^90^, which removed low quality reads and bases, as well as the trimming of adapter sequences. Filtered reads were aligned to the transcriptome using Salmon (correcting for sequence-specific, fragment GC-content and positional biases; v1.9.0,^91^ using the entire genome as a decoy sequence. For splicing analyses, reads were aligned to the genome using STAR (v2.7.10a, with two-pass mode). FastQC (v0.11.9) was used to generate read QC data before and after fastp filtering. RSEQC (v4.0.0),^92^ Qualimap (v2.2.2a)^93^ and Picard (v2.27.5) were used to generate alignment QC metrics. MultiQC (v1.13),^94^ was used to visualise and collate quality metrics from all pipeline modules. Full details for the pipelines and parameters used for each module can be found here: https://github.com/Jbrenton191/RNAseq_splicing_pipeline.

#### Sample quality control, covariate selection and outlier removal

Samples with a low read count (<60M reads aligning to the transcriptome), a high % of reads aligning to intronic regions of the genome (>30%) or a deviant average GC content (<40% or >65%) were removed from further analysis. Transcriptome aligned reads generated from Salmon were used to identify covariates and then outlier samples. For both processes, salmon read files were imported into R (v4.3.0) and converted into gene-level count matrices using the tximport (v1.30.0) and DESeq2 (v1.42.1) packages.

Covariates and outliers were detected with the same method as above. The covariate selection was run once across all bulk samples and covariates passing the selection criteria for all samples were taken forwards. The outlier analysis was run using these covariates and z-scores were calculated for each sample against the sample population from the same brain area.

#### Differential gene expression analysis

Gene-level count matrices were filtered for genes with 0 counts in at least one sample in either the disease or control group. Analysis was performed using limma/voom (ref/ versions). Briefly, voom was used to shrink the dispersion and the duplicateCorrelation function was used to remove the effect of each individual to allow multiple brain areas from the same individual to be modelled together. These functions were applied twice as recommended by the developers. Group and brain area factors were merged into a single term to allow the model to be run once and all comparisons to be extracted from that model. Contrasts were run between groups for each brain area, and multi-region comparisons incorporating either all brain regions, cortical regions (anterior cingulate cortex, frontal cortex, parahippocampal gyrus, parietal cortex and temporal cortex) or subcortical regions (subtantia nigra, putamen and caudate). Differentially expressed genes were extracted for each contrast using the eBayes and topTable limma functions and were classified as differentially expressed if their Benjamini and Hochberg FDR adjusted p value was ≤ 0.05. Differentially expressed genes were classified as upregulated if their log_2_ fold change was > 0 and downregulated if < 0.

#### Pathway analysis

Gene Ontology (GO), Kyoto Encyclopedia of Genes and Genomes (KEGG) and Reactome Knowledgebase databases were interrogated for over-representation pathways analysis using clusterProfiler (v4.10.1). The compareCluster function was used to perform over-representation analyses for all ontologies across brain areas or stages of disease. Pathways were counted as significant if their FDR-adjusted p value was ≤ 0.05. The number of GO pathways were reduced by semantic similarity using the GOSemSim and rrvgo R packages. The “Wang” method, a graph-based measure of semantic similarity, and a similarity threshold for grouping terms of 0.7 were used. The reduction was performed separately for each of the datasets.

#### Expression Weighted Cell Type Enrichment

Cell type enrichment of the differentially expressed genes was calculated using Expression Weighted Cell Type Enrichment (EWCE)^95^. EWCE determines if genes of interest have significantly higher expression in certain cell types than might be expected by chance. Bootstrapped gene lists controlled for transcript length and GC-content were generated with EWCE iteratively (n = 50,000) using “bootstrap_enrichment_test()” function. This function returns the probability of enrichment using standard deviations from an expression mean across celltypes and FDR corrected p values (q-values). A pre-formatted cell specificity matrix for the Human Multiple Cortical Areas SMART-seq dataset (https://portal.brain-map.org/atlases-and-data/rnaseq/protocols-human-cortex) was downloaded from https://github.com/RHReynolds/MarkerGenes and used as the single-cell reference. The excitatory and inhibitory neuronal subtype with the lowest q value and largest standard deviation from the cell type mean was selected to represent either major neuronal class.

#### Gene set overlap analysis

To assess the identity of our gene expression signatures, we overlapped our bulk transcriptomic DEGs stratified by anatomical location (i) all regions (up N = 847, down N = 384), (ii) cortical (up N = 609, down N = 262), and (iii) subcortical (up N = 340, down N = 309), against publicly available datasets defining microglial states. Specifically, signatures of Disease Associated Microglia (DAM, up N = 273, down N = 28),^29^ Human AD Microglia (HAM, up N = 44, down N = 21),^28^ as well as pseudobulked upregulated DEGs comparing AD or LB disease vs. control from DAM (N = 1327 and N = 560, respectively) and Disease Inflammatory Macrophage (DIM) clusters (N = 12 and N = 7, respectively) within the Human Microglia Atlas (HuMicA).^30^ Mouse symbols were converted to human via biomart (release 113, GRCm39).^96^ Gene set similarity was assessed by pairwise Fishers Exact Tests using GeneOverlap (V.1.40.0) (https://bioconductor.org/packages/release/bioc/html/GeneOverlap.html) with the universe defined as all unique genes that passed quality control (19,708), p values were then adjusted for multiple comparisons by FDR corrections within each external dataset. Heat maps were generated using ComplexHeatmap (v.2.20.0) in RStudio (R v. 4.4.1).

### Post-mortem multiome sequencing

#### Isolation of nuclei

Approximately 200–300 mg of frozen grey matter was obtained from the anterior cingulate cortex, frontal cortex, and parietal cortex. Samples were manually homogenized using a Dounce tissue grinder homogenizer in 1 ml of HB, with 25 strokes each of Grade A and Grade B, before being added to 650 µl of HB. Each tube was washed with 1 ml of HB to collect any remaining homogenate, resulting in a total volume of 2.65 ml of HB added. After homogenization, 2.65 ml of GM was added to each sample, and the tubes were inverted to ensure even distribution. 5.25 ml of each sample was layered onto a 4 ml volume of a 29% OptiPrep cushion within their respective ultracentrifuge tubes. Samples were then matched and balanced according to rotor order and underwent ultracentrifugation for 30 min at 4°C at 7700 rpm (SW41Ti swinging-bucket rotor, Optima-XPN-90).

Post-centrifugation, supernatants were collected on ice using a 1 ml Gibson pipette tip and discarded, taking care not to agitate the nuclear pellet. Pellets were then resuspended in 1 ml of ATAC-seq wash buffer before being passed through chilled 35 µm Cell Strainer Falcon Tubes (352235, Corning). Eluents were transferred into chilled 2 ml LoBind Eppendorf tubes and centrifuged at 4°C at 500 rcf for 5 min. Supernatants were discarded, and pellets were resuspended in 400 µl of ATAC-seq wash buffer. Eluents were again passed through chilled 35 µm Cell Strainer Falcon Tubes (352235, Corning), transferred to chilled 2 ml LoBind Eppendorf tubes, and centrifuged at 4°C at 500 rcf for 5 min. Finally, supernatants were discarded, and the samples were resuspended in 30–100 µl of ATAC-seq resuspension buffer, depending on pellet size.

#### Multiome library preparation

The concentration and viability of the single nuclei suspension was measured using acridine orange (AO) and propidium iodide (PI) and the Luna-FX7 Automatic Cell Counter. Approximately [5000-20,000] nuclei were transposed, then loaded on Chromium Chip and partitioned in nanolitre scale droplets using the Chromium Controller and Chromium Next GEM Single Cell Reagents (Chromium Next GEM Single Cell Multiome ATAC + Gene Expression Reagent Kits User Guide, User Guide, CG000338).

A pool of 736,000 10x Barcodes was sampled to separately and uniquely index the transposed DNA and cDNA of each individual nucleus. ATAC and gene expression (GEX) libraries were generated from the same pool of pre-amplified transposed DNA/cDNA and sequenced using the Illumina NovaSeq 6000. Sequencing read configuration: 28-10-10-90 (GEX), 50-8-24-49 (ATAC). The 10x Barcodes in each library type are used to associate individual reads back to the individual partitions, and thereby, to each individual nucleus.

#### Multiome data preparation with panpipes

Genome alignment was performed on sequencing outputs using cellranger-ARC (v. 2.0.1) using reference genome ENSEMBL (v 107, generated with cellranger-ARC v. 2.0.1). Outputs were then aggregated using cellranger-ARC function aggr. The aggregated data was then ingested to panpipes for QC, preprocessing and clustering^97^. This consisted of a total of 955,571 nuclei. Following inspection of QC metrics, nuclei were filtered based on whether they passed conventional QC metrics, namely, percentage of mitochondrial transcripts in RNA counts (<5%), percentage of ribosomal transcripts in RNA counts (<5%), percentage of haemoglobin genes (<5%), total UMI counts (<20,000), number of genes by counts (20<n<17500), number of cells per gene (>20). The data was then normalised, followed by log1p transformation. Dimensionality reduction was applied (N_PCs_ = 100) and batch correction was applied on gene-expression. Finally, community detection with the leiden algorithm was performed.

Major cell types were detectable with cell type markers (Astrocytes: *AQP4*, *GFAP*; EndoMural cells: *CLDN5*; Neurons: *GABRB2*; Inhibitory neurons: *GAD2*; Immune cells: *CD74*, *PTPRC*, *TREM2*, *APOE*; Oligodendrocytes: *PLP1*, *MBP*; OPCs: *PDGFRA*, *BCAS1*). These major cell types were then partitioned and subclustered further separately, similarly to other single nuclear RNA sequencing approaches.^98–100^ For this, each cell type was batch corrected separately using the same batch correction and community detection approach. Using clustree cluster hierarchy, leiden resolutions were chosen per cell type based on clustering stability.

Cluster annotations were performed using a combination of cell markers and the highly variable genes calculated for each cluster, as implemented by scanpy^101^. Based on a combination of data-driven highly variable genes and marker genes, cell types were annotated across 4 levels of increasing resolution, from level 1 (major cell type) to level 4 (high resolution cell type/state). During the annotation process clusters which exhibited evidence of being doublets (co-expression of marker genes, high scrublet score) and/or high rates of metrics associated with low quality were excluded. Following all QC and annotation steps, a total of 740,157 nuclei remained. snRNA-seq data were then pseudobulked for downstream analyses, since this addresses pseudoreplication bias in single nuclear data^102^. The aggregateToPseudoBulk function from the dreamlet R package (version 0.99.16)^103^ was used to sum gene expression counts of nuclei by participant ID, brain region (cingulate, frontal, parietal) and cell type annotation. This was performed separately for each annotation level.

#### Selection of covariates for gene expression

Covariates driving significant variation in the gene expression data were discovered using an unbiased, data-driven approach. First, all technical (sample preparation, nuclei extraction, sequencing, mapping), clinical, and histopathological metadata were combined. Uninformative or unusable variables were removed, including IDs, those with zero variance and those that were a proxy for disease group. We then performed a second tier of filtration, removing variables with high missingness and selecting one from variable pairs with high collinearity. Of the variables remaining, those with missing values underwent mean imputation (for numericals), or common value imputation (for categoricals). For assessing pairs of continuous variables, collinearity was assessed using the caret R package (version 6.0-94), with a Spearman’s rank correlation coefficient cut-off of set at 0.7. For every correlation above this threshold, the variable with the lowest average correlation was kept and the other covariate removed. To assess collinearity of pairs of categorical variables, a χ-squared test was used. For pairs of numeric and categorical variables, we used the Kruskal-Wallace test and for pairs of numeric variables we used a Spearman’s rank correlation test. The resulting p values were assessed using a significance threshold adjusted for multiple comparisons (p < 0.05/(number of tests)). Variables were then removed from pairs that demonstrated a significant categorical/categorical or categorical/continuous relationships. The contribution to variance in gene expression of the remaining variables was assessed by two methods: (1) variancePartition;^104^ variancePartition R package) and (2) Principal Component Analysis (PCA; prcomp R package). VariancePartition assesses the contribution of variance of putative covariates at the gene level, whereas PCA-based dimensionality reduction generates per-sample eigenvectors which can then be correlated with putative covariates to assess their relative importance. We employed these two methods to determine our final covariates, recognising the need to control not only for covariates with global impacts on expression, but also those that may confound the expression of small subsets of genes. Putative covariates were then assessed against three metrics: (1) variancePartition: the maximum variance explained percentage, providing an assessment of whether a covariate comprises a large proportion of the variation of any gene(s), (2) variancePartition: the 3rd quartile of the variance explained distribution, providing an assessment of whether a covariate comprises a sizeable proportion of the variation of a larger proportion of genes and (3) the Spearman’s rank correlation coefficient between PC-X and covariate. For each metric, data-driven cut-offs were set using visualisation and cross-cell type concordance was assessed. Covariates passing the cut-offs in >50% (4/7) of cell types were selected as the final covariates. This resulted in the selection of the following five covariates: sex, batch, downsampled, merged samples, repeated samples and total deduplicated percentage (see Table S2).

#### Cell type proportions

For the cross-group analysis, differences in cell type proportions across groups were assessed using crumblr (https://github.com/GabrielHoffman/crumblr/). Briefly, occurrences of each cell type in each sample were summed. The data was covariate corrected (excluding binary variables, making the final variables: sex, batch, total_deduplicated_percentage) using dream and the fit was smoothed using eBayes(). This comparison was performed separately for each brain region analysed, and pairwise comparisons were performed for controls vs PD. Finally, p values were FDR-corrected for each cell type compartment.

#### Outlier detection

Following pseudobulking, we performed outlier detection. The outlier detection method employed was adapted from Gandal et al. 2022^105^ and utilised pseudobulked, normalised (log_2_ Fragments Per Million (FPM)), covariate corrected expression values. In brief, a sample was considered an outlier if it (1) had an absolute Z-score greater than 3 for any of the top 10 principal components and (2) had a sample connectivity score of less than -2 and (3) fulfilled the first two criteria in at least 50% (4/7) of cell types. Sample connectivity was calculated using the WGCNA R package.^106^

#### Differential gene expression analysis

Differential expression analysis was carried out using the Dreamlet R package (version 0.99.16).^103^ Following this, pseudobulked expression values were normalised (per cell type) using the processAssays function, setting the ‘min.count’ argument (minimum number of reads for a gene to be considered expressed in a sample) to 10. The resulting normalised counts were then passed to dreamlet’s dreamlet function to perform differential expression. Contrasts were made for the two disease groups, contol and PD, for each of the three tissues (cingulate, parietal and frontal). Also supplied to the dreamlet function were normalised expression values and the following formula:

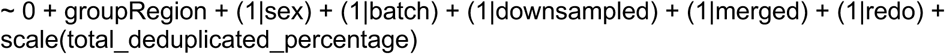

#### Gene ontology and pathway enrichment analysis

Pathway enrichment analysis was performed using the clusterProfiler R package (version 4.8.2) to identify biological pathways significantly enriched in the differentially expressed genes. Gene Ontology (GO), Kyoto Encyclopedia of Genes and Genomes (KEGG) and Reactome pathway databases were used as references for enrichment. Prior to analysis, gene IDs were converted to Entrez IDs using the AnnotationDbi package (version 1.62.2). The enrichGO and enrichKEGG functions from clusterProfiler as well as the enrichPathway function from ReactomePA (version 1.44.0) were applied to identify overrepresented terms and pathways in the set of differentially expressed genes. The model passed to the geneClusters argument was as follows:

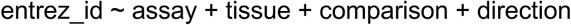

The analysis was run for each tissue (cingulate, parietal and frontal), cell type and comparison (i.e. PD vs. Control) separately. For each run, the set of all genes expressed in a given tissue and cell type was used as the background gene set for enrichment, supplied to the ‘universe’ argument. The results were adjusted for multiple testing, with terms and pathways showing an adjusted p value of < 0.05 considered statistically significant. Adjustment of p values was performed separately for each cell_type/tissue/comparison run of the analysis.

#### Familial, GWAS and PGRS implicated PD gene enrichment

To enable disease gene list enrichment analyses, we curated gene panels using publicly available datasets and standardised filtering criteria. Our approach incorporated genes implicated in PD and AD, considering both rare Mendelian disease-associated genes and common variant GWAS-implicated genes. For the PD rare Mendelian gene list, we utilised two independent sources. The first was Blauwendraat et al. (2020),^107^ from which we extracted genes listed in Table 1 of their manuscript, retaining only those classified as high or very high confidence for PD causation. The second source was OMIM, where we performed a keyword search for “Parkinson disease” and retained only genes with strong disease associations. Genes with weak, disproven, or uncertain evidence (GIGYF2, UCHL1, EIF4G1) were manually excluded. For AD, rare Mendelian genes were sourced from OpenTargets, filtering for genes associated with AD and requiring a genetic association score >0.8. For the common variant GWAS-implicated gene lists, we utilised the most recent and comprehensive datasets for each disease. For PD, we incorporated the large-scale study by Nalls,^41^ the multi-ancestry GWAS by Kim et al. (2024)^108^ and the most recent GWAS resulting from the GP2 effort.^109^ Gene symbols were mapped to Ensembl gene IDs using the org.Hs.eg.db R package, with manual curation applied to resolve missing or ambiguous identifiers. Finally, we used Fisher’s hypergeometric test in order to verify whether there was a significant enrichment of disease associated genes among differentially expressed genes. Adjustment of p values was performed for all gene lists and all cell types.

#### Characterisation of Microglia_FOXP2_OXR1 population

In order to characterise the Microglia_FOXP2_OXR1 population, which is proportionally overrepresented in PD in depth, we made use of the dreamlet analysis framework. Due to the small number of testing nuclei, the eBayes method, which accounts for outlier genes, could not be used. Therefore, we accounted for outlier genes by only including in the analysis genes that were expressed in at least 50% of samples. This allowed us to calculate differential gene expression of this population when compared to each of the other microglial population in pair-wise fashion. Finally, we employed gene set enrichment analysis to assess based on rank the upregulated pathways in this cluster.

#### snATAC-seq analysis (marker peak identification)

Using the information from the snRNA-seq data that identified high quality nuclei and provided cell type/state annotation across 4 different levels, we used <u>SnapATAC2</u>^110^ (v2.6.4) to identify marker genes for different cell states within the Immune and the Excitatory Neuron cell types. To ensure robust peak calling we assume that a cell state makes up 2% of the entire cell population (per sample) and that a feature (peak) specific to this cell state is detected at least 5% of the time. Cells falling below those thresholds (i.e. rare cell types) were removed from the subsequent peak calling.

Peaks were called at annotation level 3 with MACS3 using SnapATAC2’s function with the following parameters:

snapatac2.tl.macs3(AnnData, groupby = celltype_brainRegion_diseaseStatus, replicate = individual, qvalue = 0.1, max_frag_size = 600, nolambda = True, shift = -100, extsize = 200, blacklist = exclusion_bed,)

The resulting peaks were merged with `snap.tl.merge_peaks()` to generate a consensus peak set across cell types. This consensus peak set was then used in `snapatac2.pp.make_peak_matrix()` with the `fragment` counting strategy.

Finally, using SnapATAC2’s z-score based approach, we identified peaks that mark cell states (at annotation level 4) within the Immune and Excitatory Neuron cell types comparing each cell state against the remainder of cells within each subset cell type (e.g. `snapatac2.tl.marker_regions(AnnData_subset, groupby=’annotation_level_4’, pvalue = 0.01)`). After identifying marker peaks for microglia and excitatory cell states (`annotation_level_4`), we use <u>annotatr</u> (v 1.28.0) to assign these peaks to genomic regions and chromatin states (ChromHMM).

### Post-mortem spatial sequencing

#### CosMx slide preparation

8 μm fresh frozen brain sections were mounted onto a glass slide immediately prior to the experiment, fixed in chilled 10% NBF overnight at 4°C and processed for spatial transcriptomics according to the Nanostring fresh frozen RNA assay manual slide preparation protocol (MAN-10184-05). Briefly, slides were washed and baked vertically for 30 minutes and rehydrated with increasing concentrations of EtOH. Target retrieval was performed at 100°C for 15min after which tissue underwent protease digestion. After fiducials were applied, tissue was post fixed with 10% NBF for 1min and blocked with NHS-Acetate for 15min. In situ hybridization was performed with the CosMx™ Human 6K Discovery Panel overnight at 37°C. Following, the slides were stained with the antibody morphology stain from the Nanostring Mouse RNA Neuroscience Cell segmentation kit staining for a neuronal and histone marker, glial fibrillary acidic protein (GFAP) and DAPI. After flow cells assembly the slides were loaded into the instrument and ran according to the protocol (ID: MAN-10161-05). Run parameters were the following: panel, CosMx Hg Discovery Panel (6k); Pre Bleach config, C; Cell Segmentation Profile, Config B (neuro tissue). A total of 419 FOVs were collected.

#### CosMx data preprocessing and analysis

Cells were re-segmented using the AtoMx (v.1.3) segmentation tool according to the following basic parameters: Config B - Brain Tissue (RNA), CellDilation 0.72, CellDiameterUm: 15, NucleusDiameterUm: 15, and advanced parameters: NucleiModel: nuclei, CytoplasmModel: cyto2, MinCellSizeUm:10. Flat files were exported from AtoMx according to Nanostring Data Analysis User manual (MAN-10162-06) and imported into RStudio (R 4.4.1) as a Seurat object using the LoadNanostring function.^111–115^ FOVs including white matter were excluded from the study. Cells with counts bellow 50 and proportion of negative counts exceeding 5% were excluded. FOVs were examined using the runFOVQC function with max_prop_loss = 0.6 and max_totalcounts_loss = 0.6 (CosMx-Analysis-Scratch-Space), including only FOVs that passed the QC. A total of 68,064 high-quality cells across all tissue sections were retained for subsequent analysis. Normalisation and scaling were preformed using Seurat NormalizeData and ScaleData functions using default settings. Principal Component Analysis (PCA) was performed on the expression profile of the genes. Slides were integrated using Harmony to account for the batch effects.^116^ Cells were clustered using the Louvain algorithm at 0.4 resolution, resulting in 14 clusters. The first 30 PCs were used for Uniform Manifold Approximation and Projection (UMAP). Cells were annotated using the single nuclei dataset with Clustifyr package.^117^ Differential expression analysis was performed using the Dreamlet package.^103^ GSEA analysis was performed using the clusterProfiler package, using all genes with gseGO function to assess GO BP ontology, ranking all the genes by log_2_FC.^118–122^ Cell proportions were calculated by dividing the number of cells of specific identity with the total count of cells and compared between groups using the crumblr package. Neighbourhood analysis was performed using the CatsCradle^123^ package using the computeNBHDByCTMatrix, computeCellTypesPerCellTypeMatrix and computeNeighbourEnrichment function.

### FFPE human brain imaging

#### Histopathology imaging

Imaging of DAB-stained brain sections was accomplished using a slide scanner (Leica Aperio AT2) equipped with a NA 0.75 air objective resulting in a 0.25 µm image pixel size. Staining was done in accordance with standard procedure. The antibody used for αSyn staining was BD Transduction Laboratories™ Purified Mouse Anti-α-Synuclein (610787). The extent of pathology was quantified as the percentage of αSyn-positive area relative to total tissue area (“αSyn positivity”).

#### Sample preparation – FFPE human brain slices

Formalin-fixed paraffin-embedded (FFPE) tissue sections were obtained from the cingulate cortex (see Tables S1 and Data S1) and cut to 8 μm thickness. FFPE sections were baked at 37°C for 24 hours followed by 60 ℃ overnight. Sections were deparaffinized in xylene, and rehydrated using graded alcohols. Non-specific binding was blocked with 10% bovine serum albumin (BSA) solution in PBS for 30 minutes. The tissue was then pressure cooked in citrate buffer at pH 6 for 10 minutes. Tissue sections were incubated with primary antibodies; anti-phosphorated alpha-synuclein (**ab59264**, **AB_2270761, Abcam, 1:200**); and either ionized calcium-binding adapter molecule 1 **(GT10312, Thermo Fisher, AB_2735228, 1:200**) or Microtubule-Associated Protein 2 (**ab254143, Abcam, AB_2936822, 1:200**) for 1 hour at room temperature. The sections were then washed three times for five minutes in PBS followed by the corresponding AlexaFluor secondary antibodies (anti-rabbit 568, A11011, AB_143157, Thermo Fisher, anti-mouse 488, A11001, AB_2534069, Thermo Fisher, all at 1:200) for an additional hour at room temperature in the dark. Sections were then washed three times for five minutes again in PBS and incubated in Sudan Black (0.1% for 10 minutes, 199664-25G, Sigma Aldrich). Removal of Sudan Black occurred with three washes in 30% ethanol (E7148-500ML, Sigma Aldrich) before mounting with Vectashield+ (Vector Labs, H-1900) and coverslipping (VWR, 50 mm ✕ 24 mm #1 thickness, Catalogue Number 48404-453) for imaging. Sections were stored at 4°C until imaging was completed.

#### Optical setup

The microscope used to image the FFPE human brain slices was a spinning disk confocal microscope (3i intelligent imaging). The microscope was equipped with a 200 mW, 488 nm laser (LuxX) and a 150 mW, 561 nm laser (OBIS). These lasers were housed in a beam combiner (3i intelligent imaging), which focused them into an optical fibre which sent the illumination light into a field flattener (Yokogawa-Uniformizer for CSUW). The excitation light was then passed into a spinning disk unit (50 μm sized pinholes, Yokogawa CSU-W1 T2 Single Molecule Spinning Disk Confocal, SoRa Dual Microlens Disk) and then the microscope body (Zeiss Axio Observer 7 Basic Marianas™ Microscope with Definite Focus 3) using a dichroic mirror (FF01-440/521/607/700, Semrock). The fluorescence was filtered using either a FF01-525/45-25-STR filter (Semrock) in the case of 488 nm excitation or a FF02-617/73-25-STR filter (Semrock) in the case of 561 nm excitation. The fluorescence was then focused onto one of two sCMOS cameras (Prime 95B, Teledyne Photometrics). The objective lens was a Zeiss oil immersion objective (Alpha Plan-Apochromat 100x/1.46 NA Oil TIRF Objective, M27), and the microscope was controlled using a PC (Dell-Acquisition Workstation 310R) and software (Slidebook) provided by the manufacturer (3i intelligent imaging).

#### Camera gain calibration

To convert the pixel value to photons in a sCMOS camera, we recorded a series of image sequences at 7 different intensity levels (20,000 frames per intensity level) with uniform illumination, including one level at no illumination for the calculation of camera offset. For every pixel, the mean and variance were calculated across the 1000 frames, generating 7 different variance and mean values corresponding to the 7 non-zero illumination intensities. The camera offset per pixel was determined as the mean pixel value in the non-illuminated frame. The camera gain per pixel, expressed in photoelectrons per count, was determined by calculating the slope between the 7 variance and mean values per pixel, and subtracting the non-illuminated frame offset.^124^ Software available for this purpose is available at https://doi.org/10.5281/zenodo.10475643.

#### Brain imaging data analysis

Aggregate detection proceeded using the RASP pipeline described in Fu et al.^125^. To briefly summarise, images underwent a high-pass kernel, obtained through the difference between the original image and a Gaussian-blurred image (σ = 1.4px), followed by a Laplacian-of-Gaussian (LoG) kernel (σ = 2px). Puncta were selected as pixels in the top 95th percentile of brightness, and these were then accepted or rejected based on their integrated gradients and flatness.^125^

For cell mask detection, each 2D image was enhanced using a difference-of-Gaussian filter, with σ1 = 2px and σ2 = 60px. The image was then thresholded into a binary mask using a Yen threshold.^126^ This binary mask then had a binary opening operation applied to it with a disk morphology of radius 1 pixel, which was then followed by a binary closing operation with a disk morphology of radius 5 pixels. When all of the images in a volumetric stack had binary masks generated in this way, the scikit.image^127^ function binary_fill_holes was applied to the full 3D volume, after which small holes of <100 voxels were removed. Post this, 3D objects below a specified cell size were removed to end up with the final cell masks used for analysis.

### Human induced pluripotent stem cell (hiPSC) modelling

#### hiPSC culture

hiPSCs used in this study were derived from fibroblasts donated from healthy control and PD patients. All hiPSC line details are listed in Table S3. hiPSCs were maintained in feeder-free monolayers on Geltrex™ coated plates, were fed daily with mTeSR™1 medium, and were passaged using 0.5 µM EDTA. hiPSCs were maintained at 37°C and 5% CO2 and underwent regular mycoplasma testing and short tandem repeat profiling.

#### Differentiation of hiPSC-derived microglia-like cells (iMGLs)

hiPSC-derived microglia-like cell differentiation were performed using a modified published protocol.^46^ hiPSCs were dissociated with Accutase, collected in PBS, and were then centrifuged at 300 g for 3 minutes. hiPSCs were then resuspended in embryoid body differentiation media (EBD media). EBD media consisted of mTeSR ™1 medium supplemented with 10 µM ROCK inhibitor (Tocris), 50 ng/ml BMP-4, 50 ng/ml VEGF, and 20 ng/ml SCF (all PeproTech). Cells were plated into low adherence 96-well plates at 10,000 cells/well. The 96-well plates were then centrifuged at 800 rpm for 3 minutes before being transferred to a tissue culture incubator at 37°C and 5% CO_2_. After 48 hours, 50 µl of EBD media was added to each well. After a further 48 hours, embryoid bodies were collected from each well and were transferred into a 15 ml falcon using a P1000 pipette and wide-bore pipette tips. After aspirating any remaining EBD media from the 15 ml falcon, embryoid bodies were transferred to 75 cm^2^ flasks and were cultured in myeloid differentiation (MD) media. MD media consisted of X-vivo 15 medium (Lonza) supplemented with 1% Glutamax, 0.5% Penicillin-Streptomycin, and 50 µM 2-Mercaptoethanol, 100 ng/ml MCSF, and 25 ng/ml IL-3. Around 96 embryoid bodies were cultured in each 75 cm^2^ flask. Media was then refreshed every 3 days for 2 weeks. From this point cells were then harvested from the embryoid bodies weekly. 10 ml of media was taken from the flask and centrifuged at 300 g for 3 minutes. The resulting pellet was resuspended and filtered through a 40-micron filter. hiPSC-derived microglia-like cells were then plated and matured in X-Vivo 15 media supplemented with 1% Glutamax, 0.5% Penicillin-Streptomycin, and 100 ng/ml MCSF. All experiments were then performed >7 days post-harvest.

#### Immunocytochemistry

Cells were fixed in 4% paraformaldehyde and permeabilized with 0.2% Triton-X 100. 5% BSA was used to block non-specific binding before cells were incubated with primary antibodies overnight at 4°C. Cells were then washed three times with PBS and incubated with a secondary antibody for 1 hour at room temperature. Cells were mounted with an antifading medium after three further washes. Hoechst was included in the final wash. Imaging was performed on an Opera Phenix High-Content Screening System (PerkinElmer).

#### Human recombinant alpha-synuclein

Human recombinant monomeric αSyn was purified as previously described.^128^ For fluorescent experiments, αSyn monomers were labelled with maleimide-modified AF488 or AF594 (Invitrogen) via the cysteine thiol moiety.^129^ Unless otherwise stated cells were treated with 500 nM total monomeric αSyn. For Förster resonance energy transfer (FRET) experiments, an equimolar concentration of monomeric Alexa Fluor 488 (AF488)-labelled and Alexa Fluor 594 (AF594)-labelled αSyn was used. Human αSyn oligomers were obtained from StressMarq Biosciences (catalogue no. SPR-484). αSyn was low endotoxin (<5 EU/mL @ 2mg/mL). The equivalent concentration of LPS was added to the control/untreated condition in these experiments to account for any effect induced by residual endotoxin.

#### Live cell imaging - intracellular FRET

Either WT or A53T AF488-labelled and AF594-labelled αSyn were applied to iMGL for 24 hours. iMGL were washed twice and the media was replaced with HBSS. AF594-labelled αSyn was directly excited as a measure of total αSyn. As a measure of aggregated αSyn, AF488 was excited by 488 nm irradiation and emission was detected from AF594 (584-660 nm). FRET intensity was calculated by measuring the AF594 emission from the 488 nm irradiation in AF594-labelled αSyn positive regions.

#### Single-molecule confocal microscopy - FRET

Cell lysates of iMGL treated with AF488 and AF594 labelled αSyn were generated using mechanical dissociation. These were then analysed using single-molecule confocal microscopy with fast-flow microfluidics. The samples were diluted to 1 in 50 before being loaded into a 200-μl gel-loading tip (Life Technologies) attached to the inlet port of a microfluidic channel (25 μm in height, 100 μm in width and 1 cm in length) mounted onto the single-molecule confocal microscope. The confocal volume was focused 10 μm into the centre of the channel, and the solution was passed through the channel at an average velocity of 1 cm s^-1^ by applying a negative pressure, which was generated using a syringe pump attached to the outlet port via PTFE Tubing (1/32” inner diameter, 1/16” outer diameter; Darwin Microfluidics). After the appearance of single-molecule bursts corresponding to labelled αSyn passing through the confocal volume, the sample was measured for 300 seconds.

Single-molecule confocal microscopy measurements were performed on a custom built single-molecule confocal microscope described previously.^18,130^ A Gaussian laser beam at 488 nm (100 mW, LBX-488-100-CSB-OE, Oxxius) was collimated by a reflective collimator (RC08APC-P01, Thorlabs), and directed through the back-port of an inverted microscope (Nikon TE2000-U). A dichroic mirror (DI03-R405/488/561/635, Semrock) reflected the beam through an oil-immersion objective lens (Nikon CFI Plan Apochromat VC 100x Oil, NA 1.4, W.D 0.13 mm) that focused the light to a diffraction-limited confocal spot. The emitted fluorescence was collected by the same objective lens, and was passed through the same dichroic mirror, before being focused by a tube lens within the microscope body through a 50 μm pinhole (Thorlabs). A second dichroic mirror (585DRLP, Omega Filters) was used to separate the fluorescence from the two different fluorophores. The longer wavelength passed through the dichroic mirror and was focused by a lens (Plano apoconvex, focal length = 50 mm, Thorlabs) through a band-pass (FF01-629/53, Semrock) onto an Avalanche Photodiode (APD) detector (PerkinElmer). The shorter wavelengths were reflected and focused through a second filter set (long-pass: BLP01-488R-25 and band-pass: FF01-525/30-25, Semrock) onto a second APD. Outputs from the APDs were connected to a USB data acquisition card (USB-CTR04, Measurement Computing) controlled using previously published LabView software,^131^ which counted the signals and combined them into time-bins of 100 μs, the expected residence time of the oligomers in the confocal volume. The intensity of the 488 nm irradiation used was 2.94 mW at the back-port.

The data were analysed as in Choi et al.^18^ using custom-written scripts written in Python (https://doi.org/10.5281/zenodo.8189825). Coincident events were those that had at least 10 photon counts bin^-1^ in each channel. After accounting for autofluorescence and crosstalk (Equation 1-3), the FRET efficiency (Equation 1) and approximate size of each oligomer (Equation 4) was calculated from the intensities in each channel:

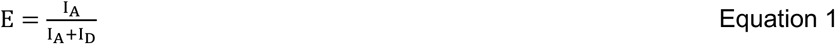

where I_D_ and I_A_ are the modified intensities in the donor and is acceptor channels with donor excitation only:

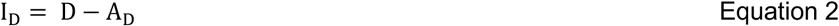

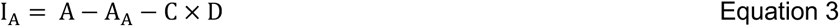

where D and A are the intensities from the donor and acceptor channels with only donor excitation only, respectively, A_D_ and A_A_ are the instrument-specific autofluorescence in the donor and acceptor channels, measured in the absence of fluorophores, and C is the instrument-specific cross-talk from the donor to acceptor channel. The cross-talk from acceptor to donor channel is negligible.

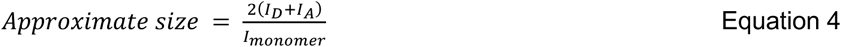

where *I_monomer_* is the average monomer brightness, calculated from the average of the non-coincident donor channel bursts intensities.

Due to the presence of long Stokes shift autofluorescence from microglia, it was only possible to quantify oligomers larger than 3 monomer units in size. The FRET histograms from this population had two obvious peaks: one centred at E = 0.26, corresponding to oligomers, and a second at E = 0.71, corresponding to autofluorescence. The histograms were globally fitted (shared x-centre and x-width for all time-points/treatments) to two Gaussian distributions to distinguish oligomers from autofluorescence, and the low-FRET peak integrated to determine the number of oligomers present in each sample.

#### Cytokine assays

To measure the levels of secreted cytokines, microglial cell culture media was collected 24 hours after treatments. The media was centrifuged for 10 minutes at 17,000 x g to remove cell debris. To measure a panel of cytokines the Meso Scale Diagnostics Pre-Coated V-Plex Proinflammatory Panel 1 (human) (MSD) was used according to the manufacturer’s instructions. Each sample was run in technical duplicate and cytokine concentrations were normalised to the total cell count within each well.

#### Single-cell RNA sequencing in iMGL

scRNAseq for iMGL was performed using the 10x Genomics Flex Single Cell Fixed RNA-seq kit. Samples were fixed in a 4% formaldehyde fixative solution, as described in the Demonstrated Protocols CG000478 and CG000553 and using Chromium Next GEM Single Cell Fixed RNA Sample Preparation Kit (10x Genomics PN-1000414). Gene expression was measured using barcoded probe pairs designed to hybridize to mRNA specifically. Using a microfluidic chip, the fixed and probe-hybridized single cell suspensions were partitioned into nanolitre-scale Gel Beads-in-emulsion (GEMs). A pool of ∼737,000 10x GEM Barcodes was sampled separately to index the contents of each partition. Inside the GEMs, probes were ligated and the 10x GEM Barcode was added, and all ligated probes within a GEM share a common 10x GEM Barcode. Barcoded and ligated probes were then pre-amplified in bulk, after which gene expression libraries were generated (User Guide: CG000527, Chromium Fixed RNA Profiling Reagent Kits for Multiplexed Samples) and sequenced on the NovaSeq 6000. Sequencing read configuration: 28-10-10-90.

Quality control metrics, including the number of detected genes, total read counts, and the percentage of mitochondrial gene reads, were calculated using the scuttle package (https://bioconductor.org/packages/3.22/bioc/html/scuttle.html). Outlier detection was performed for each sample independently, and cells failing these criteria were flagged. Doublets were identified using scDblFinder,^132^ and removed, along with QC-failed cells. Preliminary clustering revealed three clusters with aberrantly low or high total counts and detected genes, which were also excluded. Subsequent analysis was performed using Seurat.^113,114^ To account for donor-specific effects across the 6 iMGL lines, we applied Harmony integration.^116^ The number of principal components (PCs) used for downstream analyses was determined by cumulative variance explained, selecting the smallest number of PCs capturing ≥90% of total variance. Cells were clustered using the Louvain algorithm at a resolution of 0.2, yielding seven clusters.

Differential expression between clusters was performed on pseudobulked data across all samples using dreamletCompareClusters from the dreamlet package.^103^ Differential expression between genotypes for each cluster was performed on pseudobulked data using processAssays from dreamlet,^103^ controlling for sex. To explore functional enrichment of each cluster over-representation analysis was conducted using clusterProfiler (v4.10.1), with the compareCluster() function, considering pathways significant at FDR-adjusted *p* < 0.05. To reduce redundancy, GO terms were grouped by semantic similarity using GOSemSim with the “Wang” method and a threshold of 0.7. Pathway analysis for each cluster comparing *SNCA* A53T to control was performed using Gene Set Enrichment Analysis (GSEA) implemented in clusterProfiler, with the Gene Ontology (GO) database queried using ranked log₂FC effect sizes. To compare iMGL clusters with immune cell populations from a snRNAseq dataset, we used Clustifyr.^117^

#### Bulk RNA sequencing in iMGL

RNA was extracted using the Maxwell® RSC simplyRNA Cells kit (Promega) and the Maxwell® RSC instrument using the manufacturer’s protocols. Library construction was performed with either the NEBNext Ultra II Directional RNA Library Prep Kit (*SNCA* A53T experiment) or Watchmaker RNA with polyA enrichment (αSyn monomer and oligomer experiments) and sequencing was performed on the Illumina NovaSeq 6000 Sequencing system to generate 100 bp paired-end reads. Raw reads were processed using the RNA-seq nfcore pipeline (version 3.10.1). Briefly, the pipeline was executed with Nextflow version 23.04.3 where reads were trimmed using trimgalore and were subsequently aligned and quantified with STAR-RSEM against the human genome GRCh38 and annotation release 107. Differential gene expression analysis was performed in R using the DESeq2 package.^133^ The model used for the mutation paradigm (*SNCA* A53T) was: ∼individual + mutation + treatment + mutation:treatment. The model used for both the αSyn monomer and αSyn oligomer paradigms was: ∼ line + treatment. Differentially expressed genes were classified as differentially expressed if their Benjamini and Hochberg FDR adjusted p value was ≤ 0.05. Upregulated and downregulated genes were determined utilising their log_2_ fold change was > 0 and < 0 respectively.^89,133^. Briefly, the pipeline was executed with Nextflow version 23.04.3 where reads were trimmed using trimgalore and were subsequently aligned and quantified with STAR-RSEM against the human genome GRCh38 and annotation release 107. Differential gene expression analysis was performed in R using the DESeq2 package.^133^ The model used for the mutation paradigm (*SNCA* A53T) was: ∼individual + mutation + treatment + mutation:treatment. The model used for both the αSyn monomer and αSyn oligomer paradigms was: ∼ line + treatment. Differentially expressed genes were classified as differentially expressed if their Benjamini and Hochberg FDR adjusted p value was ≤ 0.05. Upregulated and downregulated genes were determined utilising their log_2_ fold change was > 0 and < 0 respectively.

The Gene Ontology (GO) database was interrogated for over-representation pathways analysis using clusterProfiler (v4.10.1). The compareCluster function was used to perform over-representation analyses. Pathways were significant if their FDR-adjusted p value was < 0.05. The number of GO pathways were reduced by semantic similarity using the GOSemSim R package. The “Wang” method and a similarity threshold for grouping terms of 0.7 were used. The reduction was performed for each experimental paradigm in iMGL separately.

## ACKNOWLEDGMENTS

JRE, MGP, JSB, JCB, CET, AFB, JWB, JL, AP, EEB, BF, LN, HL, KD, MV, NWW, SFL, MR, SG were funded by Aligning Science Across Parkinson’s [Grant numbers: ASAP-000478 and ASAP-000509] through the Michael J. Fox Foundation for Parkinson’s Research (MJFF). SG is supported by the MRC (MR/T008199/1), and acknowledges funding from the UCLH Biomedical Research Centre. The single-molecule confocal microscope used in this study was funded by Alzheimer’s Research UK (ARUK-EG2018B-004), and a kind donation from Dr Jim Love. R.F. was supported by an EPSRC CASE studentship with UCB Bio-pharmaceuticals. R.S.S. was supported by an MND Association Lady Edith Wolfson Junior Non-Clinical Research Fellowship Saleeb/Oct22/980-799.

## SUPPLEMENTAL INFORMATION

**Document S1. Figures S1–S21, and Tables S1-S3**

**Data S1. Experimental details for control and Braak 3–4 PD cohort**

**Data S2. Bulk RNA-seq (Braak 3–4 PD) differential gene expression analysis.**

**Data S3. Bulk RNA-seq (Braak 3–4 PD) pathway analysis.**

**Data S4. snRNAseq (Braak 3–4 PD) cell type proportion.**

**Data S5. snRNAseq (Braak 3–4 PD) differential gene expression analysis.**

**Data S6. snRNAseq (Braak 3–4 PD) pathway analysis.**

**Data S7. snRNAseq (Braak 3–4 PD) GWAS enrichment overlap.**

**Data S8. snRNAseq (Braak 3–4 PD) GWAS enrichment significance.**

**Data S9. CosMx (Braak 3–4 PD ACC) INC1 differential gene expression analysis.**

**Data S10. scRNAseq (*SNCA* A53T iMGL) differential gene expression analysis.**

**Data S11. scRNAseq (*SNCA* A53T iMGL) pathway analysis.**

**Data S12. Bulk RNA-seq (*SNCA* A53T iMGL) differential gene expression analysis.**

**Data S13. Bulk RNA-seq (*SNCA* A53T iMGL) pathway analysis.**

**Data S14. Bulk RNA-seq (iMGL monomer treatment) differential gene expression analysis.**

**Data S15. Bulk RNA-seq (iMGL monomer treatment) pathway analysis.**

**Data S16. Bulk RNA-seq (iMGL oligomer treatment) differential gene expression analysis.**

**Data S17. Bulk RNA-seq (iMGL oligomer treatment) pathway analysis.**

## REFERENCES

1. Poewe, W., Seppi, K., Tanner, C.M., Halliday, G.M., Brundin, P., Volkmann, J., Schrag, A.-E., and Lang, A.E. (2017). Parkinson disease. Nat. Rev. Dis. Primers 3, 17013. 10.1038/nrdp.2017.13.

2. Goedert, M., Spillantini, M.G., Del Tredici, K., and Braak, H. (2013). 100 years of Lewy pathology. Nat. Rev. Neurol. 9, 13–24. 10.1038/nrneurol.2012.242.

3. 3. Braak, H., Del Tredici, K., Rüb, U., de Vos, R.A.I., Jansen Steur, E.N.H., and Braak, E. (2003). Staging of brain pathology related to sporadic Parkinson’s disease. Neurobiol. Aging 24, 197–211. 10.1016/s0197-4580(02)00065-9.

4. Borrageiro, G., Haylett, W., Seedat, S., Kuivaniemi, H., and Bardien, S. (2018). A review of genome-wide transcriptomics studies in Parkinson’s disease. Eur. J. Neurosci. 47, 1–16. 10.1111/ejn.13760.

5. Smajić, S., Prada-Medina, C.A., Landoulsi, Z., Ghelfi, J., Delcambre, S., Dietrich, C., Jarazo, J., Henck, J., Balachandran, S., Pachchek, S., et al. (2022). Single-cell sequencing of human midbrain reveals glial activation and a Parkinson-specific neuronal state. Brain 145, 964–978. 10.1093/brain/awab446.

6. Martirosyan, A., Ansari, R., Pestana, F., Hebestreit, K., Gasparyan, H., Aleksanyan, R., Hnatova, S., Poovathingal, S., Marneffe, C., Thal, D.R., et al. (2024). Unravelling cell type-specific responses to Parkinson’s Disease at single cell resolution. Mol. Neurodegener. 19, 7. 10.1186/s13024-023-00699-0.

7. Booth, H.D.E., Hirst, W.D., and Wade-Martins, R. (2017). The role of astrocyte dysfunction in parkinson’s disease pathogenesis. Trends Neurosci. 40, 358–370. 10.1016/j.tins.2017.04.001.

8. Isik, S., Yeman Kiyak, B., Akbayir, R., Seyhali, R., and Arpaci, T. (2023). Microglia mediated neuroinflammation in parkinson’s disease. Cells 12. 10.3390/cells12071012.

9. Reynolds, R.H., Botía, J., Nalls, M.A., International Parkinson’s Disease Genomics Consortium (IPDGC), System Genomics of Parkinson’s Disease (SGPD), Hardy, J., Gagliano Taliun, S.A., and Ryten, M. (2019). Moving beyond neurons: the role of cell type-specific gene regulation in Parkinson’s disease heritability. npj Parkinsons Disease 5, 6. 10.1038/s41531-019-0076-6.

10. Bryois, J., Skene, N.G., Hansen, T.F., Kogelman, L.J.A., Watson, H.J., Liu, Z., Eating Disorders Working Group of the Psychiatric Genomics Consortium, International Headache Genetics Consortium, 23andMe Research Team, Brueggeman, L., et al. (2020). Genetic identification of cell types underlying brain complex traits yields insights into the etiology of Parkinson’s disease. Nat. Genet. 52, 482–493. 10.1038/s41588-020-0610-9.

11. Deyell, J.S., Sriparna, M., Ying, M., and Mao, X. (2023). The Interplay between α-Synuclein and Microglia in α-Synucleinopathies. Int. J. Mol. Sci. *24*. 10.3390/ijms24032477.

12. Thi Lai, T., Kim, Y.E., Nguyen, L.T.N., Thi Nguyen, T., Kwak, I.H., Richter, F., Kim, Y.J., and Ma, H.-I. (2024). Microglial inhibition alleviates alpha-synuclein propagation and neurodegeneration in Parkinson’s disease mouse model. npj Parkinsons Disease 10, 32. 10.1038/s41531-024-00640-2.

13. Bido, S., Muggeo, S., Massimino, L., Marzi, M.J., Giannelli, S.G., Melacini, E., Nannoni, M., Gambarè, D., Bellini, E., Ordazzo, G., et al. (2021). Microglia-specific overexpression of α-synuclein leads to severe dopaminergic neurodegeneration by phagocytic exhaustion and oxidative toxicity. Nat. Commun. 12, 6237. 10.1038/s41467-021-26519-x.

14. Chartier, S., and Duyckaerts, C. (2018). Is Lewy pathology in the human nervous system chiefly an indicator of neuronal protection or of toxicity? Cell Tissue Res. 373, 149–160. 10.1007/s00441-018-2854-6.

15. Bengoa-Vergniory, N., Roberts, R.F., Wade-Martins, R., and Alegre-Abarrategui, J. (2017). Alpha-synuclein oligomers: a new hope. Acta Neuropathol. 134, 819–838. 10.1007/s00401-017-1755-1.

16. Choi, M.L., and Gandhi, S. (2018). Crucial role of protein oligomerization in the pathogenesis of Alzheimer’s and Parkinson’s diseases. FEBS J. 285, 3631–3644. 10.1111/febs.14587.

17. Musteikytė, G., Jayaram, A.K., Xu, C.K., Vendruscolo, M., Krainer, G., and Knowles, T.P.J. (2021). Interactions of α-synuclein oligomers with lipid membranes. Biochim. Biophys. Acta Biomembr. 1863, 183536. 10.1016/j.bbamem.2020.183536.

18. Choi, M.L., Chappard, A., Singh, B.P., Maclachlan, C., Rodrigues, M., Fedotova, E.I., Berezhnov, A.V., De, S., Peddie, C.J., Athauda, D., et al. (2022). Pathological structural conversion of α-synuclein at the mitochondria induces neuronal toxicity. Nat. Neurosci. 25, 1134–1148. 10.1038/s41593-022-01140-3.

19. Ludtmann, M.H.R., Angelova, P.R., Horrocks, M.H., Choi, M.L., Rodrigues, M., Baev, A.Y., Berezhnov, A.V., Yao, Z., Little, D., Banushi, B., et al. (2018). α-synuclein oligomers interact with ATP synthase and open the permeability transition pore in Parkinson’s disease. Nat. Commun. 9, 2293. 10.1038/s41467-018-04422-2.

20. Danzer, K.M., Haasen, D., Karow, A.R., Moussaud, S., Habeck, M., Giese, A., Kretzschmar, H., Hengerer, B., and Kostka, M. (2007). Different species of alpha-synuclein oligomers induce calcium influx and seeding. J. Neurosci. 27, 9220–9232. 10.1523/JNEUROSCI.2617-07.2007.

21. Leandrou, E., Chalatsa, I., Anagnostou, D., Machalia, C., Semitekolou, M., Filippa, V., Makridakis, M., Vlahou, A., Anastasiadou, E., Vekrellis, K., et al. (2024). α-Synuclein oligomers potentiate neuroinflammatory NF-κB activity and induce Cav3.2 calcium signaling in astrocytes. Transl. Neurodegener. 13, 11. 10.1186/s40035-024-00401-4.

22. Rockenstein, E., Nuber, S., Overk, C.R., Ubhi, K., Mante, M., Patrick, C., Adame, A., Trejo-Morales, M., Gerez, J., Picotti, P., et al. (2014). Accumulation of oligomer-prone α-synuclein exacerbates synaptic and neuronal degeneration in vivo. Brain 137, 1496–1513. 10.1093/brain/awu057.

23. D’Sa, K., Choi, M.L., Wagen, A.Z., Setó-Salvia, N., Kopach, O., Evans, J.R., Rodrigues, M., Lopez-Garcia, P., Lachica, J., Clarke, B.E., et al. (2025). Astrocytic RNA editing regulates the host immune response to alpha-synuclein. Sci. Adv. 11, eadp8504. 10.1126/sciadv.adp8504.

24. Andrews, R., Fu, B., Toomey, C.E., Breiter, J.C., Lachica, J., Tian, R., Beckwith, J.S., Needham, L.-M., Chant, G.J., Loiseau, C., et al. (2024). Large-scale visualisation of α-synuclein oligomers in Parkinson’s disease brain tissue. BioRxiv. 10.1101/2024.02.17.580698.

25. Toomey, C.E., Heywood, W.E., Evans, J.R., Lachica, J., Pressey, S.N., Foti, S.C., Al Shahrani, M., D’Sa, K., Hargreaves, I.P., Heales, S., et al. (2022). Mitochondrial dysfunction is a key pathological driver of early stage Parkinson’s. Acta Neuropathol. Commun. 10, 134. 10.1186/s40478-022-01424-6.

26. Feleke, R., Reynolds, R.H., Smith, A.M., Tilley, B., Taliun, S.A.G., Hardy, J., Matthews, P.M., Gentleman, S., Owen, D.R., Johnson, M.R., et al. (2021). Cross-platform transcriptional profiling identifies common and distinct molecular pathologies in Lewy body diseases. Acta Neuropathol. 142, 449–474. 10.1007/s00401-021-02343-x.

27. Nido, G.S., Dick, F., Toker, L., Petersen, K., Alves, G., Tysnes, O.-B., Jonassen, I., Haugarvoll, K., and Tzoulis, C. (2020). Common gene expression signatures in Parkinson’s disease are driven by changes in cell composition. Acta Neuropathol. Commun. 8, 55. 10.1186/s40478-020-00932-7.

28. Srinivasan, K., Friedman, B.A., Etxeberria, A., Huntley, M.A., van der Brug, M.P., Foreman, O., Paw, J.S., Modrusan, Z., Beach, T.G., Serrano, G.E., et al. (2020). Alzheimer’s patient microglia exhibit enhanced aging and unique transcriptional activation. Cell Rep. 31, 107843. 10.1016/j.celrep.2020.107843.

29. Keren-Shaul, H., Spinrad, A., Weiner, A., Matcovitch-Natan, O., Dvir-Szternfeld, R., Ulland, T.K., David, E., Baruch, K., Lara-Astaiso, D., Toth, B., et al. (2017). A Unique Microglia Type Associated with Restricting Development of Alzheimer’s Disease. Cell 169, 1276–1290.e17. 10.1016/j.cell.2017.05.018.

30. Martins-Ferreira, R., Calafell-Segura, J., Leal, B., Rodríguez-Ubreva, J., Martínez-Saez, E., Mereu, E., Pinho E Costa, P., Laguna, A., and Ballestar, E. (2025). The Human Microglia Atlas (HuMicA) unravels changes in disease-associated microglia subsets across neurodegenerative conditions. Nat. Commun. 16, 739. 10.1038/s41467-025-56124-1.

31. Roodveldt, C., Bernardino, L., Oztop-Cakmak, O., Dragic, M., Fladmark, K.E., Ertan, S., Aktas, B., Pita, C., Ciglar, L., Garraux, G., et al. (2024). The immune system in Parkinson’s disease: what we know so far. Brain 147, 3306–3324. 10.1093/brain/awae177.

32. Choi, B.-K., Choi, M.-G., Kim, J.-Y., Yang, Y., Lai, Y., Kweon, D.-H., Lee, N.K., and Shin, Y.-K. (2013). Large α-synuclein oligomers inhibit neuronal SNARE-mediated vesicle docking. Proc Natl Acad Sci USA 110, 4087–4092. 10.1073/pnas.1218424110.

33. Burré, J., Sharma, M., and Südhof, T.C. (2014). α-Synuclein assembles into higher-order multimers upon membrane binding to promote SNARE complex formation. Proc Natl Acad Sci USA 111, E4274–83. 10.1073/pnas.1416598111.

34. Parra-Rivas, L.A., Madhivanan, K., Aulston, B.D., Wang, L., Prakashchand, D.D., Boyer, N.P., Saia-Cereda, V.M., Branes-Guerrero, K., Pizzo, D.P., Bagchi, P., et al. (2023). Serine-129 phosphorylation of α-synuclein is an activity-dependent trigger for physiologic protein-protein interactions and synaptic function. Neuron 111, 4006–4023.e10. 10.1016/j.neuron.2023.11.020.

35. Davalos, D., Grutzendler, J., Yang, G., Kim, J.V., Zuo, Y., Jung, S., Littman, D.R., Dustin, M.L., and Gan, W.-B. (2005). ATP mediates rapid microglial response to local brain injury in vivo. Nat. Neurosci. 8, 752–758. 10.1038/nn1472.

36. Baligács, N., Albertini, G., Borrie, S.C., Serneels, L., Pridans, C., Balusu, S., and De Strooper, B. (2024). Homeostatic microglia initially seed and activated microglia later reshape amyloid plaques in Alzheimer’s Disease. Nat. Commun. 15, 10634. 10.1038/s41467-024-54779-w.

37. Bouvier, D.S., Jones, E.V., Quesseveur, G., Davoli, M.A., A Ferreira, T., Quirion, R., Mechawar, N., and Murai, K.K. (2016). High resolution dissection of reactive glial nets in alzheimer’s disease. Sci. Rep. 6, 24544. 10.1038/srep24544.

38. Savage, J.C., Carrier, M., and Tremblay, M.-È. (2019). Morphology of microglia across contexts of health and disease. Methods Mol. Biol. 2034, 13–26. 10.1007/978-1-4939-9658-2_2.

39. O’Banion, M.K. (1999). Cyclooxygenase-2: molecular biology, pharmacology, and neurobiology. Crit. Rev. Neurobiol. 13, 45–82.

40. Corder, E.H., Saunders, A.M., Strittmatter, W.J., Schmechel, D.E., Gaskell, P.C., Small, G.W., Roses, A.D., Haines, J.L., and Pericak-Vance, M.A. (1993). Gene dose of apolipoprotein E type 4 allele and the risk of Alzheimer’s disease in late onset families. Science 261, 921–923. 10.1126/science.8346443.

41. Nalls, M.A., Blauwendraat, C., Vallerga, C.L., Heilbron, K., Bandres-Ciga, S., Chang, D., Tan, M., Kia, D.A., Noyce, A.J., Xue, A., et al. (2019). Identification of novel risk loci, causal insights, and heritable risk for Parkinson’s disease: a meta-analysis of genome-wide association studies. Lancet Neurol. 18, 1091–1102. 10.1016/S1474-4422(19)30320-5.

42. Kia, D.A., Zhang, D., Guelfi, S., Manzoni, C., Hubbard, L., Reynolds, R.H., Botía, J., Ryten, M., Ferrari, R., Lewis, P.A., et al. (2021). Identification of Candidate Parkinson Disease Genes by Integrating Genome-Wide Association Study, Expression, and Epigenetic Data Sets. JAMA Neurol. 78, 464–472. 10.1001/jamaneurol.2020.5257.

43. Tan, M.M.X., Lawton, M.A., Pollard, M.I., Brown, E., Real, R., Carrasco, A.M., Bekadar, S., Jabbari, E., Reynolds, R.H., Iwaki, H., et al. (2024). Genome-wide determinants of mortality and motor progression in Parkinson’s disease. npj Parkinsons Disease 10, 113. 10.1038/s41531-024-00729-8.

44. Tan, M.M.X., Lawton, M.A., Jabbari, E., Reynolds, R.H., Iwaki, H., Blauwendraat, C., Kanavou, S., Pollard, M.I., Hubbard, L., Malek, N., et al. (2021). Genome-Wide Association Studies of Cognitive and Motor Progression in Parkinson’s Disease. Mov. Disord. 36, 424–433. 10.1002/mds.28342.

45. Krasemann, S., Madore, C., Cialic, R., Baufeld, C., Calcagno, N., El Fatimy, R., Beckers, L., O’Loughlin, E., Xu, Y., Fanek, Z., et al. (2017). The TREM2-APOE pathway drives the transcriptional phenotype of dysfunctional microglia in neurodegenerative diseases. Immunity 47, 566–581.e9. 10.1016/j.immuni.2017.08.008.

46. Garcia-Reitboeck, P., Phillips, A., Piers, T.M., Villegas-Llerena, C., Butler, M., Mallach, A., Rodrigues, C., Arber, C.E., Heslegrave, A., Zetterberg, H., et al. (2018). Human Induced Pluripotent Stem Cell-Derived Microglia-Like Cells Harboring TREM2 Missense Mutations Show Specific Deficits in Phagocytosis. Cell Rep. 24, 2300–2311. 10.1016/j.celrep.2018.07.094.

47. Polymeropoulos, M.H., Lavedan, C., Leroy, E., Ide, S.E., Dehejia, A., Dutra, A., Pike, B., Root, H., Rubenstein, J., Boyer, R., et al. (1997). Mutation in the alpha-synuclein gene identified in families with Parkinson’s disease. Science 276, 2045–2047. 10.1126/science.276.5321.2045.

48. Iljina, M., Garcia, G.A., Horrocks, M.H., Tosatto, L., Choi, M.L., Ganzinger, K.A., Abramov, A.Y., Gandhi, S., Wood, N.W., Cremades, N., et al. (2016). Kinetic model of the aggregation of alpha-synuclein provides insights into prion-like spreading. Proc Natl Acad Sci USA 113, E1206–15. 10.1073/pnas.1524128113.

49. Tosatto, L., Horrocks, M.H., Dear, A.J., Knowles, T.P.J., Dalla Serra, M., Cremades, N., Dobson, C.M., and Klenerman, D. (2015). Single-molecule FRET studies on alpha-synuclein oligomerization of Parkinson’s disease genetically related mutants. Sci. Rep. 5, 16696. 10.1038/srep16696.

50. Virdi, G.S., Choi, M.L., Evans, J.R., Yao, Z., Athauda, D., Strohbuecker, S., Nirujogi, R.S., Wernick, A.I., Pelegrina-Hidalgo, N., Leighton, C., et al. (2022). Protein aggregation and calcium dysregulation are hallmarks of familial Parkinson’s disease in midbrain dopaminergic neurons. npj Parkinsons Disease 8, 162. 10.1038/s41531-022-00423-7.

51. Liu, W., Taso, O., Wang, R., Bayram, S., Graham, A.C., Garcia-Reitboeck, P., Mallach, A., Andrews, W.D., Piers, T.M., Botia, J.A., et al. (2020). Trem2 promotes anti-inflammatory responses in microglia and is suppressed under pro-inflammatory conditions. Hum. Mol. Genet. 29, 3224–3248. 10.1093/hmg/ddaa209.

52. Dolan, M.-J., Therrien, M., Jereb, S., Kamath, T., Gazestani, V., Atkeson, T., Marsh, S.E., Goeva, A., Lojek, N.M., Murphy, S., et al. (2023). Exposure of iPSC-derived human microglia to brain substrates enables the generation and manipulation of diverse transcriptional states in vitro. Nat. Immunol. 24, 1382–1390. 10.1038/s41590-023-01558-2.

53. Shen, L., Sigal, L.J., Boes, M., and Rock, K.L. (2004). Important role of cathepsin S in generating peptides for TAP-independent MHC class I crosspresentation in vivo. Immunity 21, 155–165. 10.1016/j.immuni.2004.07.004.

54. Zhao, K., Sun, Y., Zhong, S., and Luo, J.-L. (2024). The multifaceted roles of cathepsins in immune and inflammatory responses: implications for cancer therapy, autoimmune diseases, and infectious diseases. Biomark. Res. 12, 165. 10.1186/s40364-024-00711-9.

55. Embgenbroich, M., and Burgdorf, S. (2018). Current Concepts of Antigen Cross-Presentation. Front. Immunol. 9, 1643. 10.3389/fimmu.2018.01643.

56. Drobny, A., Prieto Huarcaya, S., Dobert, J., Kluge, A., Bunk, J., Schlothauer, T., and Zunke, F. (2022). The role of lysosomal cathepsins in neurodegeneration: Mechanistic insights, diagnostic potential and therapeutic approaches. Biochim. Biophys. Acta Mol. Cell Res. 1869, 119243. 10.1016/j.bbamcr.2022.119243.

57. Nalls, M.A., Blauwendraat, C., Sargent, L., Vitale, D., Leonard, H., Iwaki, H., Song, Y., Bandres-Ciga, S., Menden, K., Faghri, F., et al. (2021). Evidence for GRN connecting multiple neurodegenerative diseases. Brain Commun. 3, fcab095. 10.1093/braincomms/fcab095.

58. He, Y., Kaya, I., Shariatgorji, R., Lundkvist, J., Wahlberg, L.U., Nilsson, A., Mamula, D., Kehr, J., Zareba-Paslawska, J., Biverstål, H., et al. (2023). Prosaposin maintains lipid homeostasis in dopamine neurons and counteracts experimental parkinsonism in rodents. Nat. Commun. 14, 5804. 10.1038/s41467-023-41539-5.

59. Diaz-Ortiz, M.E., Seo, Y., Posavi, M., Carceles Cordon, M., Clark, E., Jain, N., Charan, R., Gallagher, M.D., Unger, T.L., Amari, N., et al. (2022). GPNMB confers risk for Parkinson’s disease through interaction with α-synuclein. Science 377, eabk0637. 10.1126/science.abk0637.

60. Zhou, S., Li, T., Zhang, W., Wu, J., Hong, H., Quan, W., Qiao, X., Cui, C., Qiao, C., Zhao, W., et al. (2025). The cGAS-STING-interferon regulatory factor 7 pathway regulates neuroinflammation in Parkinson’s disease. Neural Regen. Res. 20, 2361–2372. 10.4103/NRR.NRR-D-23-01684.

61. Huang, H., Li, H., Chen, X., Yang, Y., Li, X., Li, W., Huang, C., Meng, X., Zhang, L., and Li, J. (2017). HMGA2, a driver of inflammation, is associated with hypermethylation in acute liver injury. Toxicol. Appl. Pharmacol. 328, 34–45. 10.1016/j.taap.2017.05.005.

62. Majhi, S., Roy, P., Jo, M., Liu, J., Hurto, R., Freddolino, L., and Marsh, E.N.G. (2025). Viperin expression leads to downregulation of mitochondrial genes through misincorporation of ddhCTP by mitochondrial RNA polymerase. J. Biol. Chem. 301, 108359. 10.1016/j.jbc.2025.108359.

63. Dumbrepatil, A.B., Zegalia, K.A., Sajja, K., Kennedy, R.T., and Marsh, E.N.G. (2020). Targeting viperin to the mitochondrion inhibits the thiolase activity of the trifunctional enzyme complex. J. Biol. Chem. 295, 2839–2849. 10.1074/jbc.RA119.011526.

64. Shan, B., Vazquez, E., and Lewis, J.A. (1990). Interferon selectively inhibits the expression of mitochondrial genes: a novel pathway for interferon-mediated responses. EMBO J. 9, 4307–4314. 10.1002/j.1460-2075.1990.tb07879.x.

65. Lewis, J.A., Huq, A., and Najarro, P. (1996). Inhibition of mitochondrial function by interferon. J. Biol. Chem. 271, 13184–13190. 10.1074/jbc.271.22.13184.

66. Kam, T.-I., Hinkle, J.T., Dawson, T.M., and Dawson, V.L. (2020). Microglia and astrocyte dysfunction in parkinson’s disease. Neurobiol. Dis. 144, 105028. 10.1016/j.nbd.2020.105028.

67. Arcuri, C., Mecca, C., Bianchi, R., Giambanco, I., and Donato, R. (2017). The pathophysiological role of microglia in dynamic surveillance, phagocytosis and structural remodeling of the developing CNS. Front. Mol. Neurosci. 10, 191. 10.3389/fnmol.2017.00191.

68. Gao, C., Jiang, J., Tan, Y., and Chen, S. (2023). Microglia in neurodegenerative diseases: mechanism and potential therapeutic targets. Signal Transduct. Target. Ther. 8, 359. 10.1038/s41392-023-01588-0.

69. Le, W., Wu, J., and Tang, Y. (2016). Protective microglia and their regulation in parkinson’s disease. Front. Mol. Neurosci. 9, 89. 10.3389/fnmol.2016.00089.

70. Badanjak, K., Fixemer, S., Smajić, S., Skupin, A., and Grünewald, A. (2021). The contribution of microglia to neuroinflammation in parkinson’s disease. Int. J. Mol. Sci. 22. 10.3390/ijms22094676.

71. McGeer, P.L., Itagaki, S., Boyes, B.E., and McGeer, E.G. (1988). Reactive microglia are positive for HLA-DR in the substantia nigra of Parkinson’s and Alzheimer’s disease brains. Neurology 38, 1285–1291. 10.1212/wnl.38.8.1285.

72. Imamura, K., Hishikawa, N., Sawada, M., Nagatsu, T., Yoshida, M., and Hashizume, Y. (2003). Distribution of major histocompatibility complex class II-positive microglia and cytokine profile of Parkinson’s disease brains. Acta Neuropathol. 106, 518–526. 10.1007/s00401-003-0766-2.

73. Gerhard, A., Pavese, N., Hotton, G., Turkheimer, F., Es, M., Hammers, A., Eggert, K., Oertel, W., Banati, R.B., and Brooks, D.J. (2006). In vivo imaging of microglial activation with [11C](R)-PK11195 PET in idiopathic Parkinson’s disease. Neurobiol. Dis. 21, 404–412. 10.1016/j.nbd.2005.08.002.

74. Rojanathammanee, L., Murphy, E.J., and Combs, C.K. (2011). Expression of mutant alpha-synuclein modulates microglial phenotype in vitro. J. Neuroinflammation 8, 44. 10.1186/1742-2094-8-44.

75. Gordon, R., Albornoz, E.A., Christie, D.C., Langley, M.R., Kumar, V., Mantovani, S., Robertson, A.A.B., Butler, M.S., Rowe, D.B., O’Neill, L.A., et al. (2018). Inflammasome inhibition prevents α-synuclein pathology and dopaminergic neurodegeneration in mice. Sci. Transl. Med. 10. 10.1126/scitranslmed.aah4066.

76. Krzisch, M., Yuan, B., Chen, W., Osaki, T., Fu, D., Garrett-Engele, C.M., Svoboda, D.S., Andrykovich, K.R., Gallagher, M.D., Sur, M., et al. (2025). The A53T Mutation in α-Synuclein Enhances Proinflammatory Activation in Human Microglia Upon Inflammatory Stimulus. Biol. Psychiatry 97, 730–742. 10.1016/j.biopsych.2024.07.011.

77. Sochocka, M., Diniz, B.S., and Leszek, J. (2017). Inflammatory response in the CNS: friend or foe? Mol. Neurobiol. 54, 8071–8089. 10.1007/s12035-016-0297-1.

78. Borst, K., Dumas, A.A., and Prinz, M. (2021). Microglia: Immune and non-immune functions. Immunity 54, 2194–2208. 10.1016/j.immuni.2021.09.014.

79. Szepesi, Z., Manouchehrian, O., Bachiller, S., and Deierborg, T. (2018). Bidirectional Microglia-Neuron Communication in Health and Disease. Front. Cell. Neurosci. 12, 323. 10.3389/fncel.2018.00323.

80. Duke, T., and Graham, I. (2009). Equilibrium mechanisms of receptor clustering. Prog. Biophys. Mol. Biol. 100, 18–24. 10.1016/j.pbiomolbio.2009.08.003.

81. Lively, S., and Schlichter, L.C. (2018). Microglia Responses to Pro-inflammatory Stimuli (LPS, IFNγ+TNFα) and Reprogramming by Resolving Cytokines (IL-4, IL-10). Front. Cell. Neurosci. 12, 215. 10.3389/fncel.2018.00215.

82. Litteljohn, D., Mangano, E., Clarke, M., Bobyn, J., Moloney, K., and Hayley, S. (2010). Inflammatory mechanisms of neurodegeneration in toxin-based models of Parkinson’s disease. Parkinsons Dis 2011, 713517. 10.4061/2011/713517.

83. Fellner, L., Irschick, R., Schanda, K., Reindl, M., Klimaschewski, L., Poewe, W., Wenning, G.K., and Stefanova, N. (2013). Toll-like receptor 4 is required for α-synuclein dependent activation of microglia and astroglia. Glia 61, 349–360. 10.1002/glia.22437.

84. Kim, C., Ho, D.-H., Suk, J.-E., You, S., Michael, S., Kang, J., Joong Lee, S., Masliah, E., Hwang, D., Lee, H.-J., et al. (2013). Neuron-released oligomeric α-synuclein is an endogenous agonist of TLR2 for paracrine activation of microglia. Nat. Commun. 4, 1562. 10.1038/ncomms2534.

85. Bhatia, D., Grozdanov, V., Ruf, W.P., Kassubek, J., Ludolph, A.C., Weishaupt, J.H., and Danzer, K.M. (2021). T-cell dysregulation is associated with disease severity in Parkinson’s Disease. J. Neuroinflammation 18, 250. 10.1186/s12974-021-02296-8.

86. Williams, G.P., Schonhoff, A.M., Jurkuvenaite, A., Gallups, N.J., Standaert, D.G., and Harms, A.S. (2021). CD4 T cells mediate brain inflammation and neurodegeneration in a mouse model of Parkinson’s disease. Brain 144, 2047–2059. 10.1093/brain/awab103.

87. Contaldi, E., Magistrelli, L., and Comi, C. (2022). T lymphocytes in parkinson’s disease. J Parkinsons Dis 12, S65–S74. 10.3233/JPD-223152.

88. Lind-Holm Mogensen, F., Seibler, P., Grünewald, A., and Michelucci, A. (2025). Microglial dynamics and neuroinflammation in prodromal and early Parkinson’s disease. J. Neuroinflammation 22, 136. 10.1186/s12974-025-03462-y.

89. Di Tommaso, P., Chatzou, M., Floden, E.W., Barja, P.P., Palumbo, E., and Notredame, C. (2017). Nextflow enables reproducible computational workflows. Nat. Biotechnol. 35, 316–319. 10.1038/nbt.3820.

90. Chen, S., Zhou, Y., Chen, Y., and Gu, J. (2018). fastp: an ultra-fast all-in-one FASTQ preprocessor. Bioinformatics 34, i884–i890. 10.1093/bioinformatics/bty560.

91. Patro, R., Duggal, G., Love, M.I., Irizarry, R.A., and Kingsford, C. (2017). Salmon provides fast and bias-aware quantification of transcript expression. Nat. Methods 14, 417–419. 10.1038/nmeth.4197.

92. Wang, L., Wang, S., and Li, W. (2012). RSeQC: quality control of RNA-seq experiments. Bioinformatics 28, 2184–2185. 10.1093/bioinformatics/bts356.

93. Okonechnikov, K., Conesa, A., and García-Alcalde, F. (2016). Qualimap 2: advanced multi-sample quality control for high-throughput sequencing data. Bioinformatics 32, 292–294. 10.1093/bioinformatics/btv566.

94. Ewels, P., Magnusson, M., Lundin, S., and Käller, M. (2016). MultiQC: summarize analysis results for multiple tools and samples in a single report. Bioinformatics 32, 3047–3048. 10.1093/bioinformatics/btw354.

95. Skene, N.G., and Grant, S.G.N. (2016). Identification of vulnerable cell types in major brain disorders using single cell transcriptomes and expression weighted cell type enrichment. Front. Neurosci. 10, 16. 10.3389/fnins.2016.00016.

96. Dyer, S.C., Austine-Orimoloye, O., Azov, A.G., Barba, M., Barnes, I., Barrera-Enriquez, V.P., Becker, A., Bennett, R., Beracochea, M., Berry, A., et al. (2025). Ensembl 2025. Nucleic Acids Res. 53, D948–D957. 10.1093/nar/gkae1071.

97. Curion, F., Rich-Griffin, C., Agarwal, D., Ouologuem, S., Rue-Albrecht, K., May, L., Garcia, G.E.L., Heumos, L., Thomas, T., Lason, W., et al. (2024). Panpipes: a pipeline for multiomic single-cell and spatial transcriptomic data analysis. Genome Biol. 25, 181. 10.1186/s13059-024-03322-7.

98. Jäkel, S., Agirre, E., Mendanha Falcão, A., van Bruggen, D., Lee, K.W., Knuesel, I., Malhotra, D., Ffrench-Constant, C., Williams, A., and Castelo-Branco, G. (2019). Altered human oligodendrocyte heterogeneity in multiple sclerosis. Nature 566, 543–547. 10.1038/s41586-019-0903-2.

99. Mathys, H., Peng, Z., Boix, C.A., Victor, M.B., Leary, N., Babu, S., Abdelhady, G., Jiang, X., Ng, A.P., Ghafari, K., et al. (2023). Single-cell atlas reveals correlates of high cognitive function, dementia, and resilience to Alzheimer’s disease pathology. Cell 186, 4365–4385.e27. 10.1016/j.cell.2023.08.039.

100. COvid-19 Multi-omics Blood ATlas (COMBAT) Consortium (2022). A blood atlas of COVID-19 defines hallmarks of disease severity and specificity. Cell 185, 916–938.e58. 10.1016/j.cell.2022.01.012.

101. Wolf, F.A., Angerer, P., and Theis, F.J. (2018). SCANPY: large-scale single-cell gene expression data analysis. Genome Biol. 19, 15. 10.1186/s13059-017-1382-0.

102. Murphy, A.E., and Skene, N.G. (2022). A balanced measure shows superior performance of pseudobulk methods in single-cell RNA-sequencing analysis. Nat. Commun. 13, 7851. 10.1038/s41467-022-35519-4.

103. Hoffman, G.E., Lee, D., Bendl, J., Prashant, N.M., Hong, A., Casey, C., Alvia, M., Shao, Z., Argyriou, S., Therrien, K., et al. (2024). Efficient differential expression analysis of large-scale single cell transcriptomics data using dreamlet. BioRxiv. 10.1101/2023.03.17.533005.

104. Hoffman, G.E., and Schadt, E.E. (2016). variancePartition: interpreting drivers of variation in complex gene expression studies. BMC Bioinformatics 17, 483. 10.1186/s12859-016-1323-z.

105. Gandal, M.J., Haney, J.R., Wamsley, B., Yap, C.X., Parhami, S., Emani, P.S., Chang, N., Chen, G.T., Hoftman, G.D., de Alba, D., et al. (2022). Broad transcriptomic dysregulation occurs across the cerebral cortex in ASD. Nature 611, 532–539. 10.1038/s41586-022-05377-7.

106. Zhang, B., and Horvath, S. (2005). A general framework for weighted gene co-expression network analysis. Stat. Appl. Genet. Mol. Biol. 4, Article17. 10.2202/1544-6115.1128.

107. Blauwendraat, C., Nalls, M.A., and Singleton, A.B. (2020). The genetic architecture of Parkinson’s disease. Lancet Neurol. 19, 170–178. 10.1016/S1474-4422(19)30287-X.

108. Kim, J.J., Vitale, D., Otani, D.V., Lian, M.M., Heilbron, K., 23andMe Research Team, Iwaki, H., Lake, J., Solsberg, C.W., Leonard, H., et al. (2024). Multi-ancestry genome-wide association meta-analysis of Parkinson’s disease. Nat. Genet. 56, 27–36. 10.1038/s41588-023-01584-8.

109. Leonard, H.L., and Global Parkinson’s Genetics Program (GP2) (2025). Novel parkinson’s disease genetic risk factors within and across european populations. medRxiv. 10.1101/2025.03.14.24319455.

110. Zhang, K., Zemke, N.R., Armand, E.J., and Ren, B. (2024). A fast, scalable and versatile tool for analysis of single-cell omics data. Nat. Methods 21, 217–227. 10.1038/s41592-023-02139-9.

111. Hao, Y., Stuart, T., Kowalski, M.H., Choudhary, S., Hoffman, P., Hartman, A., Srivastava, A., Molla, G., Madad, S., Fernandez-Granda, C., et al. (2024). Dictionary learning for integrative, multimodal and scalable single-cell analysis. Nat. Biotechnol. 42, 293–304. 10.1038/s41587-023-01767-y.

112. Hao, Y., Hao, S., Andersen-Nissen, E., Mauck, W.M., Zheng, S., Butler, A., Lee, M.J., Wilk, A.J., Darby, C., Zager, M., et al. (2021). Integrated analysis of multimodal single-cell data. Cell 184, 3573–3587. 10.1016/j.cell.2021.04.048.

113. Stuart, T., Butler, A., Hoffman, P., Hafemeister, C., Papalexi, E., Mauck, W.M., Hao, Y., Stoeckius, M., Smibert, P., and Satija, R. (2019). Comprehensive Integration of Single-Cell Data. Cell 177, 1888–1902.e21. 10.1016/j.cell.2019.05.031.

114. Butler, A., Hoffman, P., Smibert, P., Papalexi, E., and Satija, R. (2018). Integrating single-cell transcriptomic data across different conditions, technologies, and species. Nat. Biotechnol. 36, 411–420. 10.1038/nbt.4096.

115. Satija, R., Farrell, J.A., Gennert, D., Schier, A.F., and Regev, A. (2015). Spatial reconstruction of single-cell gene expression data. Nat. Biotechnol. 33, 495–502. 10.1038/nbt.3192.

116. Korsunsky, I., Millard, N., Fan, J., Slowikowski, K., Zhang, F., Wei, K., Baglaenko, Y., Brenner, M., Loh, P.-R., and Raychaudhuri, S. (2019). Fast, sensitive and accurate integration of single-cell data with Harmony. Nat. Methods 16, 1289–1296. 10.1038/s41592-019-0619-0.

117. Fu, R., Gillen, A.E., Sheridan, R.M., Tian, C., Daya, M., Hao, Y., Hesselberth, J.R., and Riemondy, K.A. (2020). clustifyr: an R package for automated single-cell RNA sequencing cluster classification. F1000Res. 9, 223. 10.12688/f1000research.22969.2.

118. Yu, G. (2024). Thirteen years of clusterProfiler. Innovation (Camb) 5, 100722. 10.1016/j.xinn.2024.100722.

119. Xu, S., Hu, E., Cai, Y., Xie, Z., Luo, X., Zhan, L., Tang, W., Wang, Q., Liu, B., Wang, R., et al. (2024). Using clusterProfiler to characterize multiomics data. Nat. Protoc. 19, 3292–3320. 10.1038/s41596-024-01020-z.

120. Wu, T., Hu, E., Xu, S., Chen, M., Guo, P., Dai, Z., Feng, T., Zhou, L., Tang, W., Zhan, L., et al. (2021). clusterProfiler 4.0: A universal enrichment tool for interpreting omics data. Innovation (Camb) 2, 100141. 10.1016/j.xinn.2021.100141.

121. Yu, G., Wang, L.-G., Han, Y., and He, Q.-Y. (2012). clusterProfiler: an R package for comparing biological themes among gene clusters. OMICS 16, 284–287. 10.1089/omi.2011.0118.

122. Korotkevich, G., Sukhov, V., Budin, N., Shpak, B., Artyomov, M.N., and Sergushichev, A. (2016). Fast gene set enrichment analysis. BioRxiv. 10.1101/060012.

123. Laddach, A., and Shapiro, M. (2024). CatsCradle (Bioconductor).

124. Huang, F., Hartwich, T.M.P., Rivera-Molina, F.E., Lin, Y., Duim, W.C., Long, J.J., Uchil, P.D., Myers, J.R., Baird, M.A., Mothes, W., et al. (2013). Video-rate nanoscopy using sCMOS camera-specific single-molecule localization algorithms. Nat. Methods 10, 653–658. 10.1038/nmeth.2488.

125. Fu, B., Brock, E.E., Andrews, R., Breiter, J.C., Tian, R., Toomey, C.E., Lachica, J., Lashley, T., Ryten, M., Wood, N.W., et al. (2024). RASP: optimal single puncta detection in complex cellular backgrounds. J. Phys. Chem. B 128, 3585–3597. 10.1021/acs.jpcb.4c00174.

126. Yen, J.C., Chang, F.J., and Chang, S. (1995). A new criterion for automatic multilevel thresholding. IEEE Trans. Image Process. 4, 370–378. 10.1109/83.366472.

127. van der Walt, S., Schönberger, J.L., Nunez-Iglesias, J., Boulogne, F., Warner, J.D., Yager, N., Gouillart, E., Yu, T., and scikit-image contributors (2014). scikit-image: image processing in Python. PeerJ 2, e453. 10.7717/peerj.453.

128. Hoyer, W., Antony, T., Cherny, D., Heim, G., Jovin, T.M., and Subramaniam, V. (2002). Dependence of alpha-synuclein aggregate morphology on solution conditions. J. Mol. Biol. 322, 383–393. 10.1016/s0022-2836(02)00775-1.

129. Thirunavukkuarasu, S., Jares-Erijman, E.A., and Jovin, T.M. (2008). Multiparametric fluorescence detection of early stages in the amyloid protein aggregation of pyrene-labeled alpha-synuclein. J. Mol. Biol. 378, 1064–1073. 10.1016/j.jmb.2008.03.034.

130. Bąk, K.M., Edwards, D.C., George, D., Singh, B., Ferguson, R., Zhao, T., Piché, K., Louwrier, A., Cockroft, S.L., and Horrocks, M.H. (2025). A Single-Molecule Liposome Assay for Membrane Permeabilization. Angew. Chem. Int. Ed 64, e202503678. 10.1002/anie.202503678.

131. Brown, J.W.P., Bauer, A., Polinkovsky, M.E., Bhumkar, A., Hunter, D.J.B., Gaus, K., Sierecki, E., and Gambin, Y. (2019). Single-molecule detection on a portable 3D-printed microscope. Nat. Commun. 10, 5662. 10.1038/s41467-019-13617-0.

132. Germain, P.-L., Lun, A., Garcia Meixide, C., Macnair, W., and Robinson, M.D. (2021). Doublet identification in single-cell sequencing data using scDblFinder. F1000Res. 10, 979. 10.12688/f1000research.73600.2.

133. Love, M.I., Huber, W., and Anders, S. (2014). Moderated estimation of fold change and dispersion for RNA-seq data with DESeq2. Genome Biol. 15, 550. 10.1186/s13059-014-0550-8.

