## Supplemental Document for "Microglial activation and alpha-synuclein oligomers drive the early inflammatory phase of Parkinson’s disease"

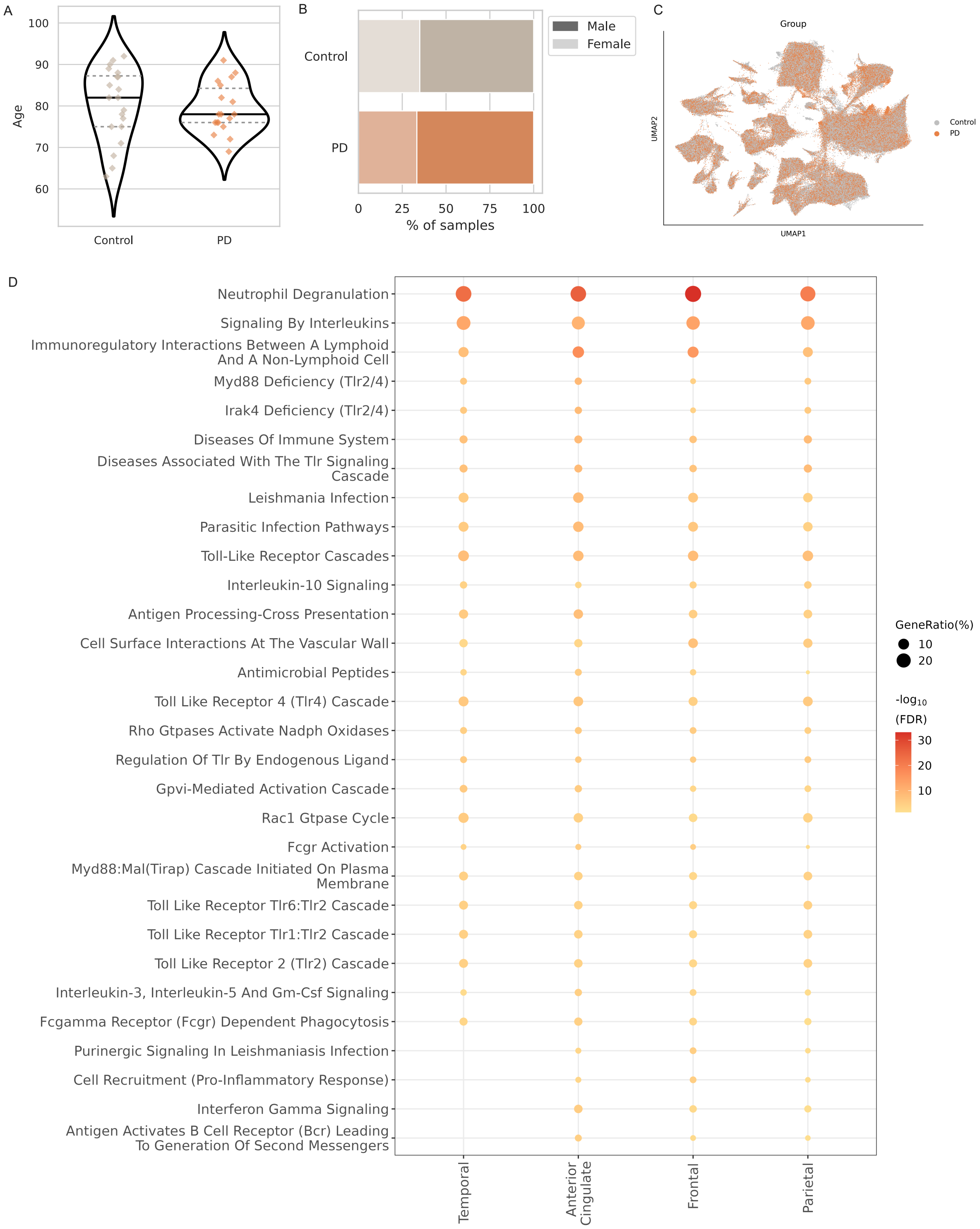


**Supplementary Figure 1. Extended data related to main Figure 2.**
(A) Age at time of death (years) for the control and PD cohorts

(B) Sex distribution for the control and PD cohorts

(C) Distribution of control and PD nuclei in snRNAseq data following batch correction

(D) REACTOME pathway analysis of significantly upregulated genes in the four cortical brain areas analysed by bulk RNA-seq. The top 30 pathways are shown and arranged by descending FDR-adjusted p values. The colour of each dot denotes the -log10(FDR-adjusted p value) and the size of dot indicates the percent of genes from a pathway that were significantly upregulated in each brain area.

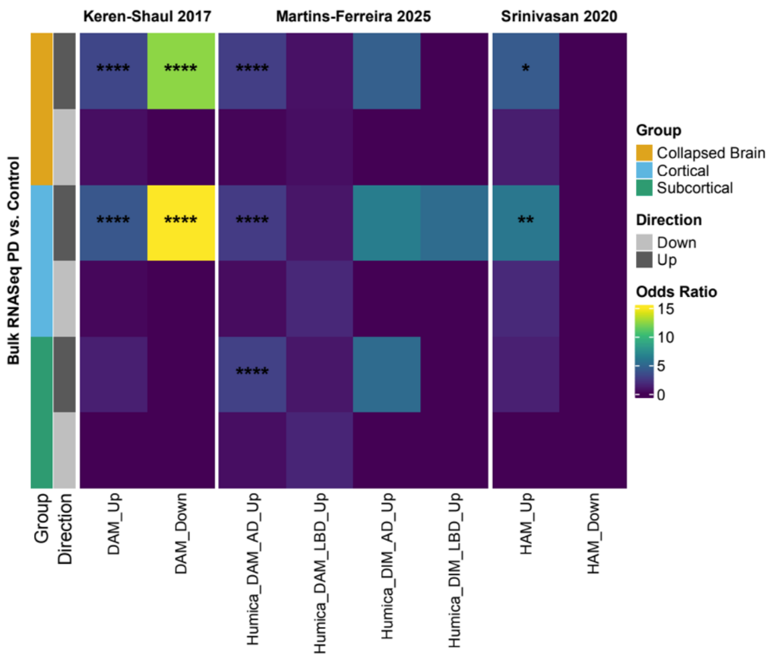


**Supplementary Figure 2. Overlap of Braak stage 3–4 PD bulk RNA-seq with AD and disease-associated microglial signatures.**

Heatmap showing the overlap of significant up- and downregulated differentially expressed genes derived from cortical, subcortical and combined (collapsed) regions with published disease associated microglial signatures. Collapsing brain regions: upregulated genes overlapped with the DAM signature (up: FDR-corrected $p = 3.96 \times 10^{-7}$, OR 3.15; down: FDR-corrected $p = 3.96 \times 10^{-7}$, OR 12.50); HuMiCA DAM AD signature (up: FDR-corrected $p = 9.50 \times 10^{-19}$, OR 2.73); and HAM signature (up: FDR-corrected $p = 1.51 \times 10^{-2}$, OR 4.24). These patterns persisted when confined to cortical regions (FDR-corrected $p = 2.30 \times 10^{-8}$, OR 4.02, FDR-corrected $p = 3.96 \times 10^{-7}$, OR 15.05, FDR-corrected $p = 7.87 \times 10^{-13}$ OR 2.61 and $p = 4.42 \times 10^{-3}$ OR 5.99, respectively). Subcortical regions only significantly overlapped with HuMiCA DAM signatures (up: FDR-corrected $p = 1.12 \times 10^{-9}$, OR 2.87). DAM = Disease Associated Microglia, DIM = Disease-inflammatory macrophages, HAM = Human Alzheimer's Microglia, OR = odds ratio.

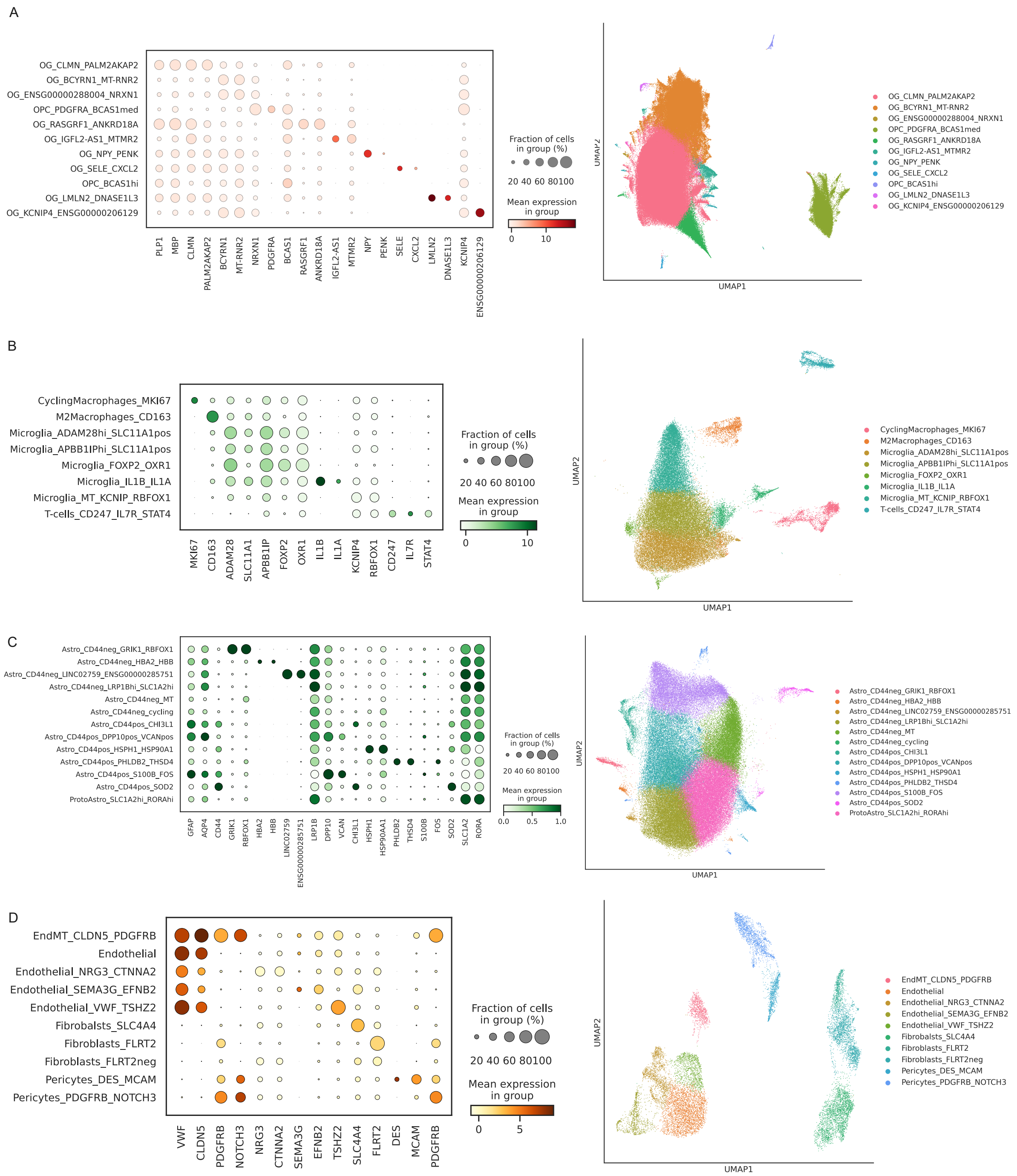


**Supplementary Figure 3. Cell state annotation of snRNAseq data for oligodendrocytes, immune cells, astrocytes, and the endomural compartment.**

(A) Cell state annotation for oligodendrocytes. UMAP shows nuclei following preprocessing, dimensionality reduction and clustering. Dotplots show the fraction of cells in that cluster which express the gene (dot diameter) and mean expression of that gene (colour intensity). Presented genes include in-house annotation gene panel and data-driven markers.

(B) Cell state annotation for immune cells (CD74+). UMAP shows nuclei following preprocessing, dimensionality reduction and clustering. Dotplots show the fraction of cells in that cluster which express the gene (dot diameter) and mean expression of that gene (colour intensity). Presented genes include in-house annotation gene panel and data-driven markers.

(C) Cell state annotation for astrocytes. UMAP shows nuclei following preprocessing, dimensionality reduction and clustering. Dotplots show the fraction of cells in that cluster which express the gene (dot diameter) and mean expression of that gene (colour intensity). Presented genes include in-house annotation gene panel and data-driven markers.

(D) Cell state annotation for endomural compartment. UMAP shows nuclei following preprocessing, dimensionality reduction and clustering. Dotplots show the fraction of cells in that cluster which express the gene (dot diameter) and mean expression of that gene (colour intensity). Presented genes include in-house annotation gene panel and data-driven markers.

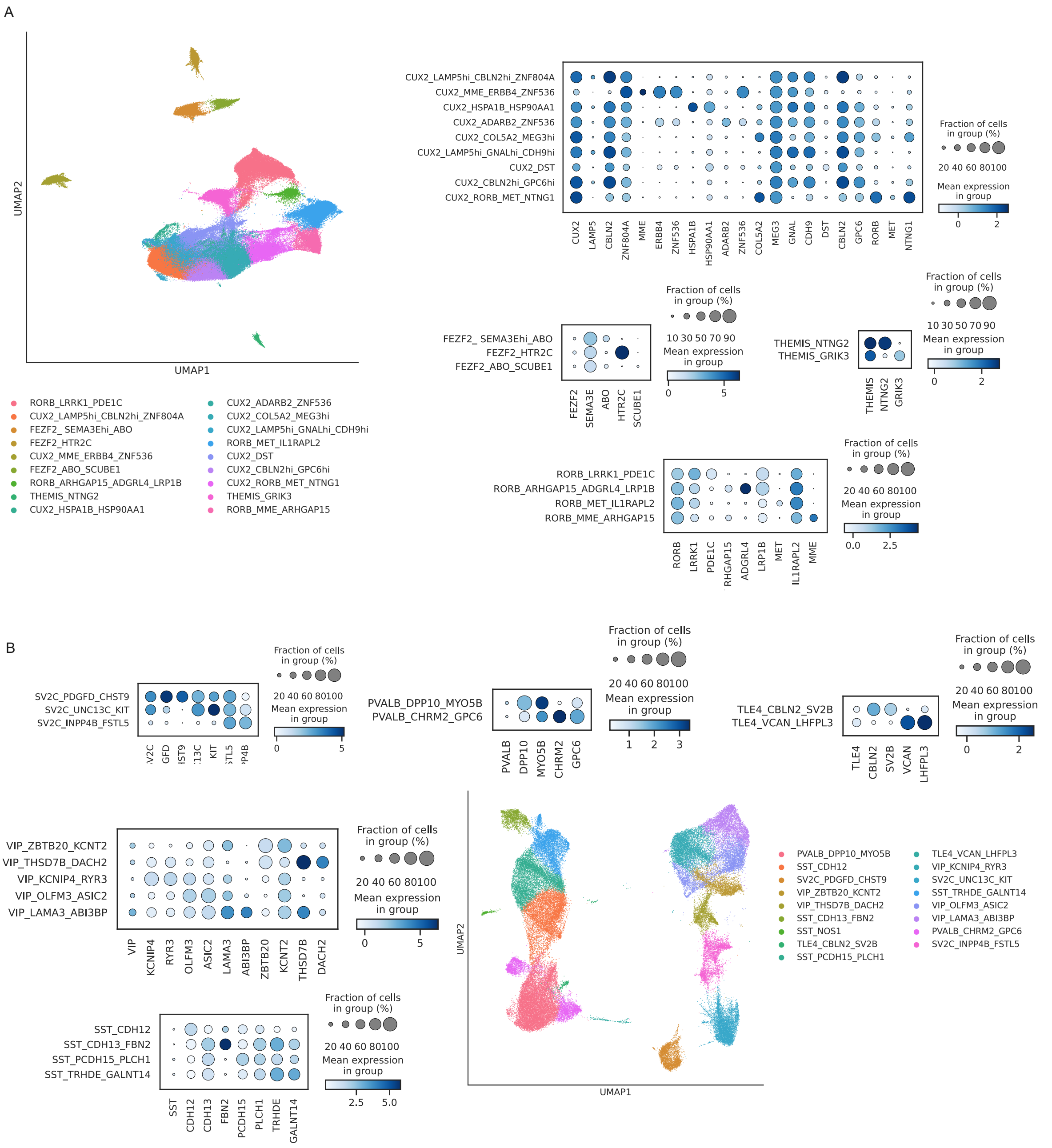


**Supplementary Figure 4. Cell state annotation of snRNAseq data for neurons.**
(A) Cell state annotation for excitatory neurons. UMAP shows nuclei following preprocessing, dimensionality reduction and clustering. Dotplots show the fraction of cells in that cluster which express the gene (dot diameter) and mean expression of that gene (colour intensity). Excitatory neurons have been partitioned into CUX2+, RORB+, FEZF2+ and THEMIS+ cell types. Presented genes include in-house annotation gene panel and data-driven markers.

(B) Cell state annotation for inhibitory neurons. UMAP shows nuclei following preprocessing, dimensionality reduction and clustering. Dotplots show the fraction of cells in that cluster which express the gene (dot diameter) and mean expression of that gene (colour intensity). Inhibitory neurons have been partitioned into SST+, VIP+, TLE4+, PVALB+ and SV2C+ cell types. Presented genes include in-house annotation gene panel and data-driven markers.

A

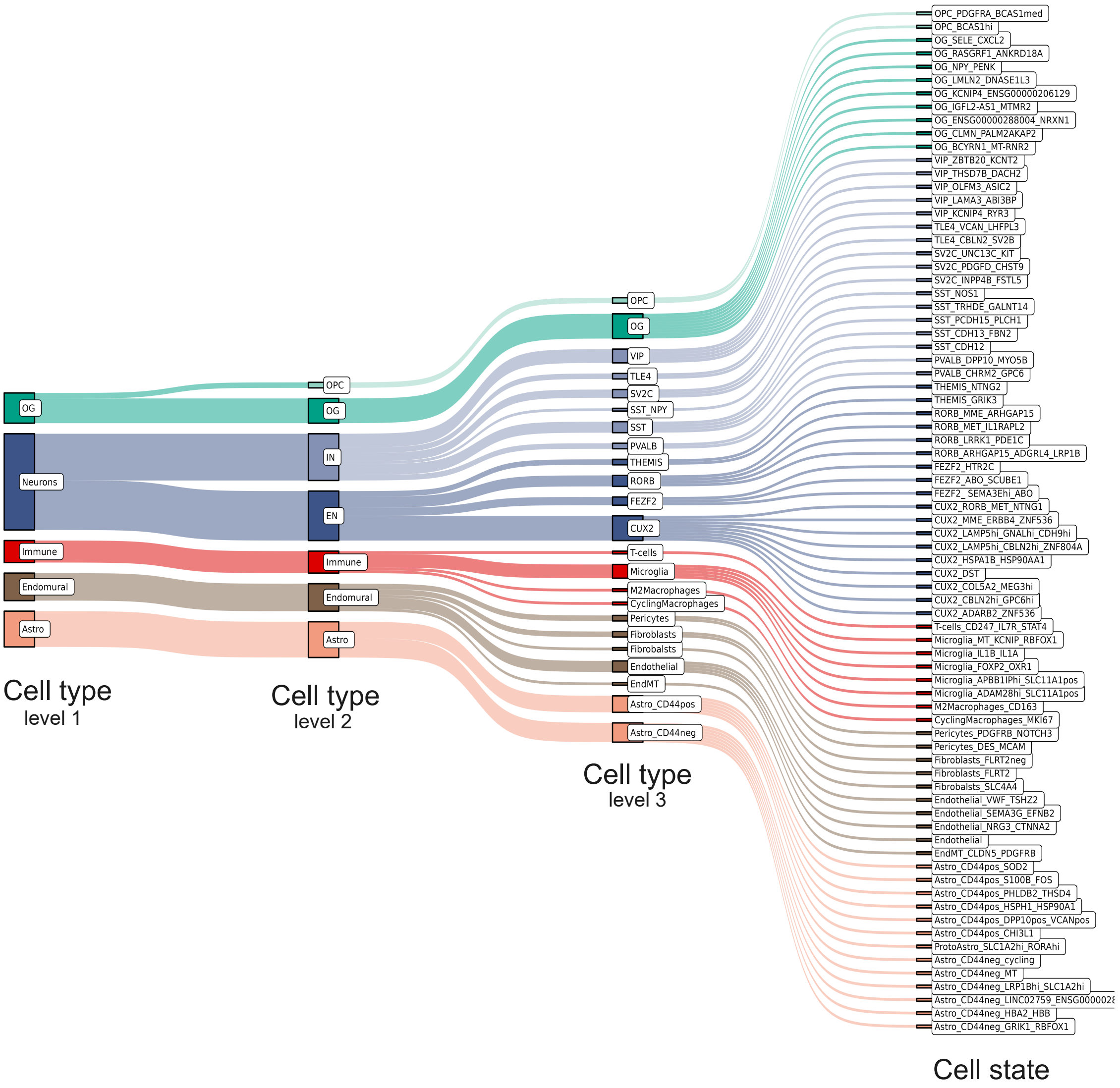


**Supplementary Figure 5. Hierarchical structure of cell type and cell state annotation.**
(A) Schematic representation of the hierarchy used for annotating cell types and their corresponding cell states in the snRNAseq dataset. Levels 1, 2 and 3 have annotation based on gene expression markers, whilst level 4 annotation is based on data-driven gene expression.

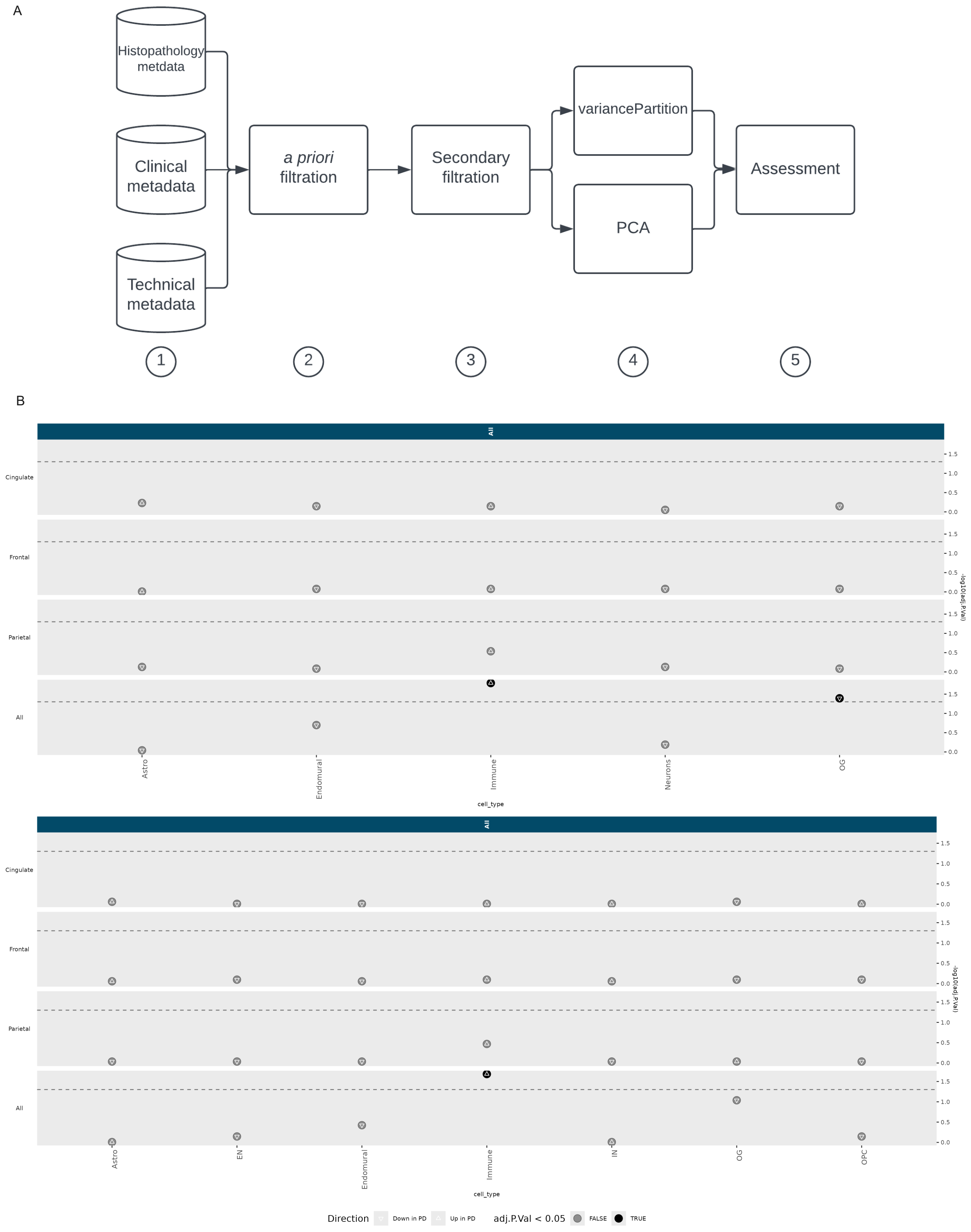


**Supplementary Figure 6. Covariate correction pipeline and cell type proportion differences across annotation levels.**

(A) Covariate correction pipeline used for all downstream analysis.

(B) Results of cell type proportion analysis in annotation level 1 and 2. Dotted line represents significance threshold, where clusters with significant cell type proportion differences are indicated in black. This analysis was performed for each brain region in snRNAseq analysis (frontal, parietal and cingulate cortex), as well as pooling all brain regions together. Immune cells and oligodendrocyte have significant differences in cell type proportion when pooling all brain regions at annotation level 1. The difference in immune cell proportion remains significant at annotation level 2.

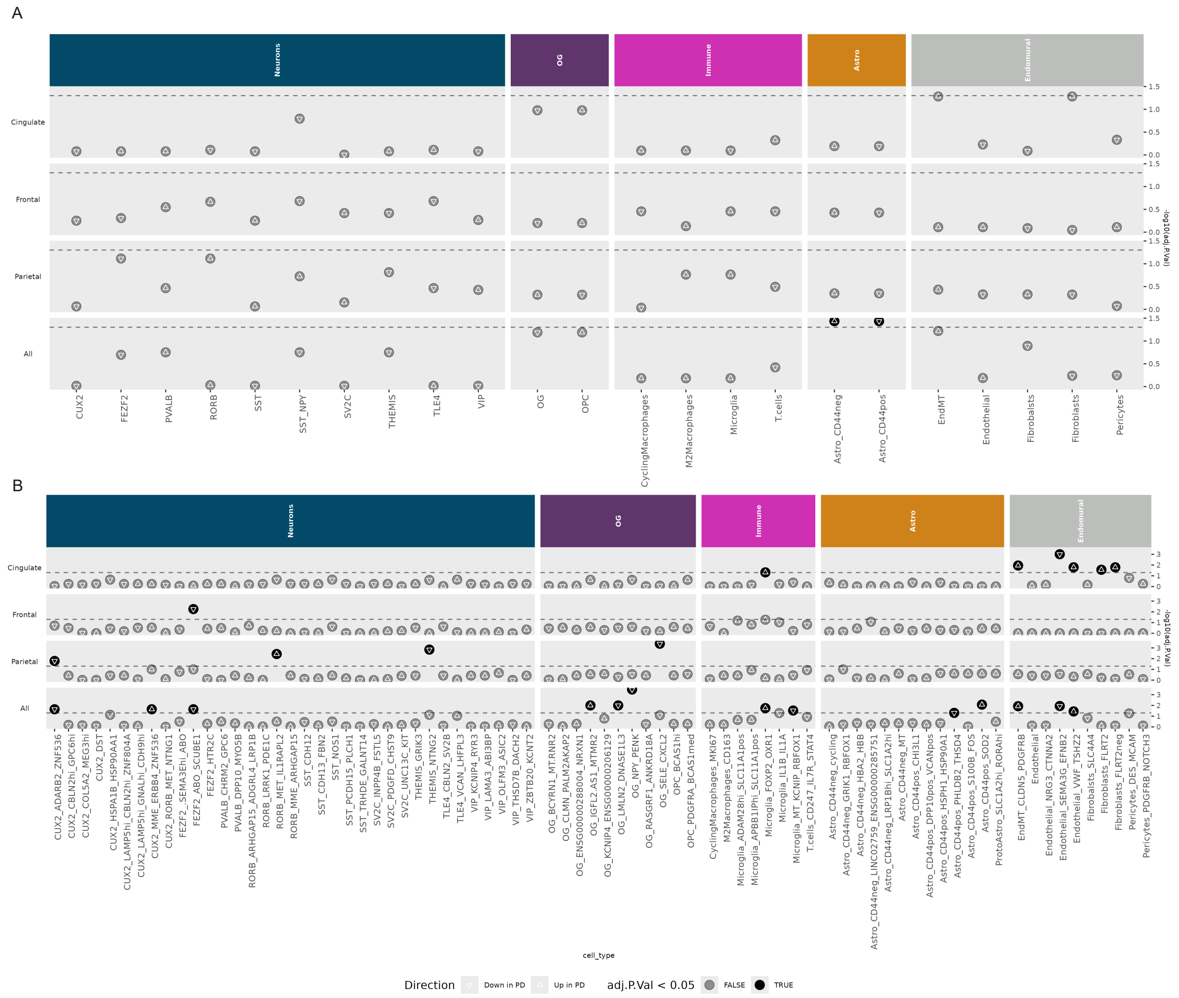

**Supplementary Figure 7. Cell type and cell state proportion differences across annotation levels 3 and 4.**

(A) Cell type proportion analysis in annotation level 3. Dotted line represents significance threshold, where clusters with significant cell type proportion differences are indicated in black. This analysis was performed for each brain region in snRNAseq analysis (frontal, parietal and cingulate cortex), as well as pooling all brain regions together. Both astrocyte populations have a significant difference when pooling all brain regions.

(B) Cell type proportion analysis in annotation level 4. Dotted line represents significance threshold, where clusters with significant cell type proportion differences are indicated in black. This analysis was performed for each brain region in snRNAseq analysis (frontal, parietal and cingulate cortex), as well as pooling all brain regions together. In the cingulate cortex microglia and the endomural compartment have cell type proportion differences. In the frontal cortex FEZF2 inhibitory neuron also has significant differences. In the parietal cortex excitatory neurons and oligodendrocytes have significant differences. In the analysis pooling all brain regions a total of 13 cell states have proportion differences which are present across all cell type compartments.

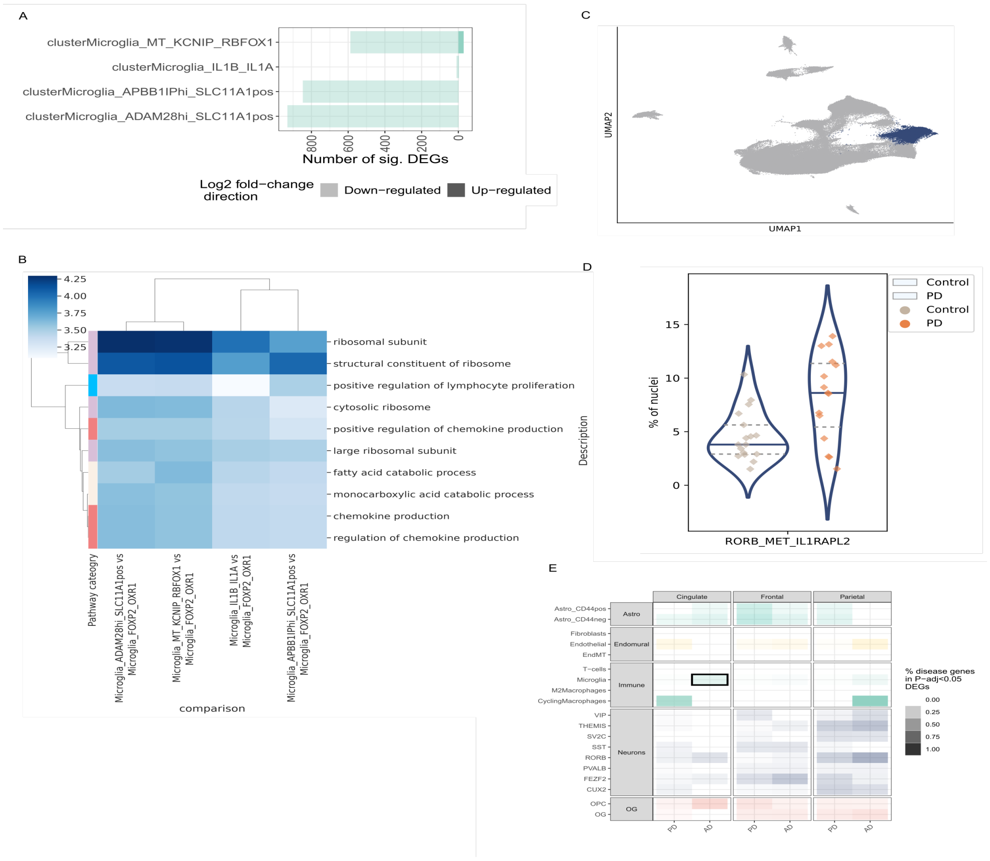


**Supplementary Figure 8. Further characterisation of cell states of interest.**

(A) Results of differential gene expression analysis among microglial clusters, specifically when comparing FOXP2_OXR1 microglia to all other microglia clusters.

(B) Top 10 pathways resulting from gene set enrichment analysis of FOXP2_OXR1 microglia DEGs when compared to each of the other microglia clusters. In order to identify pathways uniquely characteristic of the FOXP2_OXR1 cluster, we present here pathways which were significantly different in FOXP2_OXR1 microglia and every other microglial cluster. The colour bar indicates the pathway category: lavender – ribosomal genes; sky blue – lymphocyte-associated pathways; pink – chemokine production; off-white – metabolic.

(C) UMAP highlighting the RORB_MET_IL1RAPL2 cluster among excitatory neurons.

(D) Cell type proportion differences of the RORB_MET_IL1RAPL2 in the parietal cortex (FDR-adjusted p value = $3.70 \times 10^{-3}$), contrasting proportions in control and PD.

(E) Enrichment of causal disease gene among cell-specific differentially expressed genes. This was assessed using PD and AD causal genes. Highlighted with the black box are the cell types per region which have significant enrichment of disease-associated genes (FDR-corrected p = $4.13 \times 10^{-2}$). Differentially expressed genes in the microglia of the cingulate cortex are enriched for genes associated with AD.

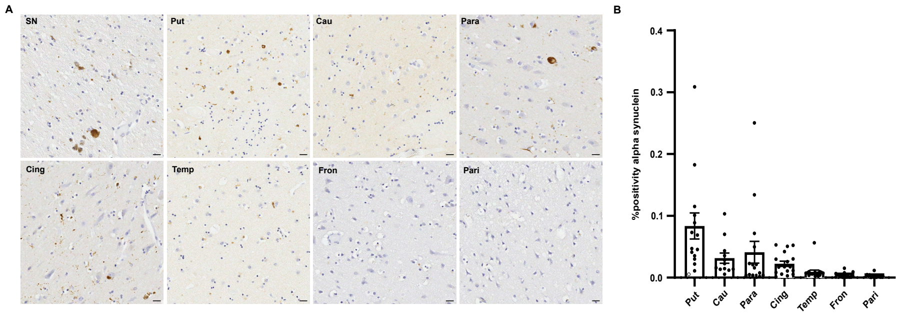


### Supplementary Figure 9. LB pathology quantified across regions in our Braak stage 3–4 PD cohort

(A) Exemplar images across all brain regions DAB-stained for αSyn, highlighting differing pathology dependent on brain region.

(B) Quantification of the αSyn positivity from DAB-staining normalised to area.

### ASA-PD (Advanced Sensing of Aggregates—Parkinson's Disease)

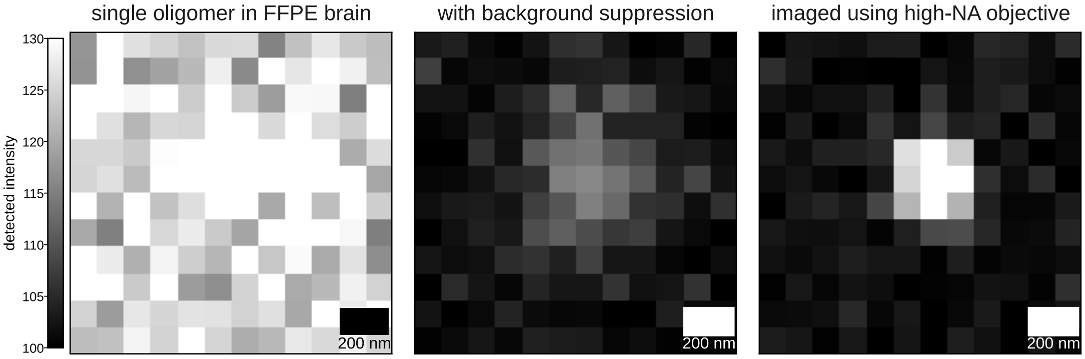


**Supplementary Figure 10. Schematic of the ASA-PD concept (see Andrews et. al.).**
A single oligomer in the FFPE brain, imaged with a low-NA objective, is difficult to see due to high background and the low collection efficiency. Background suppression improves the visibility of the punctum, and increasing the NA improves the visibility yet further due to the higher collection efficiency.

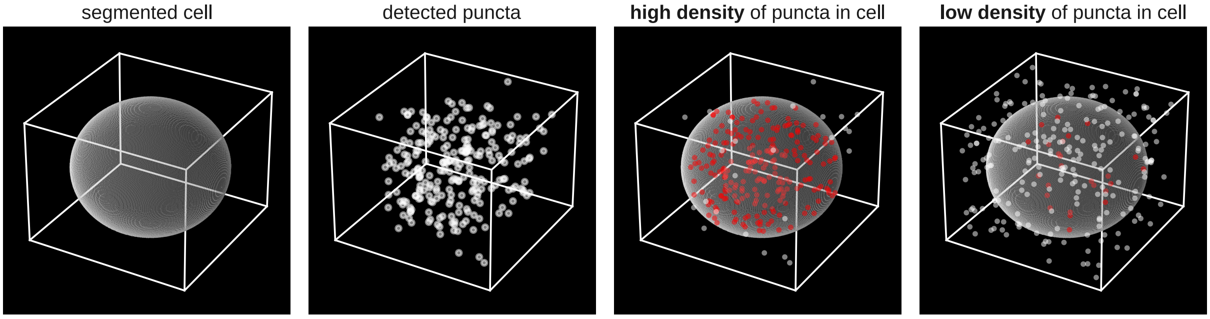


**Supplementary Figure 11. Schematic of the colocalization analysis for ASA-PD.**
From left to right, we show a “cartoon” segmented cell, overlaid with detected puncta. We then show examples where would detect a high density of puncta inside the cell volume, and a low density of puncta inside the cell volume.

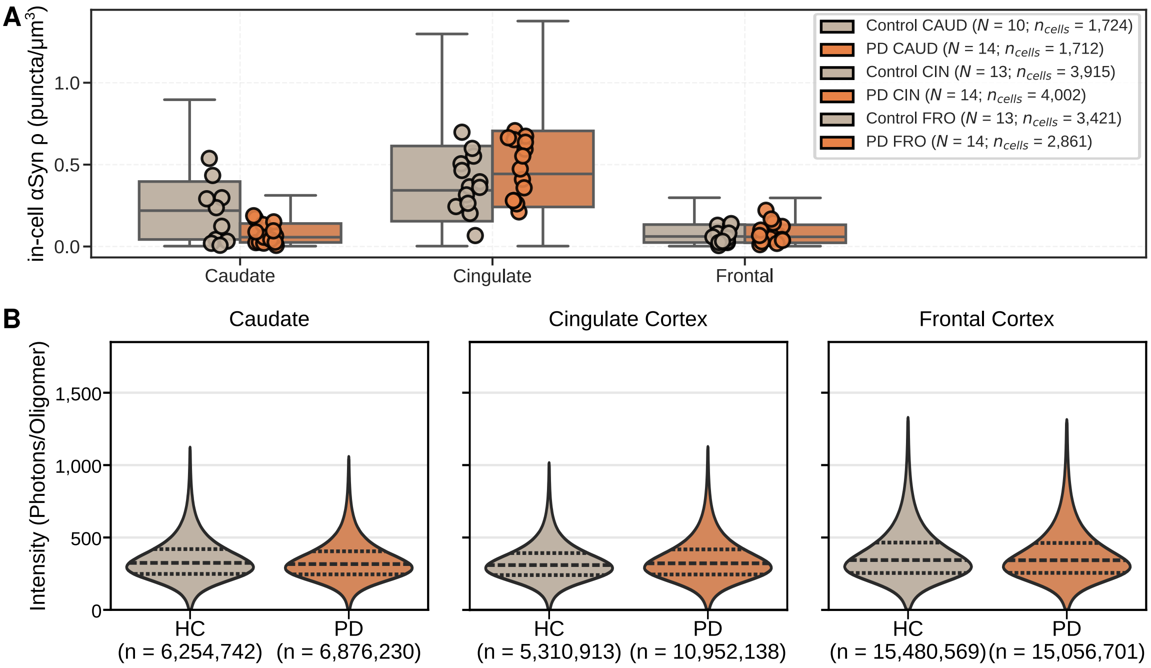


**Supplementary Figure 12. Extended data related to main Figure 4.**

(A) Puncta density in neurons across brain regions. Unlike in microglia, we do not see a noticeable difference across regions.

(B) Puncta intensity across brain regions, highlighting that there is no quantitative difference in the puncta between HC and PC cases.

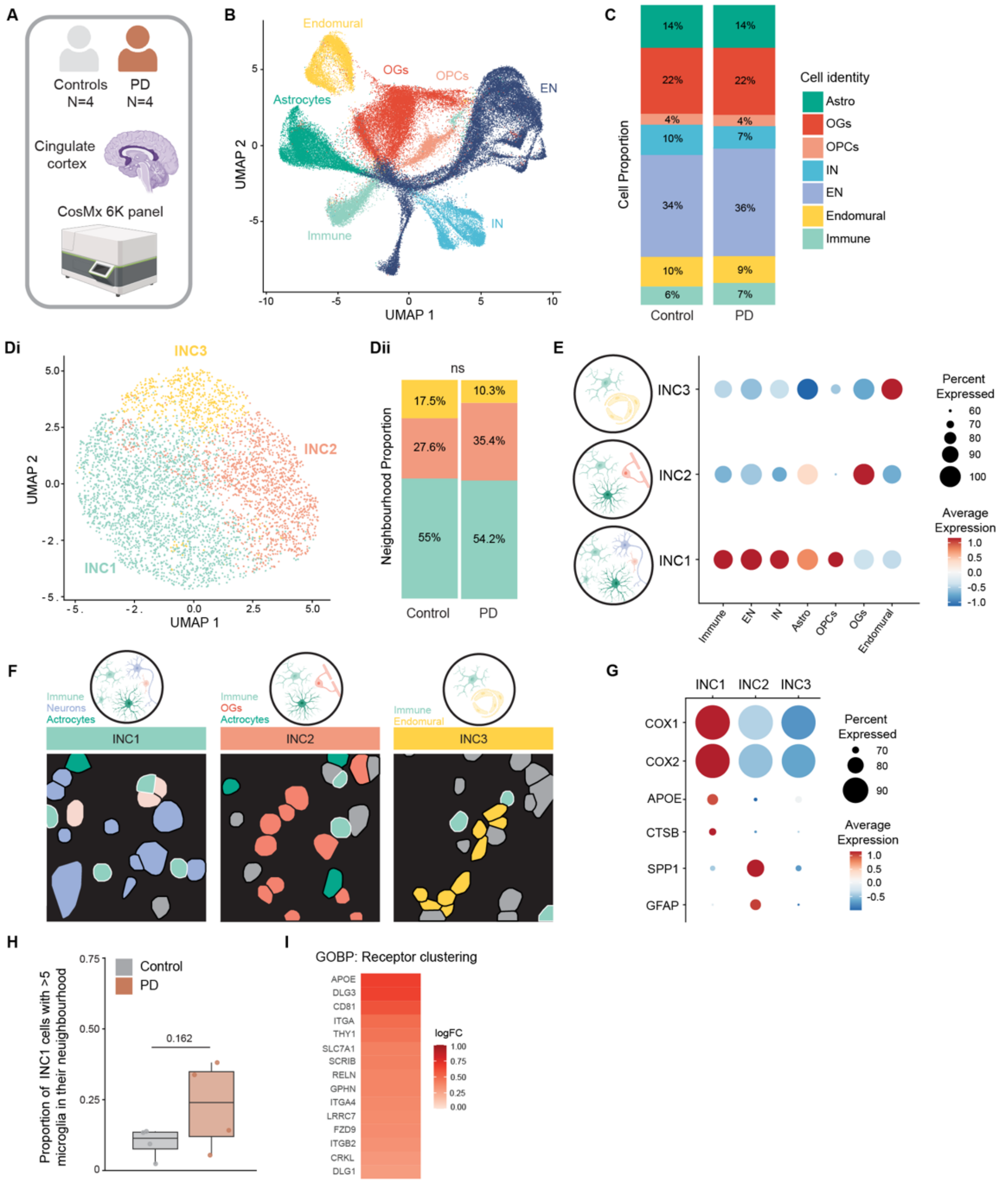

**Supplementary Figure 13. Spatial transcriptomics analysis of anterior cingulate cortex in Braak 3–4 PD.**

(A) Schematic of the CosMx pipeline.

(B) UMAP showing different CosMx cell types annotated with snRNAseq from the same individuals.

(C) Cell proportion of the 7 major cell types in control and PD.

(Di) UMAP showing the three different CosMx immune neighbourhood clusters (INCs).

(Dii) Proportion of the three immune neighbourhoods in control and PD.

(E) Dotplot presenting INCs and neighbouring cell types. The size of the dot encodes the percentage of cells within a class, while the colour encodes the number of neighbours (average expression level) across all cells within a class.

(F) CosMx segmentation representative images of INCs. Cells in light blue with a white egde represent the immune cells belonging to either INC1, INC2 or INC3. As presented in cartoon above summarising main neighbours found within the cluster, segmented cells in violet represent neurons, dark green astrocytes, light pink OPCs, red OGs and yellow endomural cells. Other cells not found as key neighbours in the cluster as shaded in grey for visual purposes.

(G) Dotplot representing cell markers for INC1, INC2 and INC3 determined using Seurat's FindAllMarkers function (p-value adj. <= 0.05, present in >70% cells). The size of the dot encodes the percentage of cells within a class, while the colour encodes the average gene expression level across all cells within a class.

(H) Analysis of INC1 immune-immune interaction, suggesting a trend in PD microglia being more commonly present in a neighbourhood of other microglia. We separated the immune-immune interactions into high (n = 410, immune cell neighbouring >5 other immune cells) and low (n = 2037, immune cell neighbouring <5 other immune cells) (Unpaired t-test, p value = 0.162).

(I) INC1 GSEA analysis of ranked log2FC values identified GO:BP receptor clustering pathway as significantly upregulated (FDR-corrected p value = 0.049) in PD. Genes found in the pathway and their associated logFC are presented.

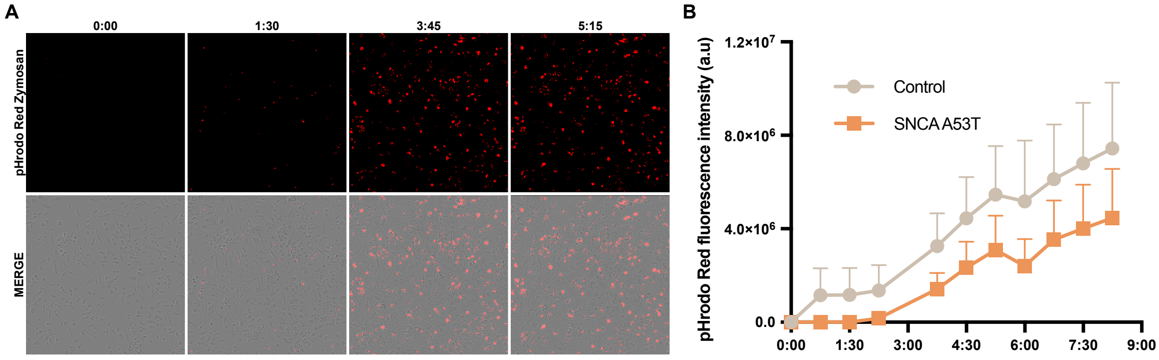


**Supplementary Figure 14. Phagocytosis time course of iMGL from control and *SNCA* A53T donors.**

(A) Representative images of iMGL incubated with pHrodo™ Zymosan particles at 0:00, 1:30, 3:45, and 5:15 hours, showing progressive particle uptake.

(B) Quantification of phagocytosis over a 9 hour time course in control (n = 4 donor lines) and *SNCA* A53T (n = 3 donor lines) iMGL. Data represent mean ± SEM.

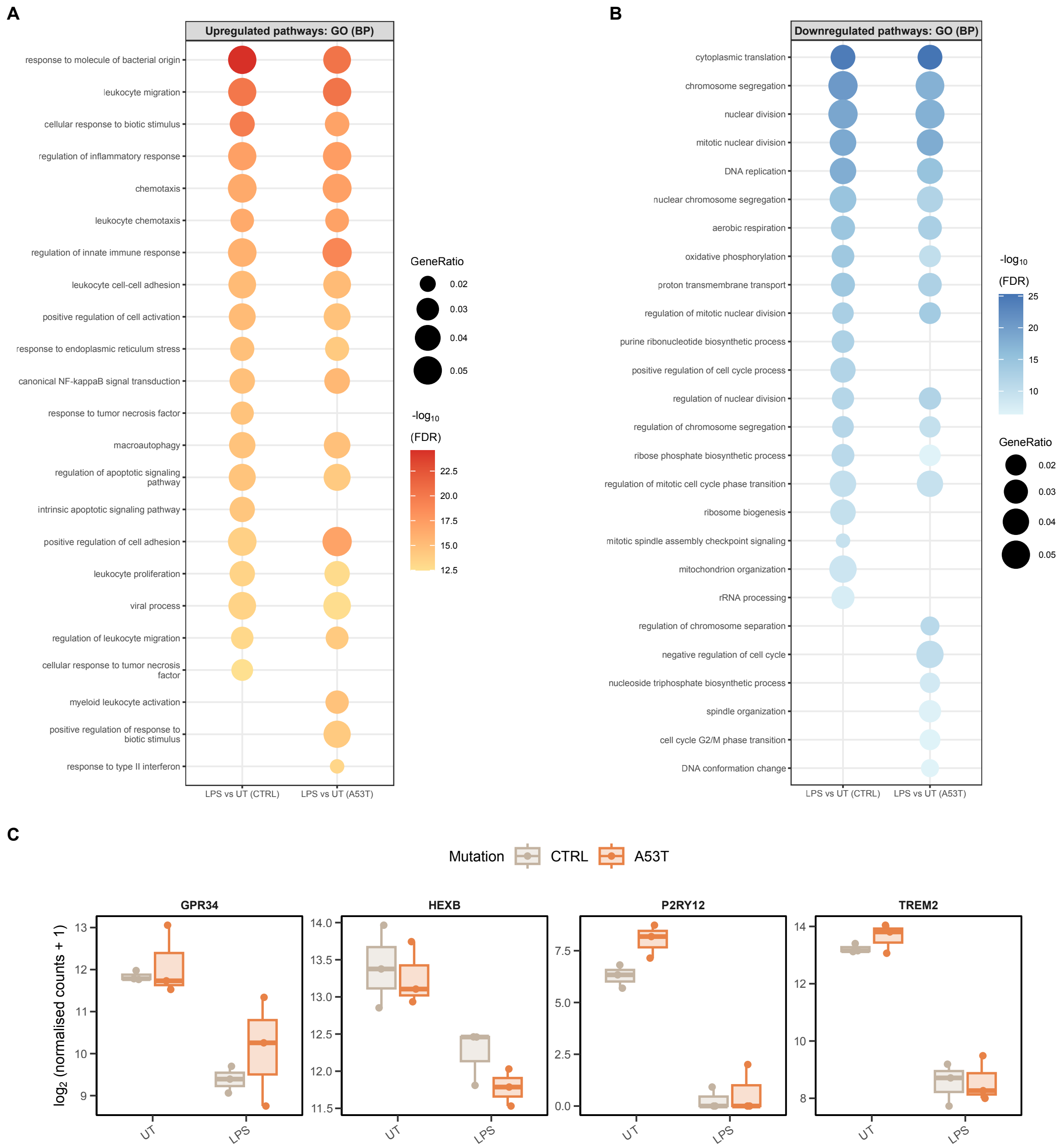


**Supplementary Figure 15. Bulk RNA-seq analysis of control and *SNCA* A53T iMGL with and without LPS stimulation.**

(A) Gene Ontology (GO) Biological Process (BP) pathways significantly upregulated in LPS vs UT for control and *SNCA* A53T genotypes. The top 20 terms are shown, ranked by FDR-corrected p value.

(B) Gene Ontology (GO) Biological Process (BP) pathways significantly downregulated in LPS vs UT for control and *SNCA* A53T genotypes. The top 20 terms are shown, ranked by FDR-corrected p value.

(C) Expression of microglial genes *GPR34*, *HEXB*, *P2RY12*, *TREM2* ($log_2$(normalised counts + 1) across genotype and treatment.

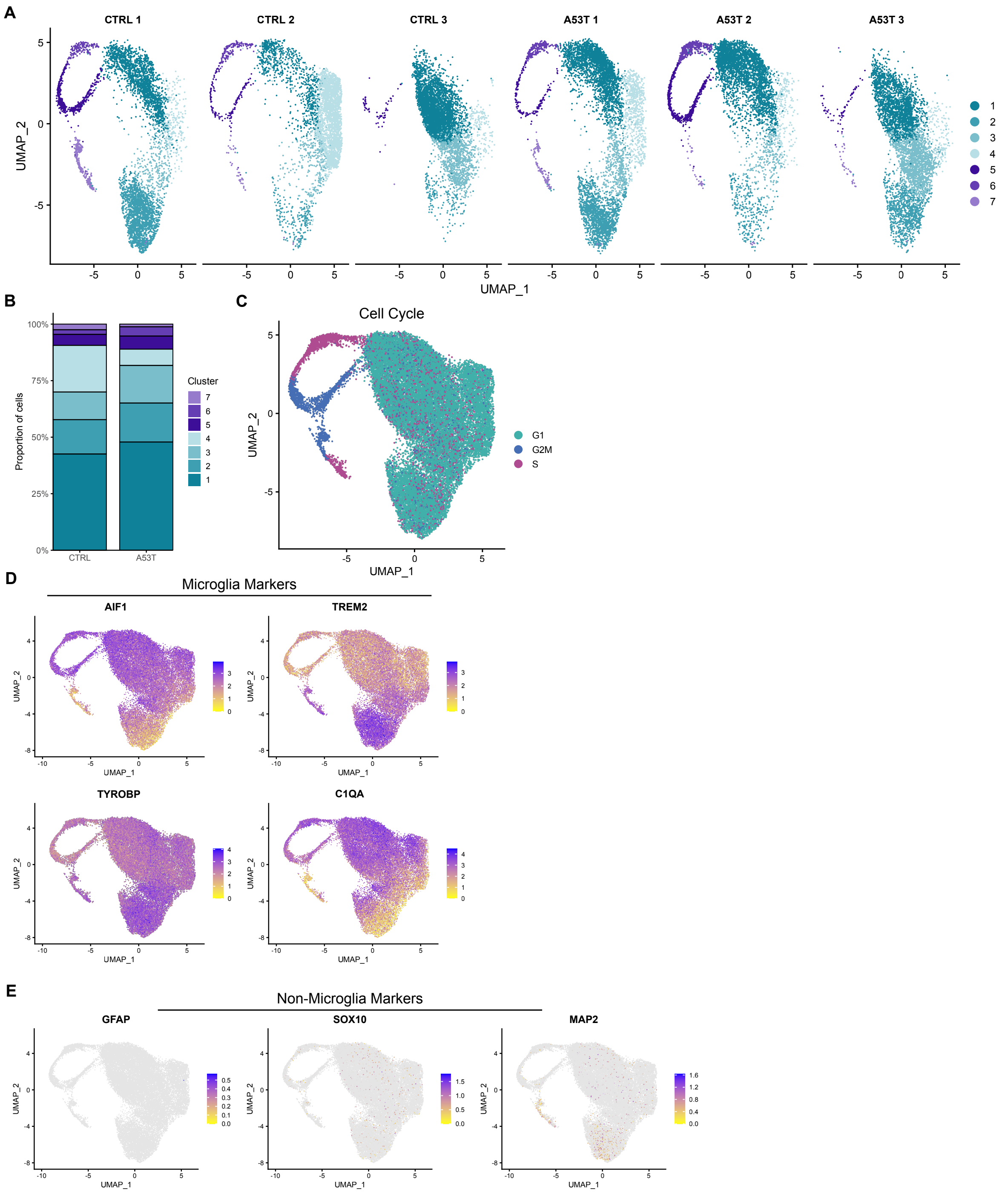


**Supplementary Figure 16. Extended single-cell RNA-seq analysis of iMGL derived from control and *SNCA* A53T iPSC lines.**

(A) UMAP plots of the integrated single-cell dataset shown separately for each donor.

(B) Proportion of each identified cluster in control and *SNCA* A53T lines.

(C) UMAP coloured by cell cycle phase (G1, G2M, S) based on cell cycle scoring.

(D) UMAP feature plots showing expression of canonical microglial markers *AIF1*, *TREM2*, *TYROBP*, and *C1QA*.

(E) UMAP feature plots showing low expression of non-microglial lineage markers. *GFAP* for astrocytes, *SOX10* for oligodendrocytes, and *MAP2* for neurons.

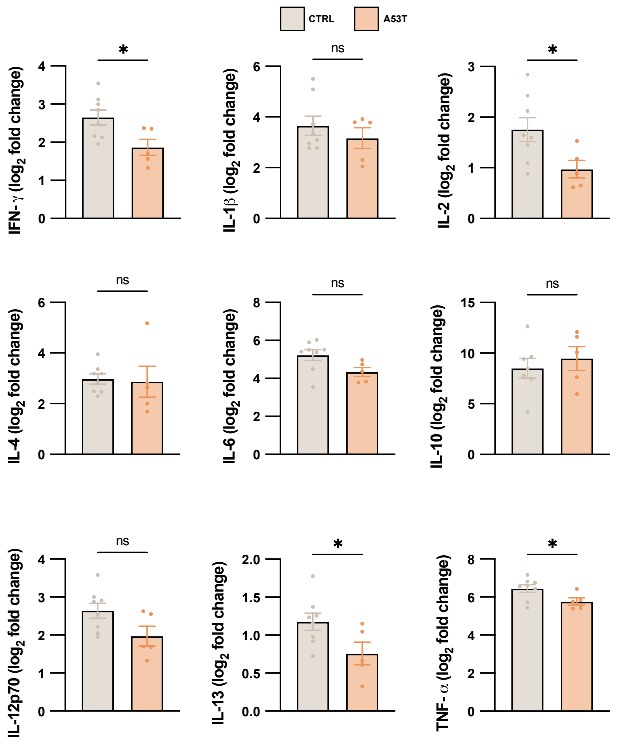


**Supplementary Figure 17. Cytokine responses of control and A53T iMGL to LPS stimulation.** iMGL from control (n = 8 iPSC lines) or A53T (n = 5 iPSC lines) donors were treated with LPS (10 ng/ml) and cytokine secretion was measured by ELISA from conditioned media 24 hours after exposure. Data are shown as fold change relative to baseline (untreated) for each genotype (data represent mean ± SEM; unpaired t-tests; $^{*}p < 0.05$).

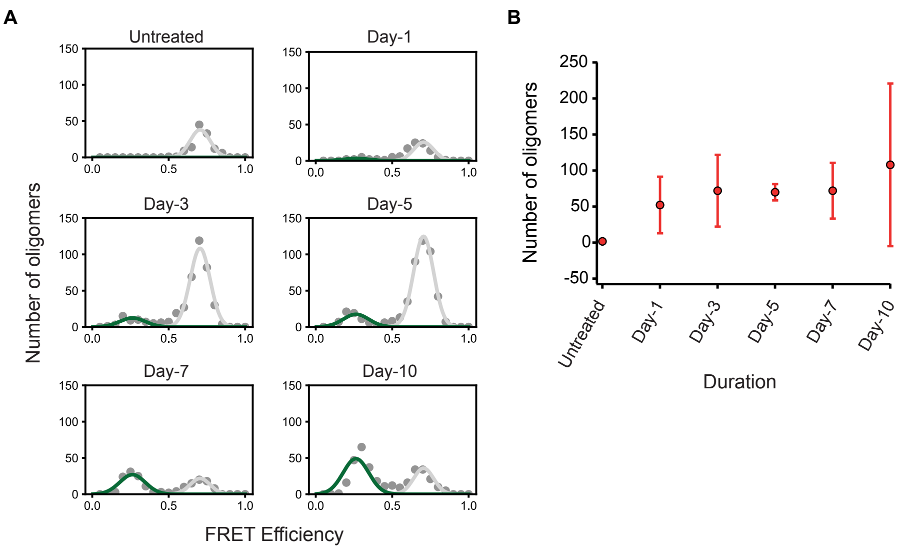


**Supplementary Figure 18. Single-molecule detection of oligomers in cell lysate**

(A) Example FRET histograms of oligomers present in cell lysate from cells either untreated or 1-, 3-, 5-, 7-, and 10-days post-treatment. Histograms were globally fit to two Gaussian distributions, centred at E = 0.26 and E = 0.71. The low-FRET peak (green) corresponds to oligomers, whereas the high-FRET peak (light grey) is from long Stokes shift autofluorescence.

(B) Number of oligomers determined by integrating the low-FRET peak (points show mean +/- S.D., n = 3 iPSC lines).

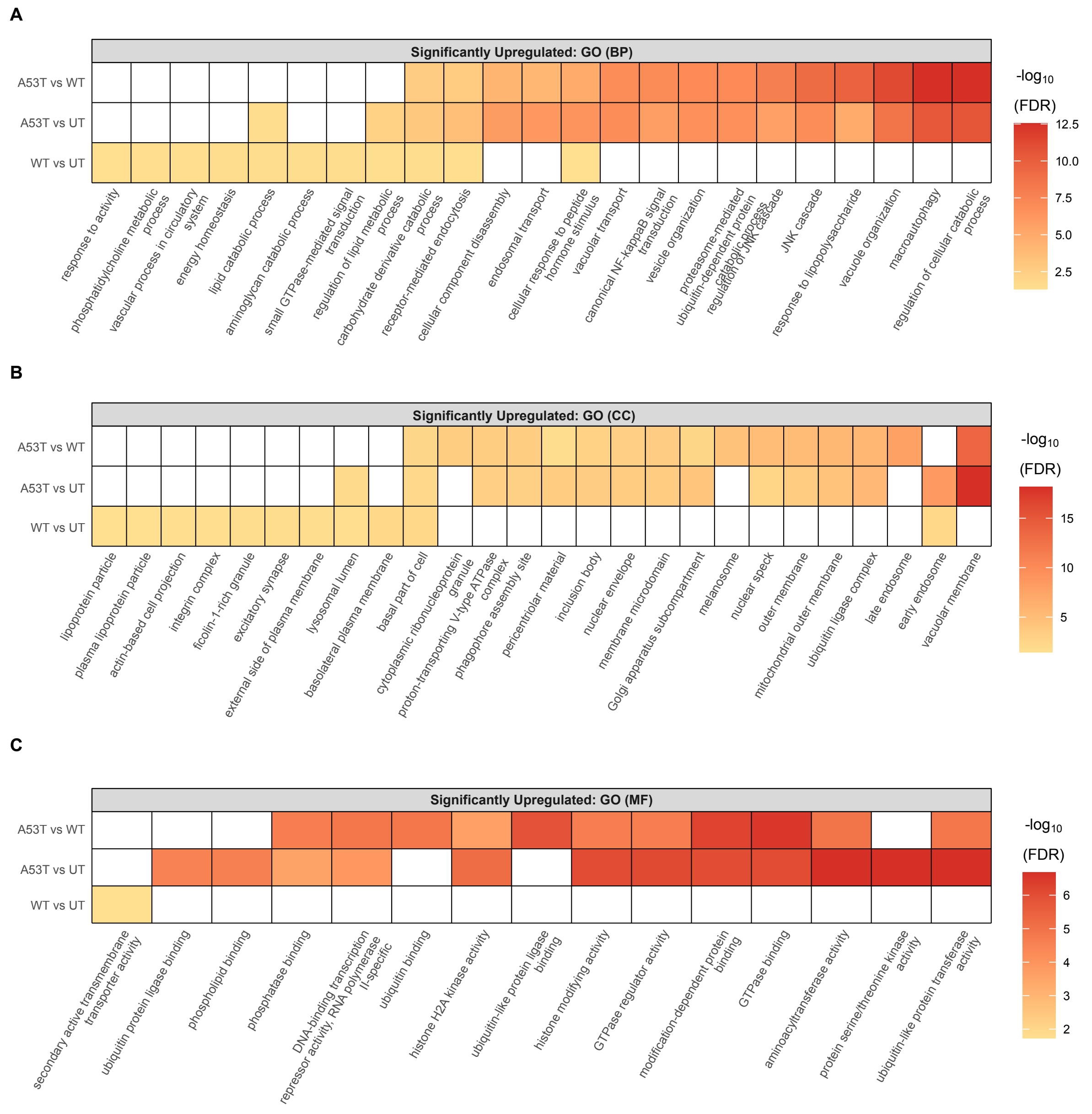


**Supplementary Figure 19. Upregulated Gene Ontology (GO) terms from bulk RNA-seq analysis of αSyn monomer treatment in iMGL.**

Bulk RNA-seq was performed on iMGL treated with wild-type (WT) or A53T αSyn monomers or untreated (vehicle). The top 10 significantly upregulated GO terms, ranked by FDR-corrected p value, are shown for each comparison: (A) Biological Process (BP), (B) Cellular Component (CC), and (C) Molecular Function (MF).

A

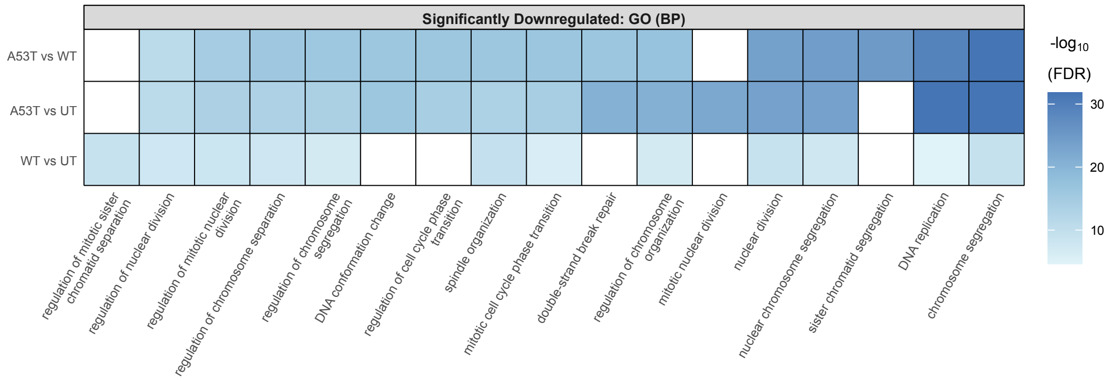


B

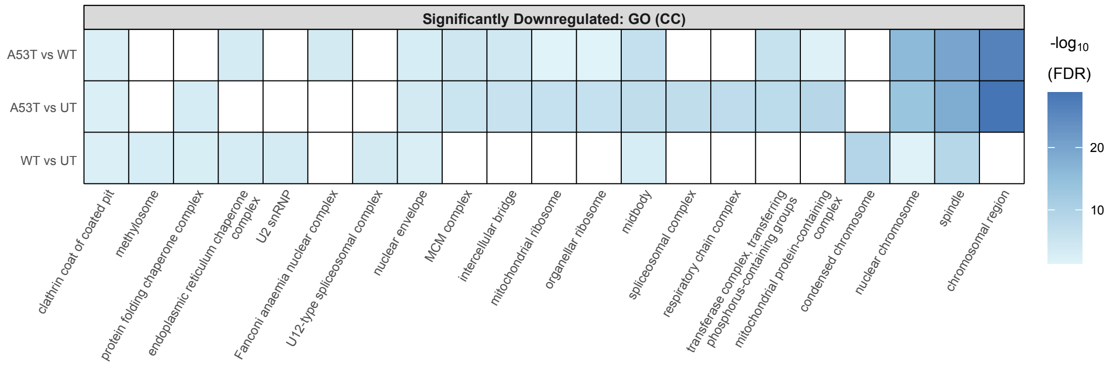


C

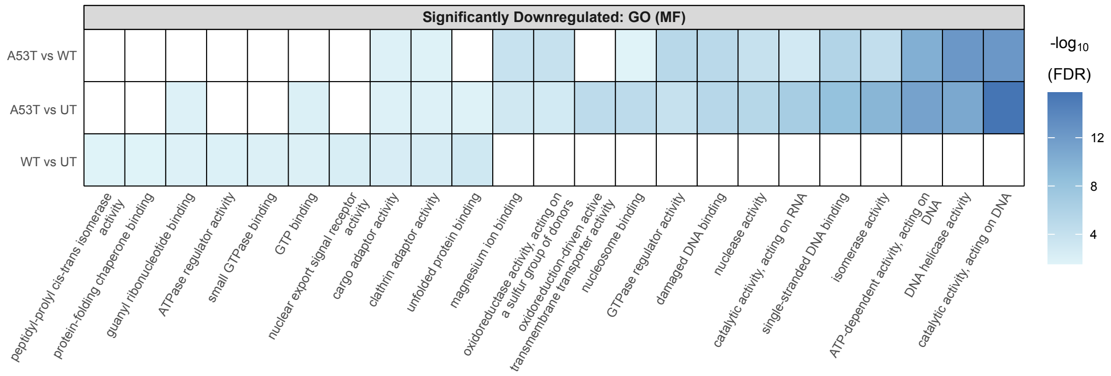


**Supplementary Figure 20. Downregulated Gene Ontology (GO) terms from bulk RNA-seq analysis of αSyn monomer treatment in iMGL.**

Bulk RNA-seq was performed on iMGL treated with wild-type (WT) or A53T αSyn monomers or untreated (vehicle). The top 10 significantly downregulated GO terms, ranked by FDR-corrected p value, are shown for each comparison: (A) Biological Process (BP), (B) Cellular Component (CC), and (C) Molecular Function (MF).

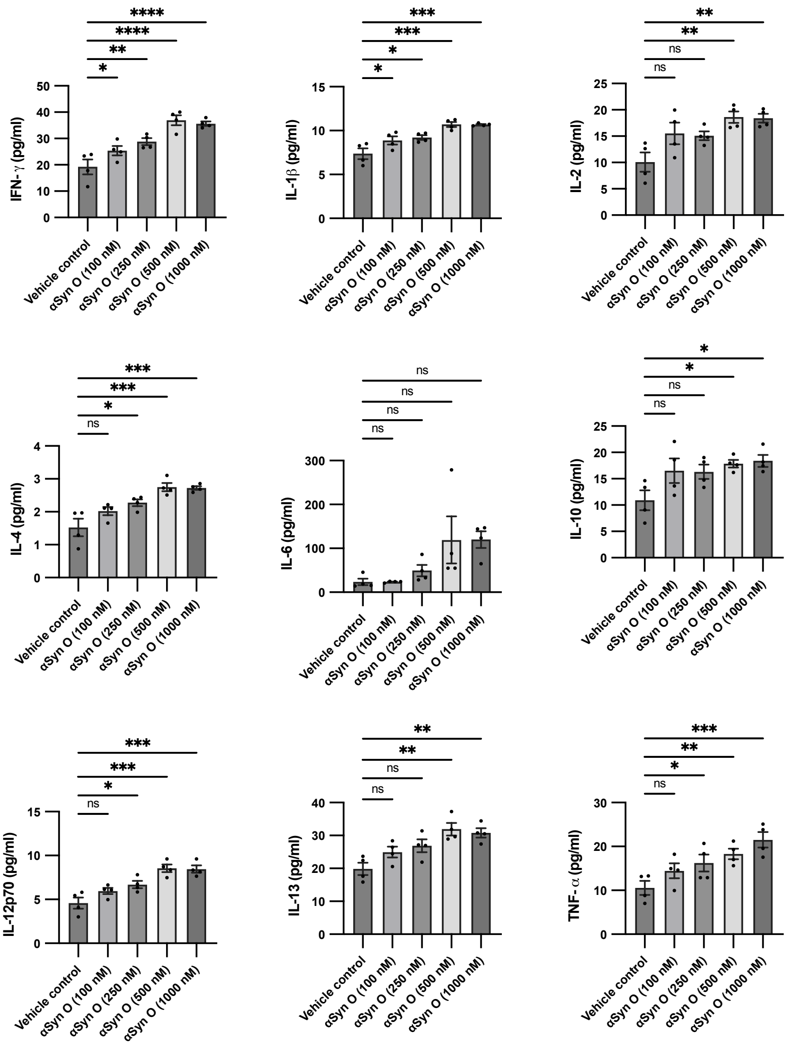


**Supplementary Figure 21. Dose-dependent induction of cytokine secretion in iMGL following αSyn oligomer treatment.**

Control iMGL were exposed to increasing concentrations of αSyn oligomers (100–1000 nM; StressMarq Biosciences; catalogue no. SPR-484). A panel of cytokines were measured by ELISA from conditioned media 24 hours after exposure (n = 4 iPSC donor lines; data represent mean ± SEM; repeated-measures one-way ANOVA with Dunnett's multiple comparison test comparing each concentration to the vehicle control; *$p < 0.05$, **$p < 0.01$, ***$p < 0.001$, ****$p < 0.0001$).

**Supplementary Table 1. Post-mortem cohort details**

| Case ID | Group | Sex | Onset | Duration (yrs) | PMI (h) | NPD | aSN | Tau | aB |
|---|---|---|---|---|---|---|---|---|---|
| PD1 | PD | F | 65 | 10 | 14 | LBDBS | 4 | 2 | NA |
| PD2 | PD | F | 77 | 9 | 22 | LBDBS | 4 | 2 | NA |
| PD3 | PD | M | 66 | 6 | 9 | LBDBS | 4 | 2 | NA |
| PD4 | PD | F | 71 | 11 | 16 | LBDBS | 3 | NA | NA |
| PD5 | PD | M | 70 | 15 | 16 | LBDBS | 4 | 2 | NA |
| PD6 | PD | M | 62 | 16 | 11 | LBDL | 4 | NA | NA |
| PD7 | PD | M | NA | NA | 16 | LBDBS | 3 | 1 | NA |
| PD8 | PD | F | 83 | NA | 5 | LBDBS | 3 | 3 | NA |
| PD9 | PD | F | NA | NA | 12 | LBDL | 4 | 3 | 5 |
| PD10 | PD | M | NA | NA | 22 | LBDL | 4 | 2 | NA |
| PD11 | PD | F | NA | NA | 8 | LBDL | 4 | 1 | 1 |
| PD12 | PD | M | NA | NA | 24 | LBDL | 4 | 2 | 3 |
| PD13 | PD | M | NA | NA | 13 | LBDBS | 3 | 2 | 2 |
| PD14 | PD | M | NA | NA | 24 | LBDBS | 3 | 1 | 3 |
| PD15 | PD | M | NA | NA | 17 | LBDL | 4 | 0 | 0 |
| PD16 | PD | M | NA | NA | 19 | LBDL | 4 | 1 | 1 |
| PD17 | PD | M | NA | NA | 6 | LBDBS | 4 | 2 | NA |
| PD18 | PD | M | 64 | 13 | 24 | LBDL | 4 | 1 | 0 |
| CON1 | Control | M | NA | NA | 29 | Control | NA | NA | NA |
| CON2 | Control | M | NA | NA | 8 | Control | NA | NA | NA |
| CON3 | Control | F | NA | NA | 24 | Control | NA | NA | NA |
| CON4 | Control | F | NA | NA | 12 | Control | NA | NA | NA |
| CON5 | Control | M | NA | NA | 12 | Control | NA | NA | NA |
| CON6 | Control | F | NA | NA | 21 | Control | NA | NA | NA |
| CON7 | Control | M | NA | NA | 26 | Control | NA | NA | NA |
| CON8 | Control | M | NA | NA | 31 | Control | NA | NA | NA |
| CON9 | Control | M | NA | NA | 24 | Control | NA | NA | NA |
| CON10 | Control | F | NA | NA | 22 | Control | NA | NA | NA |
| CON11 | Control | M | NA | NA | 23 | Control | NA | NA | NA |
| CON12 | Control | M | NA | NA | 17 | Control | NA | NA | NA |
| CON13 | Control | M | NA | NA | 30 | Control | NA | NA | NA |
| CON14 | Control | F | NA | NA | 23 | Control | NA | NA | NA |
| CON15 | Control | F | NA | NA | 13 | Control | NA | NA | NA |
| CON16 | Control | M | NA | NA | 18 | Control | NA | NA | NA |
| CON17 | Control | M | NA | NA | 48 | Control | NA | NA | NA |
| CON18 | Control | M | NA | NA | 12 | Control | NA | NA | NA |
| CON19 | Control | M | NA | NA | 21 | Control | NA | NA | NA |
| CON20 | Control | F | NA | NA | 20 | Control | NA | NA | NA |

**Supplementary Table 2. snRNAseq covariates**

| Sample_id | Batch | Downsampled | Merged | Redo | Sex | Total deduplicated percentage |
|---|---|---|---|---|---|---|
| CON12C | 1 | no | no | no | M | 63.0295252 |
| CON12F | 1 | no | no | no | M | 47.460259 |
| CON12P | 1 | no | no | no | M | 33.7817136 |
| CON13C | 6 | no | no | no | M | 48.7090577 |
| CON13F | 6 | no | no | no | M | 50.2297925 |
| CON13P | 6 | no | no | no | M | 56.2141767 |
| CON4C | 7 | no | no | no | F | 31.9986249 |
| CON9C | 16 | no | no | no | M | 50.8151681 |
| CON9F | 16 | yes | yes | no | M | 53.1835766 |
| CON9P | 16 | no | no | no | M | 63.443826 |
| CON11C | 10 | no | no | no | M | 58.5175134 |
| CON11F | 10 | no | no | no | M | 53.3195972 |
| CON11P | 10 | no | no | no | M | 57.9509041 |
| CON6C | 3 | no | no | no | F | 31.6323236 |
| CON6F | 3 | no | no | no | F | 49.3785169 |
| CON6P | 3 | no | no | no | F | 42.7226389 |
| CON7F | 17 | no | no | no | M | 60.6224419 |
| CON7P | 17 | no | no | no | M | 47.5449473 |
| CON1C | 8 | no | no | no | M | 44.732686 |
| CON1F | 8 | no | no | no | M | 32.3549518 |
| CON1P | 8 | no | no | no | M | 49.7077773 |
| CON10C | 12 | no | no | no | F | 53.0375431 |
| CON10F | 12 | no | no | no | F | 55.5104425 |
| CON10P | 12 | no | no | no | F | 64.6312644 |
| CON2F | 18 | no | no | no | M | 51.2596404 |
| CON2P | 18 | no | no | no | M | 47.9335444 |
| CON8C | 2 | no | no | no | M | 62.6864509 |
| CON8F | 2 | no | no | no | M | 54.3268192 |
| CON8P | 2 | no | no | no | M | 74.7299339 |
| PD1C | 6 | no | no | no | F | 41.8324678 |
| PD1F | 6 | no | no | no | F | 59.2197321 |
| PD1P | 6 | no | no | no | F | 50.3483204 |
| PD2C | 4 | no | no | no | F | 57.8819946 |
| PD2F | 4 | no | no | no | F | 58.7890805 |
| PD2P | 4 | no | no | no | F | 55.4175621 |
| PD3C | 12 | no | no | no | M | 47.6121641 |
| PD3F | 12 | no | no | no | M | 60.8644547 |
| PD3P | 12 | no | no | no | M | 61.9275557 |
| PD4C | 8 | no | no | no | F | 57.1668181 |

| | | | | | | |
|---|---|---|---|---|---|---|
| PD4P | 8 | no | no | no | F | 24.813816 |
| PD5C | 7 | no | no | no | M | 50.0798035 |
| PD5F | 7 | no | no | no | M | 38.5124574 |
| PD5P | 7 | no | no | no | M | 32.0669177 |
| PD6C | 2 | no | no | no | M | 64.5088451 |
| PD6F | 2 | no | no | no | M | 60.7189321 |
| PD6P | 2 | no | no | no | M | 60.8496189 |
| PD7F | 5 | no | no | no | M | 57.6198319 |
| PD7P | 5 | no | no | no | M | 26.4220735 |
| PD8C | 11 | no | no | no | F | 55.5219291 |
| PD8F | 11 | no | no | no | F | 51.3304649 |
| PD8P | 11 | no | no | no | F | 38.9381057 |
| CON3F | 18 | no | no | no | F | 57.3330375 |
| PD17F | 18 | no | no | no | M | 60.2251435 |
| PD9F | 15 | no | no | no | F | 7.7882303 |
| PD10C | 1 | yes | no | no | M | 54.4395075 |
| PD10F | 1 | no | no | no | M | 54.4128755 |
| PD10P | 1 | no | no | no | M | 61.001961 |
| PD11C | 9 | no | no | no | F | 63.702405 |
| PD11F | 9 | no | no | no | F | 47.2305358 |
| PD11P | 9 | no | no | no | F | 44.7010871 |
| PD12C | 10 | no | no | no | M | 61.0850901 |
| PD12F | 10 | no | no | no | M | 55.7814537 |
| PD12P | 10 | no | no | no | M | 56.1159596 |
| PD13C | 17 | no | no | no | M | 52.7656164 |
| PD13P | 17 | no | no | no | M | 59.6665313 |
| PD15P | 14 | no | no | no | M | 32.1341964 |
| PD16C | 13 | no | no | no | M | 51.9876366 |
| PD16F | 13 | no | no | no | M | 55.0206699 |
| PD16P | 13 | no | no | no | M | 54.4492993 |
| PD18C | 19 | no | no | no | M | 61.6791978 |
| PD18P | 19 | no | no | no | M | 62.0107876 |
| CON18F | 17 | no | no | no | M | 59.2895722 |
| CON18P | 17 | no | no | no | M | 58.1802813 |
| CON14C | 14 | no | no | no | F | 34.4360749 |
| CON19C | 19 | no | yes | partial | M | 35.0758348 |
| CON19F | 19 | no | no | no | M | 38.2895356 |
| CON19P | 19 | no | no | no | M | 40.874229 |
| CON17C | 5 | no | no | no | M | 37.2747627 |
| CON17F | 5 | no | no | no | M | 52.851149 |
| CON5C | 9 | no | no | no | M | 51.7071122 |
| CON5F | 9 | no | no | no | M | 41.1843152 |

| | | | | | | |
|---|---|---|---|---|---|---|
| CON5P | 9 | no | no | no | M | 58.1582315 |
| CON15C | 4 | no | no | no | F | 49.3556546 |
| CON15F | 4 | no | no | no | F | 54.121887 |
| CON15P | 4 | no | no | no | F | 48.2614804 |
| CON16C | 11 | no | no | no | M | 60.009812 |
| CON16F | 11 | no | no | no | M | 60.7097847 |
| CON16P | 11 | no | no | no | M | 56.3116219 |
| CON20C | 13 | no | no | no | F | 51.4450558 |
| CON20F | 13 | no | no | no | F | 46.8279334 |
| CON20P | 13 | no | no | no | F | 34.775863 |
| PD17C | 18 | no | no | no | M | 51.2403963 |
| CON3P | 18 | no | no | no | F | 56.8371608 |

**Supplementary Table 3. hiPSC line details**

| **hiPSC line** | **Mutation** | **Sex of donor** | **Source** |
|---|---|---|---|
| Control 1 | None | Female | Thermo Fisher Scientific A18945 |
| Control 2 | None | Male | The Jackson Laboratory (KOLF2.1J) |
| Control 3 (isogenic for A53T 3) | *SNCA* A53T corrected | Female | Applied StemCell (project ID: C1729) |
| Control 4 | None | Male | EBiSC: WTSIi019-B |
| Control 5 | None | Female | PPMI |
| Control 6 | None | Female | PPMI |
| Control 7 (isogenic for A53T 4) | *SNCA* A53T corrected | Female | PPMI |
| Control 8 (isogenic for A53T 5) | *SNCA* A53T corrected | Female | PPMI |
| A53T 1 | *SNCA* A53T | Male | StemBANCC: STBCi023-C |
| A53T 2 | *SNCA* A53T | Female | PPMI |
| A53T 3 | *SNCA* A53T | Female | StemBANCC: STBCi019-C |
| A53T 4 | *SNCA* A53T | Female | PPMI |
| A53T 5 | *SNCA* A53T | Female | PPMI |
